# Improved ancestral genome reconstruction using a learned gene-content grammar

**DOI:** 10.64898/2026.09.03.749268

**Authors:** Gergely J. Szöllősi, Anja Spang, Bastien Boussau, Tom A. Williams

**Affiliations:** Model-Based Evolutionary Genomics Unit, Okinawa Institute of Science and Technology Graduate University, Okinawa, Japan; Institute of Evolution, HUN-REN Center for Ecological Research, H-1121 Budapest, Hungary; Department of Marine Microbiology and Biogeochemistry, NIOZ, Royal Netherlands Institute for Sea Research, 1790 AB Den Burg, The Netherlands; Department of Evolutionary & Population Biology, Institute for Biodiversity and Ecosystem Dynamics (IBED), University of Amsterdam, 1090 GE Amsterdam, The Netherlands; Laboratoire de Biométrie et Biologie Evolutive, Univ Lyon, Univ Lyon 1, CNRS, VetAgro Sup, Villeurbanne, France; Centre for Evolution, Department of Life Sciences, University of Bath, BA2 7AX, United Kingdom

**Keywords:** ancestral genome reconstruction, gene content, phylogenetic reconciliation, Ising model, gene co-occurrence, LBCA, LACA

## Abstract

Ancestral gene content inferences allow inferring the set of genes - and by extension, the cellular features and metabolic capabilities - of ancestral organisms, based on data from modern genomes. Current methods differ in their approach to ancestral inferences and the kinds of errors they make: reconciliation methods map gene trees onto species trees, and tend to underestimate ancestral contents due to phylogenetic noise; profile methods model the evolution of phylogenetic profiles (presence-absence or count data) on the species tree, and tend to return inflated ancestors because they ignore gene trees and as a result can only account for horizontal gene transfer (HGT) in a limited manner. For reasons of tractability, both approaches also share a core limitation: neither uses the fact that genes do not act alone but belong to operons, protein complexes, and metabolic pathways that may be gained and lost together or experience shared selective constraints. Here, we show that this context - the grammar of gene content - provides a rich source of information that can be used to greatly improve ancestral gene content inference and metabolic reconstruction under both the reconciliation-and profile-based approaches. We model this structure as an Ising model and infer its parameters from 113,104 bacterial and archaeal genomes (one per species representative in GTDB). We validate the model on extant taxa using phylum-level holdout (i.e. using test data from different prokaryotic phyla than training data), showing that it can accurately “denoise”, i.e., reconstruct gene repertoires from highly fragmented and noisy input data, learning about protein-protein interactions and gene essentiality during the training process. When applied to ancestral reconstructions, the denoiser fills gaps in conservative reconstructions and removes excess genes from overly-generous ones, such that different reconstruction methods converge to broadly concordant conclusions. By using this gene content grammar, patchy method-dependent ancestral reconstructions can be turned into organism-like ones, and yield agreement on the gene families, cell-biological features, and metabolic capabilities of the deepest nodes in the tree of life.

## Introduction

The geological record of ancient microbial life is sparse and much of our understanding of early evolution derives from phylogenomic analyses of modern genomes. One approach is ancestral gene content reconstruction, in which the gene family repertoires of ancestral nodes in the tree of life can be inferred based on extant genomes as a basis to infer characteristics of these ancestors, including metabolic potential and cell biological traits. Functional gene family annotations are ultimately derived from experiments on model organisms and thus assume that function has been conserved across evolutionary time. For example, inferences about the nature of the last universal common ancestor (LUCA) (Weiss et al., 2016; Moody et al., 2024), and the common ancestors of the archaeal (Williams et al., 2017; Huang et al., 2026) and bacterial (Coleman et al., 2021) domains, are based on ancestral gene content reconstruction using reconciliation approaches.

Two kinds of methods are widely used to infer ancestral gene contents: gene tree-species tree (or phylogenetic) reconciliation (Williams et al., 2024), compare gene family trees with a rooted species tree, with the differences between them reconciled using a model of gene duplication, transfer and loss (Szöllősi et al., 2012, 2013; Morel et al., 2020, 2024); and profile-based methods, in which the presence-absence or copy number patterns of gene families, but not their gene trees, are modelled as evolving along the rooted species tree using a birth-death process (De Bie et al., 2006; Csűrös, 2010). Both provide reasonable frameworks for ancestral reconstruction, but their different properties lead to different kinds of systematic errors (Williams et al., 2024; Csűrös, 2026). Reconciliation methods tend to under-estimate ancestral gene contents because they interpret errors in reconstructed gene trees as spurious gene transfers, biasing the reconstruction in favour of scenarios in which even widely-distributed gene families may originate relatively recently in evolution and are spread by transfer (Williams et al., 2024; Csűrös, 2026). Although this bias is lessened by more complex and better-fitting reconciliation models (for example, by relaxing the assumption that rates of gene origination, duplication, transfer and loss are constant across the tree (Coleman et al., 2021; Huang et al., 2026)), reconciliation models nonetheless tend to recover smaller ancestors than profile-based methods. By contrast, profile-based methods (such as the gain-loss-duplication (GLD) model implemented in Count (Csűrös, 2010) and Recount (Szöllősi and Williams, 2026) have an opposite bias. As they do not consider gene tree information, they cannot distinguish an ancient, vertically-inherited gene from a widespread but recently-transferred one. As a result, GLD over-attributes broadly-distributed gene families to ancestors, returning large ancestral genomes that tend to be outside the range of extant gene family complements for a group (Williams et al., 2024). While reconciliation-based reconstructions are almost certainly conservative compared to profile-based ones (in extreme cases, recovering almost no gene families confidently at the root of large species trees, (Davín et al., 2025), the practical question of how to interpret the results of both methods in order to draw inferences about character or organismal evolution remains difficult to resolve.

In addition to these challenges, both approaches share a significant common limitation: although some parameter values may be shared across families, the history of each gene family is reconstructed in isolation, without reference to the reconstructed histories of other gene families. However, genes do not act alone: they interact with other genes to carry out metabolic reactions within interconnected pathways, form protein complexes, take part in regulatory networks, and ultimately give rise to organismal function. As a result, functionally-coupled genes co-occur on genomes, while incompatible genes are unlikely to be encoded on the same genome (Beavan et al. PNAS). This contextual structure, which might be viewed as a statistical “grammar” of gene content, is a rich source of information about the composition of gene repertoires that is not currently used by either approach to ancestral reconstruction. Here, we show that this grammar can be used both to judge the plausibility of, and significantly improve, gene content reconstructions from both reconciliation and profile methods.

We capture this information about gene family co-occurrence and co-avoidance using a pairwise Ising model (Nguyen et al., 2017) that assigns each presence-absence pattern a probability: high when it is compatible with the learned patterns in extant genomes, and low when it is not. We fit the Ising model to 113,104 prokaryotic genomes, one for each species in r220 of the Genome Taxonomy Database (GTDB; Parks et al., 2022). We then treat the improvement (or denoising) of a gene content estimate as inference under the Ising model, shifting the profile towards a high-probability configuration by the same iterative machinery that powers some modern denoising generative models (Ho et al., 2020) but over our explicit, interpretable scoring function. To validate the genome denoiser, we use whole-phylum hold-outs: the model is trained on genomes from one set of prokaryotic phyla, and tested on genomes from unseen phyla, representing *>* 1 Ga evolutionary divergence in each case. We show that the trained model recovers most of the gene content of modern archaea and bacteria from highly partial and “contaminated” inputs – 82% of the true genes at moderate loss and 68% under deep-ancestral corruption, at 80–86% precision – while remaining statistically well-calibrated. Interestingly, the trained models can also predict experimentally-validated protein-protein interactions and gene essentiality, consistent with previous work suggesting that co-occurrence data are also informative about these aspects of biology.

When applied to estimated ancestral gene contents, the denoiser fills in missing genes in patchy reconciliation-based inputs and prunes unlikely combinations of genes from inflated profile-based reconstructions. As a result, previous reconciliation-and GLD-based reconstructions of the last bacterial common ancestor (LBCA) and last archaeal common ancestor (LACA), which disagree moderately to strongly as inputs, converge to concordant reconstructions that fall within the gene content size ranges of modern prokaryotes after denoising. The denoised LBCA was predicted to bea free-living diderm, while the denoised LACA was reconstructed to be a methanogen in line with previous inferences. Overall, this work shows that Ising-based denoising is a powerful tool for improving ancestral genome reconstruction that can resolve differences among existing reconstruction methods, and may help in the task of drawing biological conclusions about ancient life forms from ancestral reconstructions.

## Results

### A learned grammar of gene content

We describe each genome as the set of homologous gene families it encodes (out of a set of *N* = 4,789 COGs, Clusters of Orthologous Genes (Galperin et al., 2021)), so a genome is a length-*N* binary vector ***x*** *∈ {−*1, +1*}^N^*, with *x_i_* = +1 for presence of a gene family. We model the distribution of gene content as a pairwise Ising (Boltzmann) distribution,

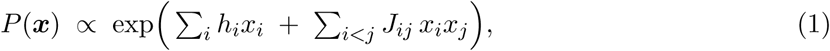

with learned fields ***h*** *∈* R*^N^*(how common each family is on its own; 4, 789 parameters) and a symmetric, hollow coupling matrix ***J*** whose entry *J_ij_* says whether families *i* and *j* tend to occur together (*J_ij_ >* 0) or to exclude one another (*J_ij_ <* 0) (*C*^2^ = 11,464,866 parameters). *P* (*x*) scores a genome by how well it respects the co-occurrence structure of the training data, so gene combinations observed on sampled extant genomes have high-probability. Fitting one joint model, rather than scoring gene pairs separately, lets ***J*** distinguish families that are directly coupled from those that co-occur through shared partners. Joint pairwise models have been fit to gene presence/absence before, to predict functional and physical interaction (Croce et al., 2019; Fukunaga and Iwasaki, 2022), generalising the phylogenetic profiles of Pellegrini et al. (1999); we instead use the fitted model generatively, as a prior under which a corrupted genome is denoised. By “corruption”, we mean a gene content profile that contains errors, either deliberately introduced during training and validation, or as the result of error-prone and noisy ancestral reconstruction algorithms. The task of “denoising” a corrupted profile is then framed as the search for the high-probability configuration consistent with it. Starting from the noisy observation ***x***^0^, each gene presence indicator *x_i_* is repeatedly re-estimated from the current guess of its neighbours, using a mean-field approximation

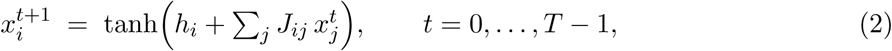

Repeated steps of *x_i_*re-estimation drive the configuration toward a self-consistent, high-probability genome. The number of steps *T* is a relaxation depth that sets how far the configuration may move from the input data, and is learned during model training. The model includes four additional components that adapt the Ising model to the gene content denoising problem: a Thouless-Anderson-Palmer (TAP/Onsager) reaction term that removes the self-reinforcement bias of naive mean field (Thouless et al., 1977); learned, step-dependent gates that decide, for each gene, how much to trust the raw observation versus the couplings; an adaptive temperature that sharpens the update with genome richness; and module-completeness conditioning that helps the model reconstruct complete pathways (Materials and Methods; Supplementary Material, Sec. S1). In particular, the completeness module affects at each step *t* how much the input data *x*_i_^0^ is used, and how much coupling should affect the *x_i_^t^* value.

The Ising model captures pairwise interactions between genes, but it seems reasonable to suppose that accounting for higher-order interactions, for example among sets of interacting genes, might further improve performance. To investigate this, we explored two extensions to the model to capture higher-order structure: marginalising a vector of hidden units coupled to the visible genes, and a learned attention-parameterised third-order correction head.

In all three cases, we trained the model on the 113,104 representative species sampled from the GTDB under noise introduced through two channels: a false-negative channel that deletes truly present genes, and a false-positive channel that inserts spurious genes as contaminants.

All models were trained using a staged curriculum, where tasks of increasing difficulty are used to train a part of the model or the model in its entirety (Materials and Methods; Supplementary Material, Sec. S1.4). In the first stage, only the parameters of the Ising model are trained, then step-dependent gates, temperatures, and parameters of the module completeness conditioning get optimized, as levels of noise (false negatives in particular) are progressively increased. At the last stage, parameters of the denoiser were optimized for a denoising process with *T* = 20 steps, using AdamW to estimate, with parameters of the Ising model frozen.

We additionally fine-tuned models for each prokaryotic domain. To develop models that are able to deal with both overly-conservative and overly-liberal ancestral gene content reconstructions, we explored settings in which both the false negative rate was high (e.g., 80-90% of genes deleted, which may be comparable to the extreme patchiness of reconciliation-based ancestral gene content reconstructions under certain conditions), and in which false-positive contaminants were preferentially drawn from the most widespread gene families among prokaryotes. This is because we expect genes that are widespread at the tips of the tree to be among the most likely to be spuriously mapped to the root in profile-based reconstructions, and also because these families are harder to detect as false-positives based on co-occurrence information (see Materials and Methods).

### The denoiser recovers held-out genomes across billion-year divergences

To determine the performance of the Ising denoisers, we first investigated their ability to reconstruct extant genomes (for which the ground truth is known) given corrupted - that is, partial and noisy - inputs. For training, we simulated these corrupted genomes by sweeping the false-negative rate *f*_N_ from 0 to 0.9, with a (low) false-positive rate of 0.01; this is intended to replicate the situation for deep nodes on the tree of life, where we expect most errors in reconciliation-based gene content reconstructions to be false-negatives, due to the under-estimation of the age of origination described above.

Since related genomes have similar gene content, a model for which the training data included close relatives of the test genome might perform well merely by taking advantage of this phylogenetic inertia, rather than by learning more generalisable aspects of gene content grammar. To guard against this possibility and help the models generalise, we trained several models with a phylogenetically-aware hold-out strategy, in which individual prokaryotic phyla were allocated either to the training or test datasets. Given the ages of prokaryotic phyla, this strategy means that training and test data are separated by at least 1 Gyr, and more typically 2-2.5 Gyr, of evolutionary divergence (Davín et al., 2025).

We evaluated each trained model on two distinct metrics. *Recovery* is how accurately gene content is reconstructed, summarized by the Matthews correlation coefficient (MCC), which is the Pearson correlation coefficient between the true and inferred presence-absence profile. *Calibration* measures the accuracy of the presence probabilities assigned to each gene family; for example, whether a gene assigned a presence probability of 0.8 is actually present about 80% of the time, summarized by the expected calibration error (ECE, the average difference between the predicted probability and observed frequency). When trained in this way, the denoiser recovers held-out genomes across the noise spectrum (Fig. 2). The best model recovers far more accurate genomes than the raw input (MCC 0.657 versus 0.22 at *f*_N_ = 0.90, recovering roughly three-fifths of the presence/absence signal after 90% of the true genes are deleted). Interestingly, the denoiser also improves calibration (that is, the accuracy of the advertised presence probabilities) compared to the raw reconciliation-based reconstruction (ECE *∼*0.04 versus *∼*0.24).

**Figure 1:**
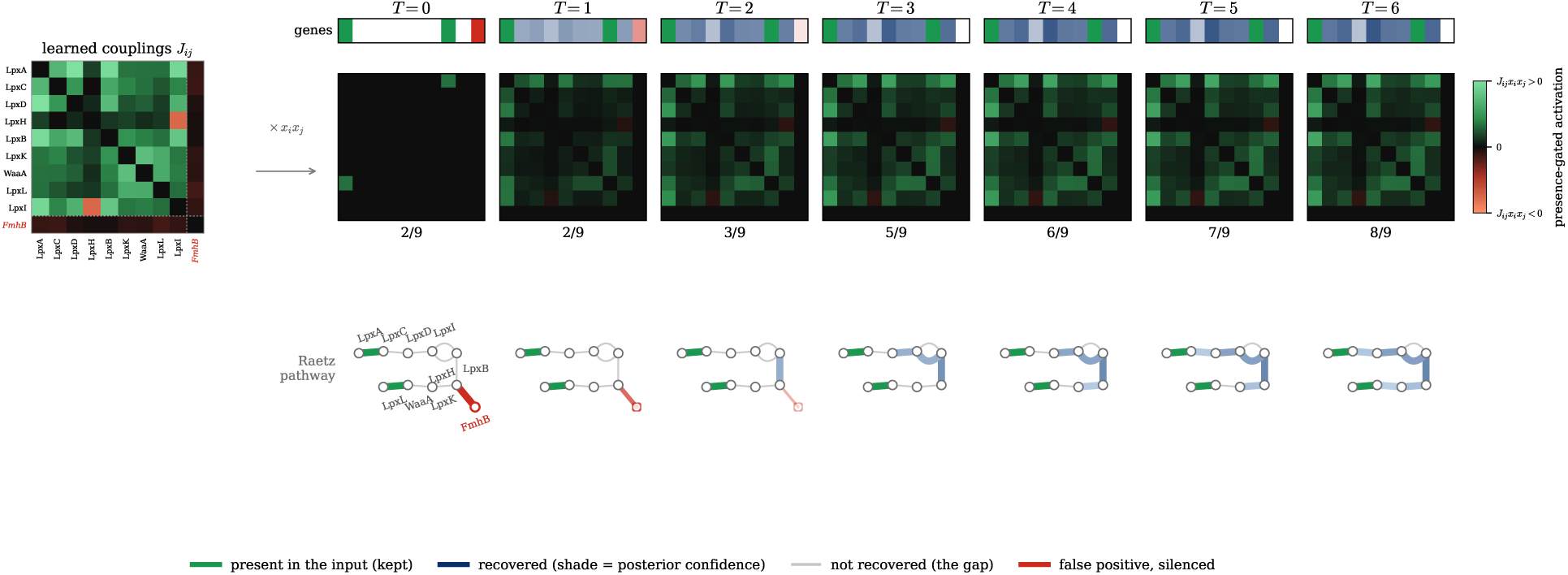
The denoiser iteratively completes a metabolic module, as illustrated by a worked example on the Raetz lipid A pathway (KEGG M00060, the KDO_2_-lipid A module of the diderm envelope). The input is partial and noisy, with two of the module’s nine gene families present, plus one false positive gene (FmhB) added. **(Left)** the module’s learned coupling sub-block *J_ij_*itself, with the genes labelled - nine Raetz enzymes plus the false positive FmhB in **red**); the heatmap sequence to its right is this same matrix gated by gene presence, *J_ij_x_i_x_j_*, at each iteration. **(Top)** the module gene vector at each step: **green** families were present in the input (kept), **blue** families are recovered by the model (shade = posterior confidence), and the **red** family is the false positive; the present count climbs from 2 to 8 of 9 by *T* =6. **(Middle)** the presence-gated activation *J_ij_x_i_x_j_* of the module’s learned coupling sub-block — **green** where a co-present pair lowers the energy (*J_ij_x_i_x_j_ >* 0), **red** where it raises it, and black where a family is still absent and its couplings are gated off. **(Bottom)** the same reconstruction on the pathway itself, with gene families as edges, coloured as above with opacity = posterior confidence and grey for the families still not recovered (the remaining gap). The module is filled back in from its own internal co-occurrence couplings, illustrating how the denoiser operates genome-wide.

**Figure 2:**
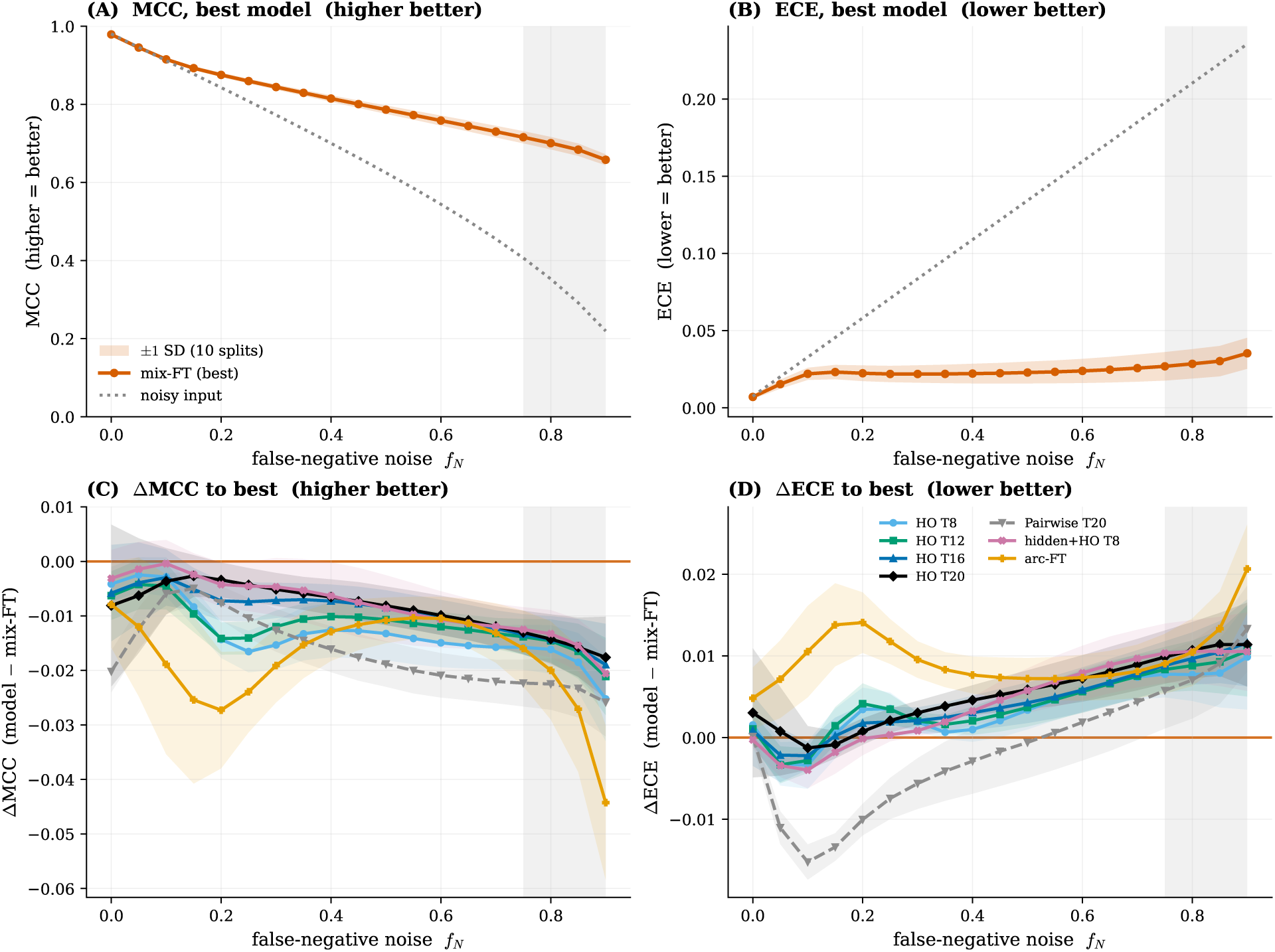
Recovery and calibration across the noise spectrum, under whole-phylum hold-out. (A,B) The best model (mix-FT, *T* =20) against the noisy input, in MCC (A, higher is better) and ECE (B, lower is better) versus the false-negative rate *f*_N_ at *f*_P_ = 0.01: the denoiser recovers far more signal than the raw input and stays calibrated where the raw input does not. (C,D) The difference from mix-FT for the other model families at matched depth; all trail it in recovery, least so the deeper higher-order models, while in calibration the plain pairwise model is *better* than mix-FT (lower is better) at low-to-moderate noise and only high levels of noise (typical of deep-ancestral reconstructions, grey band) favour mix-FT. Bands are *±*1 s.d. across the 10 whole-phylum cross-validation splits (8 for arc-FT, which exists for only 8 of the 10 folds).

The 10-way folds hold out whole phyla, so each test genome comes from a lineage absent from the pretraining set: the generalist root never sees the evaluation phyla. The fine-tuned specialists re-split their data independently of that root, however, so a genome held out of the pretraining may still have had its phylum in the fine-tune training set; this applies to almost all (99%) of the genomes on which mix-FT is scored. We find no evidence that this inflates recovery: on the leak-free remainder MCC is 0.016–0.025 *higher* than on the validation set as a whole (for example 0.680 versus 0.658 at *f*_N_ = 0.9), although that subset is small (*n* = 349 in the stratified comparison). For the per-genome illustrations below we require the genome to be held out at both training stages.

Of the three model types considered, the pairwise Ising model already shows strong performance, and in fact is better calibrated than the more complex models that consider higher-order interactions (ECE 0.006–0.007 at *f*_N_ = 0.10, roughly a third of the higher-order models). The third-order head gives a small but consistent improvement in recovery, particularly for noisier genomes (Fig. 2C,D). The third approach, using explicit hidden units, does not provide a material improvement. We therefore use the simpler pairwise model with a learned third-order head as the focal approach in what follows. Since the training set is overwhelmingly bacterial, we additionally fine-tuned this model for each domain: a mixed-domain rebalance used for the LACA (mix-FT), and a bacteria-only specialist used for the LBCA (bac-FT). Both give a clearer improvement in calibration over the generalist, and mix-FT a small further gain in recovery (mix-FT on mixed hold-outs at *f*_N_ = 0.5: MCC 0.786 vs. 0.778, ECE 0.023 vs. 0.029; bac-FT is evaluated on bacteria-only hold-outs, Supplementary Material, Sec. S2), so we use the node-appropriate fine-tune for each ancestor. The ancestral reconstructions reported below are made with the contamination-trained descendants of these two models (bac-FT-fp-marginal-HQ for the LBCA, mix-FT-fp-marginal-cons for the LACA), which reject the coherent false positives that survive reconciliation (Supplementary Material, Sec. S3.1). Overall, this model performs well as a gene content denoiser: across all 105 held-out phyla with at least 30 genomes MCC is uniformly high (Fig. 2), archaea can be recovered as well as bacteria, and even fragmented genome assemblies (e.g. MAGs below 80% CheckM completeness) recover at MCC 0.59 at *f*_N_ = 0.90 (Supplementary Material, Sec. S2).

A potential concern with this approach might be that the denoiser is not inferring missing genes based on gene-gene co-ocurrance, but simply filling in gaps in the input with widespread genes (that is, regressing towards an average genome from the starting input). With a highly incomplete input, such a strategy might perform better than chance. To determine the added value of interactions (captured in J), we therefore re-evaluated the pairwise model (*T* = 20) with all *J_ij_* set to zero at inference. This performed substantially worse than the full denoiser, and increasingly so as the input degrades: mean MCC falls from 0.768 to 0.624 at *f*_N_ = 0.5 and from 0.632 to 0.230 at *f*_N_ = 0.9 (ten whole-phylum splits, *f*_P_ = 0.01). At *f*_N_ = 0.9 the ablated model barely improves on the noisy input it is given (MCC 0.22): stripped of the couplings, the denoiser returns roughly what it started with. At the noise levels relevant to ancestral reconstruction the couplings therefore carry almost all of the denoising signal, which cannot be explained by regression towards an average genome.

As a simple illustration of model performance, we applied the models to recover the *E. coli* genome based on corrupted input data. In training, Pseudomonadota (formerly named Proteobacteria, the phylum containing *E. coli*) was left out, so the model cannot “cheat” by simply memorizing the gene content of close *E. coli* relatives. Even with extensive corruption (false-negative rates 0.4*/*0.6*/*0.8; Fig. 3 with *f*_N_ = 0.4 and 0.6 in Supplementary Material, Sec. S3)), the genome can be largely rebuilt (recall 77 *±* 1*/*70 *±* 1*/*62 *±* 2%, mean *±* s.d. over 500 corruption draws).

**Figure 3:**
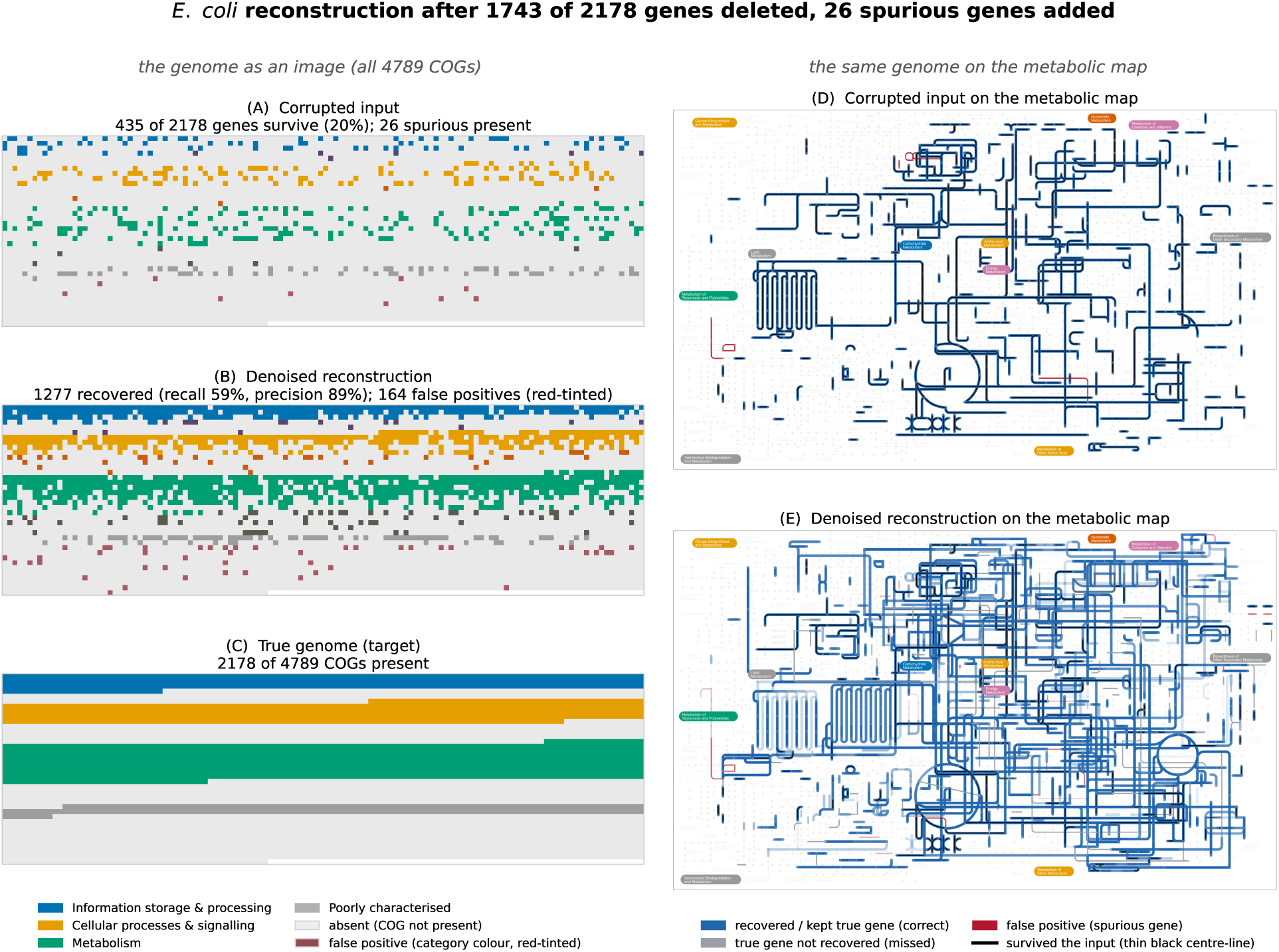
Ground-truth recovery of *E. coli* from corrupted input at a high false-negative rate (*f*_N_ = 0.8: 1,743 **of its** 2,178 **genes deleted,** 26 common-gene false positives added), with its entire phylum held out during training. *Left* (**A**–**C**), the genome as an image – each tile a COG family, grouped by functional category: the corrupted input (**A**), the denoised reconstruction (**B**), and the truth (**C**). Only 20% of the genome survives the corruption (**A**, mostly blank), yet the contamination-trained model (marginal-HQ, split 5) restores it to 59% recall at 89% precision. *Right* (**D**, **E**), the corrupted *input* (**D**) above the denoised *reconstruction* (**E**) on the iPath3 global metabolic map (Darzi et al., 2018), in the same scheme as the ancestral maps (Figs. 6, 7) but in E. coli’s own three correctness states: **blue** = a true gene the denoiser keeps or recovers (correct), **grey** = a true gene it does *not* recover (missing), **red** = a spurious gene it adds (false positive); grey and red are drawn thin so the correct reconstruction stands out. Input genes carry a thin black centre-line on both maps, so the sparse scaffold reads identically; on the reconstruction the genes the denoiser *recovered* are pure blue (no centre-line), and each present gene’s opacity is its posterior confidence (the input is solid, a binary observation). The skeleton fills back into a near-connected network, the thin red false positives making the cost honest – 164 common genes the prior adds that the true genome lacks. The same reconstruction at the milder *f*_N_ = 0.4 and 0.6 is in Supplementary Material, Sec. S3.

### Reconstruction quality scales with evolutionary distance to the training set

While the above results suggest that the denoiser can reconstruct partial inputs even when the training data is very distant to the target, it is also interesting to consider how reconstruction quality improves with access to more closely related training data. To investigate this, we re-ran the analysis on a nested series of training sets, each holding out a progressively smaller clade around *E. coli*: its phylum (Pseudomonadota), class (Gammaproteobacteria), an intermediate Gammaproteobacterial sub-clade, order (Enterobacterales), family (Enterobacteriaceae), and finally the single *E. coli* genome alone, leaving other *Escherichia* species in training. As might be expected, recovery rises monotonically with increasingly more closely related taxa in the trainin set (Fig. 4). At the false-negative and false-positive rates roughly comparable to deep-time inference (*f*_N_ = 0.8), the median Matthews correlation increases from 0.57 (recall 61%), when the closest available relative diverged *>* 2.5 Ga ago, to 0.77 (recall 82%) once same-genus relatives are present. The greatest improvement is confined to recent distances: performance is similar across a broad range of greater evolutionary distances, recovery plateauing near the 0.57 floor from *∼*1 to *∼*5.5 Ga, so it is proximity within the last few hundred million years, not deeper relatedness, that enables the model to improve accuracy. This suggests that, beyond “cheating” with the help of phylogenetic inertia on close timescales, performance is based on generalizable features of gene content grammar.

**Figure 4:**
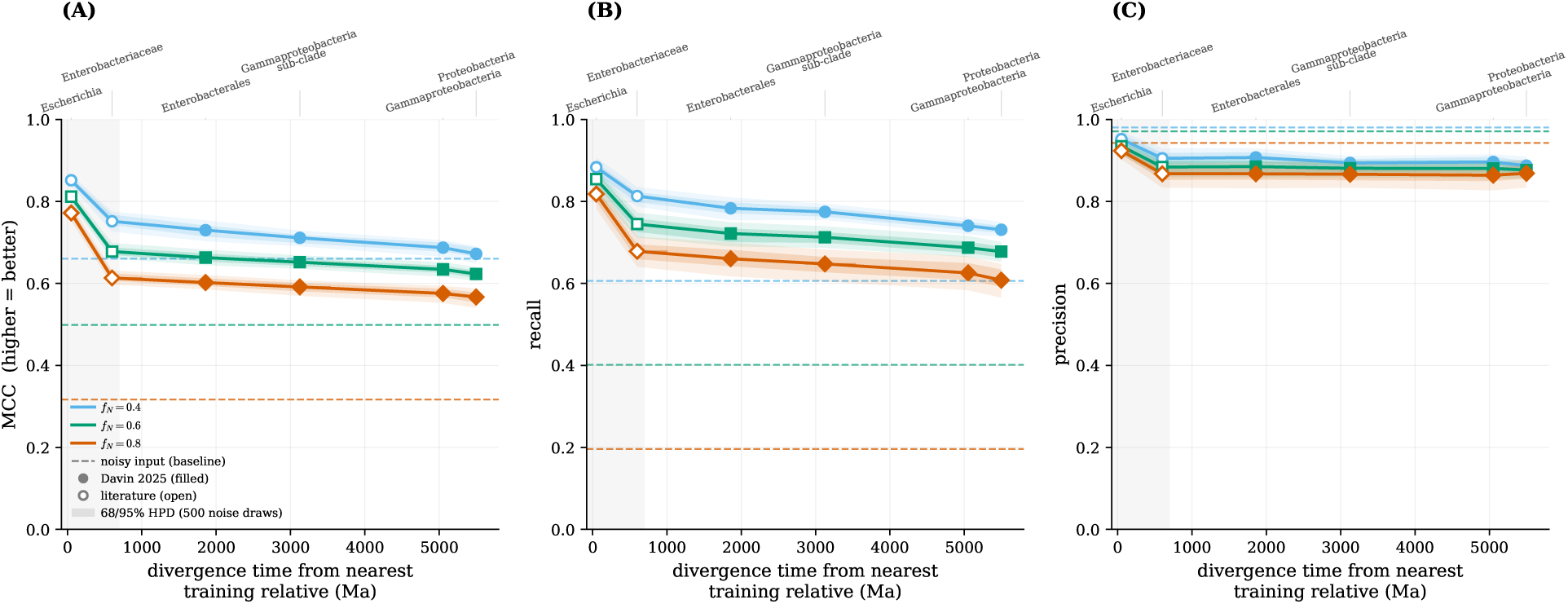
Ground-truth recovery of *E. coli* improves as the training set approaches it phylogenetically. The contamination-trained model (marginal-HQ) was fine-tuned on a nested series of holdouts around the Fig. 3 genome; the *x*-axis is the divergence time from the nearest training relative – summed over both lineages, i.e. twice the age of their most recent common ancestor – linear, in millions of years (Ma). **(A)** Matthews correlation, **(B)** recall, **(C)** precision, each at false-negative rates *f*_N_ = 0.4*/*0.6*/*0.8; dashed lines are the model-free noisy-input baseline at each *f*_N_. Markers are the median over 500 independent corruption draws at that *f*_N_, and the shaded bands behind each curve are the nested 95%*/*68% highest-density intervals of the metric over the same draws – the sampling spread is widest for recall at *f*_N_ = 0.8 (it is set by *which* 80% of genes the false-negative channel removes) and negligible for precision. The ages of the four deep nodes (Pseudomonadota, Gammaproteobacteria, the intermediate Gammaproteobacterial sub-clade, Enterobacterales; *filled* markers) are taken from the time-calibrated tree of Davín et al. (2025) (on which *E. coli* K-12 is the sole Enterobacteriaceae tip); the two shallow nodes (Enterobacteriaceae, *Escherichia*; *open* markers, shaded region) are molecular-clock estimates (Ochman et al., 1999; Walk et al., 2009), drawn with their published ranges as horizontal bars. Reading outward from the present, recovery drops sharply over the first few hundred million years and then *plateaus*: from the order rung outward – nearest relative *∼*1.9 to *∼*5.5 Ga diverged – it stays flat at MCC *≈*0.57–0.60 (recall *≈*61–66% at *f*_N_ = 0.8). Phylogenetic proximity to the training set therefore helps only at recent, sub-billion-year distances (the genus- and family-level rungs); beyond *∼*1 Ga the contamination-trained prior sets a floor that is insensitive to how deeply the nearest relative diverged. Divergence times are from the time-calibrated tree of Davín et al. (2025) for the deeper nodes and from other published molecular-clock analyses for the shallow ones (Ochman et al., 1999; Walk et al., 2009).

### Denoiser couplings encode physical interactions and gene essentiality

All else being equal, co-occuring genes are expected to be more likely than chance to function together and to potentially be involved in physical interactions (Pellegrini et al., 1999). To investigate whether the denoiser models capture this biological signal, we asked whether the couplings *J_ij_*learned on the complete set of genomes correspond to physical interactions between the proteins their families encode.

We compared *J* against experimentally validated protein-protein interactions in STRING v12.0 (Szklarczyk et al., 2023) for 19 bacterial pathogens from a recent deep-learning interactome screen (Humphreys et al., 2024), taking as ground truth STRING’s experimental channel. The coupling recovers interactions well above chance (Table 1): pooled area under the ROC curve (AUROC, where 0.5 is chance and 1 a perfect ranking) 0.62 at high-confidence interactions, rising to 0.65 at the highest confidence, with per-species AUROC reaching 0.89. The signal does not appear to reflect phylogenetic inertia, because interactions are predicted at above-chance levels even for phyla not in the training set. For example, holding out *E. coli*’s clade from the training set, from its phylum Pseudomonadota down to the species itself, leaves its AUROC at 0.63 throughout. Indeed, for the 17 pathogens whose phylum is held out in at least one split, AUROC is essentially unchanged whether that phylum is in training or held out (Supplementary Material, Sec. S6.1).

**Table 1:** The Ising coupling*J* predicts experimentally validated protein interactions across the 19 human bacterial pathogens. Per-species area under the ROC curve (AUROC) for the gene-content coupling *J* ranking intra-species protein pairs against STRING v12.0’s *experimental* channel alone – not combined score, which embeds the co-occurrence signal *J* encodes – at high (*≥* 700) and highest (*≥* 900) confidence. *J* is unsupervised, uses gene presence/absence only, and has seen no interaction data; 0.5 is chance. Pooled across species, it matches STRING’s purpose-built co-occurrence channel. “pairs” is the number of intra-species protein pairs scored (millions). Rows ordered by AUROC (*≥* 700). The last two columns are read-outs of the same model: AUROC (*c_i_c_j_*) is the per-gene denoising-confidence product *c_i_c_j_*(*c_i_* = 1*/*(1 *− m_i_*^2^)) alone, a node-degree baseline; AUROC (sup.) is a supervised gradient-boosted head over *J* and *c_i_c_j_*, cross-validated within each species against the *≥* 700 channel. The supervised score (*∼*0.80) tracks the degree baseline *c_i_c_j_*, not the pair-specific coupling *J*: its gain over *J* is a node-degree bias, not new pairwise signal (text and Supplementary Material).

| species | pairs (M) | AUROC |  |  |  |
| --- | --- | --- | --- | --- | --- |
| | | $\geq 700$ | $\geq 900$ | $c_i c_j$ | sup. |
| *S. enterica* | 7.3 | 0.66 | 0.74 | 0.68 | 0.78 |
| *P. aeruginosa* | 11.8 | 0.65 | 0.66 | 0.70 | 0.80 |
| *Neisseria gonorrhoeae* | 1.3 | 0.64 | 0.65 | 0.75 | 0.81 |
| *Neisseria meningitidis* | 1.3 | 0.63 | 0.66 | 0.76 | 0.81 |
| *Streptococcus pyogenes* | 1.2 | 0.63 | 0.66 | 0.76 | 0.82 |
| *Bordetella pertussis* | 5.1 | 0.63 | 0.67 | 0.78 | 0.84 |
| *Clostridioides difficile* | 4.8 | 0.63 | 0.66 | 0.77 | 0.82 |
| *Haemophilus influenzae* | 1.3 | 0.63 | 0.65 | 0.78 | 0.82 |
| *E. coli* | 6.6 | 0.63 | 0.67 | 0.69 | 0.77 |
| *Klebsiella pneumoniae* | 9.1 | 0.62 | 0.67 | 0.65 | 0.75 |
| *Enterococcus faecalis* | 2.7 | 0.62 | 0.66 | 0.75 | 0.80 |
| *Helicobacter pylori* | 0.7 | 0.62 | 0.63 | 0.73 | 0.78 |
| *Acinetobacter baumannii* | 4.8 | 0.62 | 0.65 | 0.73 | 0.79 |
| *Vibrio cholerae* | 4.6 | 0.62 | 0.89 | 0.73 | 0.78 |
| *S. aureus* | 2.3 | 0.62 | 0.66 | 0.76 | 0.80 |
| *Campylobacter jejuni* | 1.0 | 0.61 | 0.63 | 0.75 | 0.80 |
| *Streptococcus pneumoniae* | 1.6 | 0.61 | 0.83 | 0.74 | 0.79 |
| *Mycobacterium tuberculosis* | 5.5 | 0.61 | 0.66 | 0.76 | 0.81 |
| *Listeria monocytogenes* | 3.0 | 0.61 | 0.64 | 0.77 | 0.81 |
| **Pooled (19 species)** | 75.9 | 0.62 | 0.65 | 0.74 | 0.80 |
| STRING co-occurrence channel | – | 0.64 | 0.67 | – | – |

A single matrix trained to denoise gene content matches STRING’s purpose-built co-occurrence channel (a phylogenetic-profiling pipeline; 0.62 versus 0.64). A supervised classifier over *J* and the model’s per-gene denoising confidence reaches AUROC 0.84 on *E. coli* with only that species held out – above the 0.77 that Table 1 reports for the production model, whose training set excludes the whole phylum (Supplementary Material, Sec. S6.1), narrowing the gap to supervised, sequencebased predictors such as ProteomeLM (Malbranke et al., 2026); but this lift over *J* is a per-gene node-degree effect rather than a pair-specific signal – *c_i_c_j_* alone reaches 0.77–0.80 – and it sharpens when the target’s lineage is in training, from 0.827 under whole-phylum hold-out to 0.843 with only the species held out, while *J* itself stays at 0.63 throughout (Supplementary Material, Sec. S6.1). This is a lower bound: the couplings driving the ancestral reconstructions carry externally validated interaction signal.

We also reasoned that the co-occurrence structure should contain information about gene essentiality, because the presence of essential genes should be strongly predicted by the rest of the genome. To test this hypothesis, we calculated each gene’s leave-one-out reconstruction probability: we delete a single gene from the *E. coli* genome, denoise with the best-fit model, and infer the probability that the deleted gene ought to be present. This reconstruction probability predicts essentiality above chance (AUROC 0.64) and, interestingly, is not due to information about conservation (how widespread a gene is across bacteria). Indeed, the denoiser’s single-site field *h_i_* (reflecting gene distribution across the training data) is actually *anti* - predictive of *E. coli* essentiality (AUROC 0.38), presumably because gene essentiality has an important lineage-specific component.

While the denoiser with a higher-order head did not greatly help in gene content reconstruction (see above), it does help here: the leave-one-out reconstruction probability predicts essentiality at AUROC 0.64 under the higher-order model, against 0.54 for the pairwise model. This may be because essentiality reflects a gene’s embedding in the global functional network of the cell, which is the kind of information captured by the higher-order head (Supplementary Material, Sec. S6). A classifier trained on the deleted gene’s presence probability at every step of the relaxation reaches AUROC 0.80 and, unlike the endpoint posterior, is flat across the leave-clade-out series (0.79–0.81 against 0.70–0.74) – still short of the supervised, sequence-based ProteomeLM (0.93–0.95), but using no amino acid sequence information (Supplementary Material, Sec. S6.2).

### Denoising improves the accuracy of phenotype prediction

A number of methods now exist for predicting microbial phenotypes from genomes (Koblitz et al., 2025; Fistarol et al., 2025; He et al., 2025; Zampieri et al., 2019). We recently developed one such method for predicting phenotypes based on gene content (Koldaeva et al., 2026) which, by comparison to other genome representations, has the advantage of being applicable also to ancestral gene content profiles, as reconstructed using reconciliation or profile-based methods. However, a significant limiting factor for ancestral phenotypic prediction is the accuracy of the reconstructed ancestral gene content, which in past work substantially curtailed predictive accuracy and certainty at deeper nodes of the bacterial tree (Koldaeva et al., 2026).

To determine whether denoised gene contents can improve predictive performance, we reinvestigated predictive accuracy and confidence using the noisy extant genomes from Koldaeva et al. (2026) as well as these genomes after denoising by our method. We tested three of the five predictors from that method, i.e. oxygen use (aerobe/anaerobe), monoderm/diderm cell envelope, and optimal growth temperature (OGT, in *^◦^*C), under increasing gene loss in extant genomes, which simulates the loss of gene family signal at deeper nodes associated with reconciliation-based reconstruction (Fig. 5). For the two binary traits the effect is large: at 90% gene loss, denoising improves oxygen-use MCC from 0.35 to 0.80 and cell-envelope (monoderm/diderm) MCC from 0.49 to 0.92 without affecting performance on near-complete genomes. Optimal growth temperature behaves differently, possibly because the signal is genome-wide, rather than restricted to a few marker genes: denoising a near-complete genome slightly reduces accuracy (RMSE 3.4 *→* 6.8*^◦^*C), the raw and denoised curves cross near 80% loss, and denoising helps only for highly incomplete input genomes (RMSE 11.7 *→* 8.7*^◦^*C at 90%), with a modest calibration gain (4.0 *→* 3.1*^◦^*C at 90%). These results suggest that denoised gene content profiles can improve gene-content-based phenotype prediction from partial or reconstructed ancestral genomes, most clearly for binary traits. Full per-phenotype numbers are provided in Supplementary Material S10.

**Figure 5:**
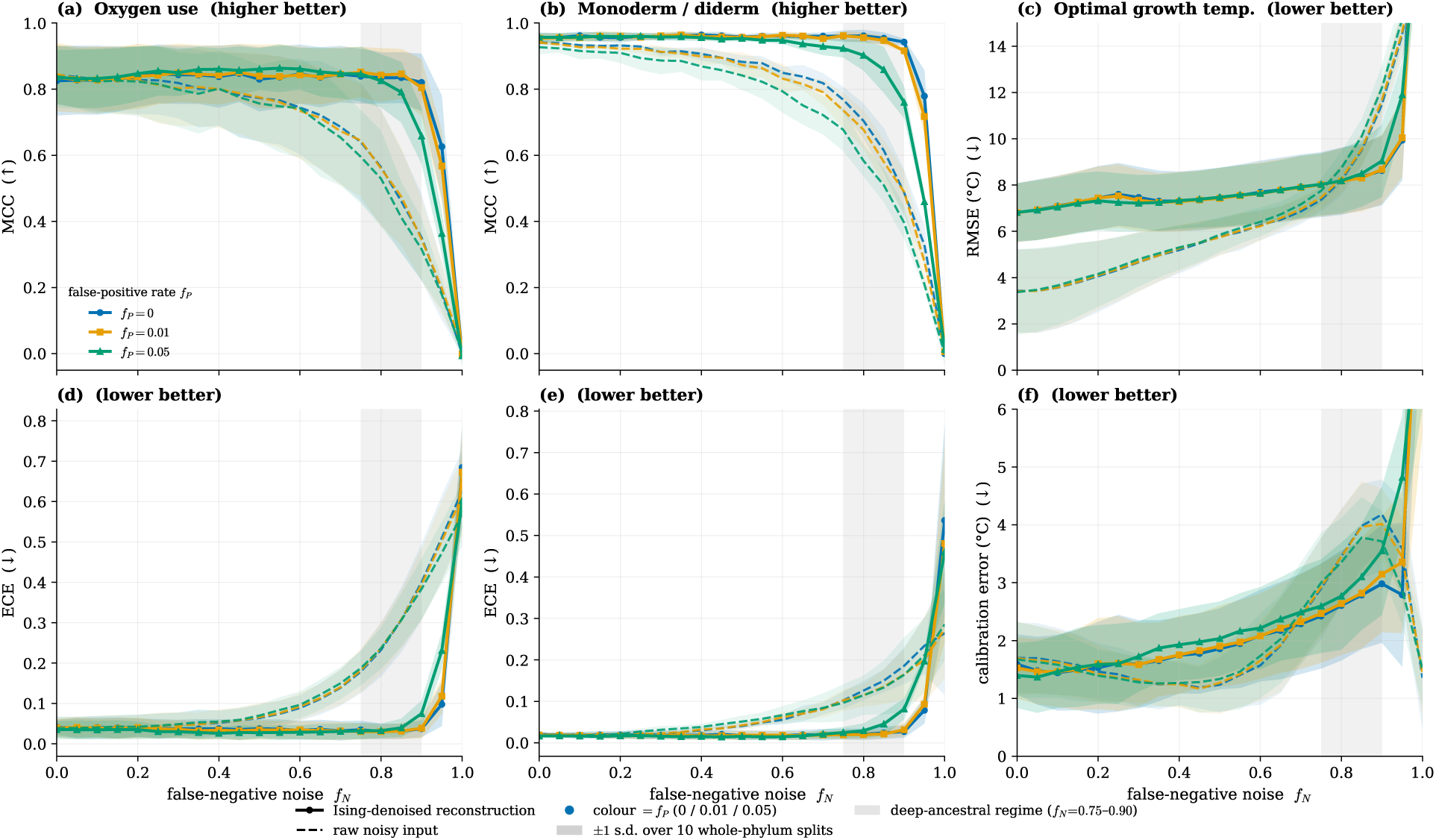
Denoising rescues phenotype prediction under gene loss, most clearly for the two binary traits. **(a,d)** oxygen use (aerobe/anaerobe), **(b,e)** monoderm/diderm cell envelope, and **(c,f)** optimal growth temperature (OGT), for three of the gene-content trait predictors of Koldaeva et al. (2026) against the false-negative (gene-loss) rate *f*_N_. The top row is predictive accuracy, in MCC for the two binary traits (higher is better) and RMSE in *^◦^*C for OGT (lower is better); the bottom row is calibration, in ECE for the binary traits and a calibration error in *^◦^*C for OGT (the stated uncertainty against the realized error, lower is better). Solid lines with markers are the denoised reconstruction, dashed lines are the raw noisy input, and colour is the false-positive rate *f*_P_ *∈ {*0, 0.01, 0.05*}*. Bands are *±*1 s.d. across the 10 whole-phylum cross-validation splits, and the grey band marks the deep-ancestral regime (*f*_N_ = 0.75–0.90). The entire 4,789-COG presence vector is corrupted before denoising, not only the genes the predictors read. At *f*_N_ = 1 MCC falls to zero for both binary traits, the control showing that the predictors read no gene content the corruption missed. The full protocol and the metric definitions are in Supplementary Material, Sec. S10.

**Figure 6:**
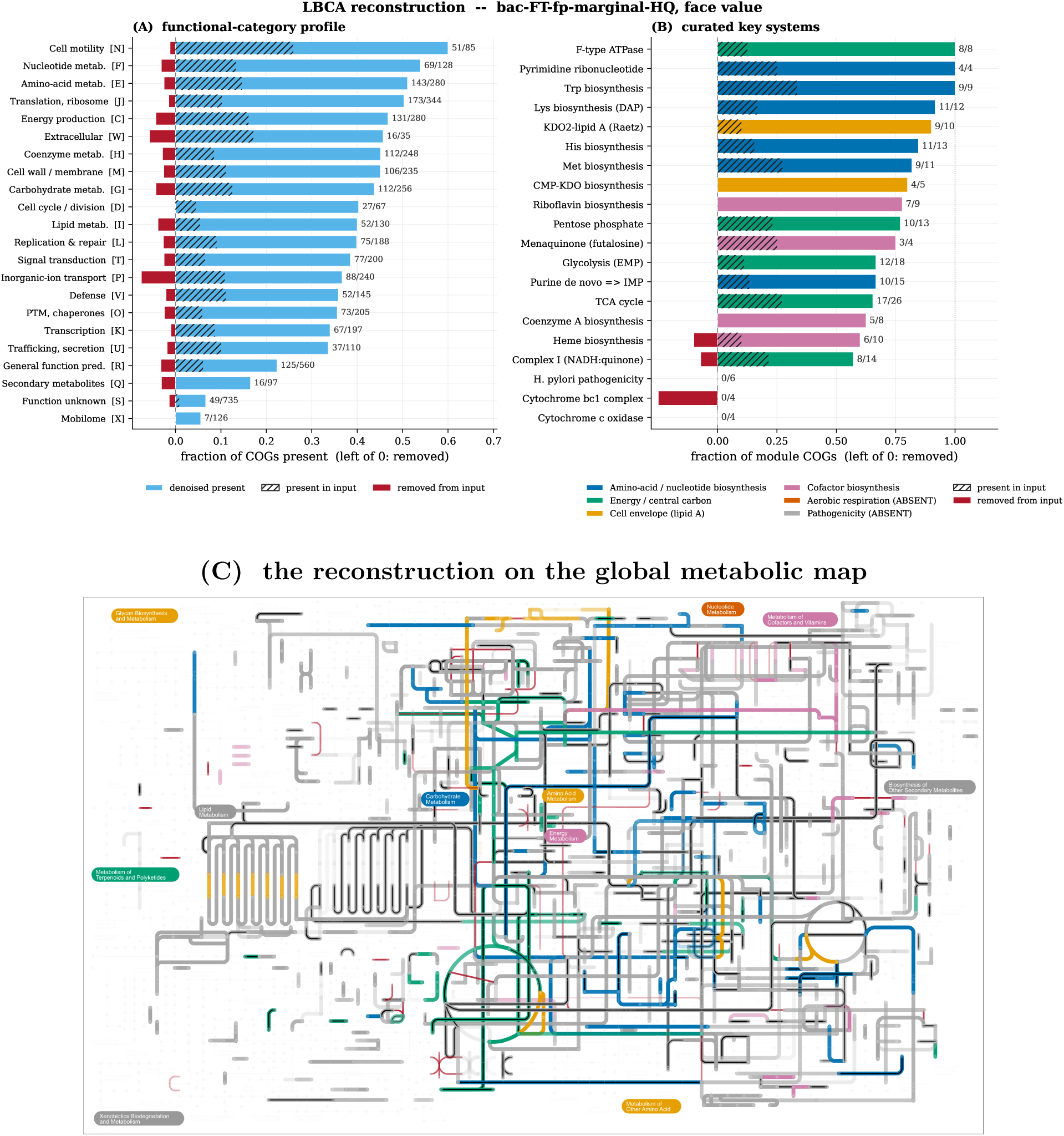
The reconstructed LBCA is a metabolically coherent bacterium. (A) per-COG functional-category profile and (B) curated key KEGG systems of the denoised reconstruction, as *thresholded* counts over an ensemble of ten models (a family is present when its mean posterior exceeds 0.5): the bar is the fraction of a category or module called present, the hatched portion was also supported by the input, the unhatched extension is recovery, and bars left of zero are families the input supported that are not called present after denoising. (C) The reconstruction on the iPath3 global metabolic map (Darzi et al., 2018). Each present family is a thick line in its curated key-system colour of panel (B) – amino-acid / nucleotide biosynthesis (**blue**), energy / central carbon (**green**), cell envelope (**orange**), cofactor biosynthesis (**purple**) – or neutral grey if it is not in a key system; opacity is the family’s ensemble posterior confidence (solid at *p*=1, fading to nothing at the *p*=0.5 call boundary). Families already present in the *input* carry a thin **black** centreline (the sparse scaffold the denoiser was given). The families added to the output are drawn in pure colour, with no black, so the reconstruction stands out against the input. The few removed families are red, opacity by how confidently they are absent. The sparse input, counted as the 493 families with any non-zero support, fills in to a coherent metabolism (1,519 present – 1,145 reconstructed, 374 kept). Bars left of zero are the 119 families that carried non-zero reconciliation support but are not called present after denoising; their support was very weak (median 0.01) and the denoiser raised 107 of them (median 0.24) without reaching 0.5. Only metabolic COGs are depicted.

**Figure 7:**
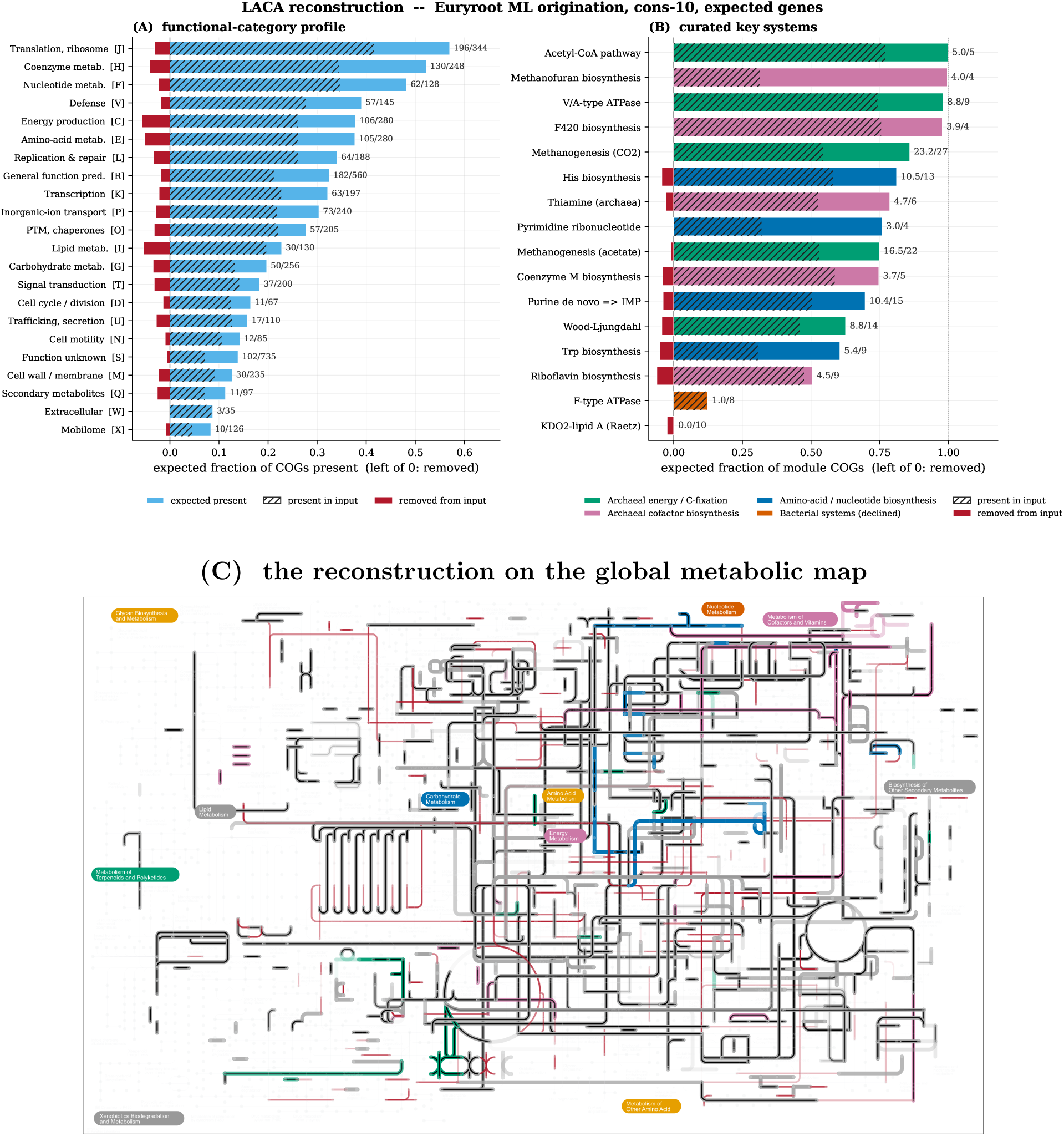
The reconstructed LACA is a hyperthemophilic methanogen, consistent with recent analyses. (A) per-COG functional-category profile and (B) curated key KEGG systems of the denoiser’s reconstruction of the focal Euryarchaeota-rooted input, as *expected* gene counts over an ensemble of ten models: posteriors are summed rather than thresholded, so a category’s denoised content is Σ*_i_* p*_i_*. The bar is that expected content, the hatched portion is the posterior mass shared with the input (Σ*_i_* min(p*_i_*^in^,p*_i_*^out^)), the unhatched extension is the mass the denoiser adds, and bars left of zero are the mass it withdraws. Panels (A) and (B) of Fig. 6 are thresholded counts rather than expected mass, so the two figures are not directly comparable bar for bar. The denoised LACA completes archaeal features such as for example the V/A-type ATPase (9 of 9). (C) The reconstruction on the iPath3 global metabolic map (Darzi et al., 2018). Each present family is a thick line in its curated key-system colour of panel (B) – archaeal energy / C-fixation (**green**), cofactor biosynthesis (**purple**), amino-acid / nucleotide biosynthesis (**blue**), the declined bacterial systems (**vermilion**) – or neutral grey if it is not in a key system; opacity is the family’s ensemble posterior confidence (solid at *p*=1, fading to nothing at the *p*=0.5 call boundary). Families already present in the *input* carry a thin **black** centreline (the scaffold the denoiser was given); the families it *adds* in the output are drawn in pure colour, with no black. The removed families are red, opacity by how confidently they are absent. In expected-count terms the input carries 1,032 genes of posterior mass and the reconstruction 1,322: 909 kept, 413 added and 123 withdrawn. Thresholded at 0.5 the same reconstruction calls 1,321 families present, of which 880 were already called present in the input. Only metabolic COGs are depicted.

### Applying the denoiser to empirical ancestral reconstructions

We and others have previously attempted to reconstruct the gene content of early ancestors (including the last archaeal and last bacterial common ancestors), in order to draw inferences about their cell biology and metabolic potential. As discussed above, different reconstruction methods have different properties (Williams et al., 2024). Phylogenetic reconciliation is a useful approach for studying early microbial evolution because inferences can accommodate HGT, which is known to be extensive among prokaryotes. However, reconciliation-based ancestral inferences are sometimes very conservative because gene tree noise can be mis-interpreted as support for a high HGT rate, which then reduces the number of gene families that can be mapped to the root of the tree. The extent of this conservatism varies among analyses. When gene trees can be accurately reconstructed (for example, when few species are analysed, so that gene family alignments are relatively long with respect to the possible number of gene trees), reconciliation-based methods give similar results to more liberal approaches (Csűrös, 2026). When the species tree is large and the root note is ancient - for example in the recent work of Davín et al. (2025) - then reconciliation methods can map very few genes to the root with confidence. In addition, recent work suggests that the fit of the reconciliation model is an important factor, with better-fitting, branch-heterogeneous models that allow the rates of duplication, transfer and loss to vary across the species tree (e.g. Huang et al. (2026)) recovering much more complete ancestors than the simpler branch-homogeneous models that were, until recently, the norm (e.g., (Williams et al., 2017; Coleman et al., 2021; Davín et al., 2025)).

To investigate whether the denoiser can recover realistic ancestors from a range of different inputs, including partial reconstructions from analyses with simple reconciliation models, we applied it to published gene content reconstructions from two recent papers (Davín et al., 2025; Huang et al., 2026). To compare to the alternative gain-loss-duplication (GLD) model (Csűrös, 2026), we ran that model on the same datasets using our recent Recount implementation (Szöllősi and Williams, 2026), which enables the method to scale to larger datasets. In what follows, we comment on the inferred biology of the denoised reconciled ancestors ((Davín et al. (2025), (Huang et al., 2026)) and compare agreement between distinct reconstruction approaches before and after denoising, and c.

### The last bacterial common ancestor

The reconciliation-based reconstruction of the LBCA, the bacterial root node of the time-calibrated, genome-scale reconciliation of Davín et al. (2025) (rooted following Coleman et al., 2021), is extremely conservative: of 493 families with any appreciable signal, only six reach *PP ≥* 0.5. The denoiser greatly expands this, inferring 1,519 families to be present: adding 1,145 families with no appreciable signal in the raw reconciliation-based input. A further 119 families entered with weak support (median input posterior 0.01); the denoiser raised 107 of them and lowered the remaining 12, in neither case reaching the 0.5 calling threshold (median output 0.24).

The denoised LBCA is metabolically coherent, encoding 77 full KEGG modules (versus 3 in the raw input), with a further 175 modules at least half complete. This falls within the range of extant genomes: 47.0% of its KEGG modules with at least four member families are left fragmentary at 1,519 families, against 47.1 *±* 5.0% for the 41,696 present-day genomes of 1,300–1,500 families in the held-out data.

Based on functional annotations of the denoised LBCA Davín et al. (2025) and phenotypic predictions (Koldaeva et al., 2026), the LBCA is inferred to be a free-living, motile, anaerobic diderm. It is predicted to have encoded largely complete pathways for the biosynthesis of amino acids, nucleotides and various cofactors as well as a gram-negative-type envelope, peptidoglycan precursors, and bacterial cell division proteins consistent with a diderm bacterial root (Coleman et al., 2021). It was also predicted to encode a bacterial Wood-Ljungdahl pathway, Rnf complex, and membrane-bound hydrogenases, as well as both an F-type and archaeal-vacuolar (AV)-type ATP synthase. The latter is consistent with a recent study on the evolution of ATP synthases, that emphasizes the presence of both types in a diversity of Bacteria and potentially also in LBCA (Mahendrarajah et al., 2023). While the denoiser predicted the presence of a Cytochrome bd-type quinol oxidase (subunit 1 and 2) at low PPs (sparse: PP 0.6907 and 0.5994, GLDmin1: PP 0.6040 and 0.3129, GLDmin4: PP 0.4931 and 0.2612), phenotypic predictions from the denoised reconstructions (Koldaeva et al., 2026) indicate that LBCA was an anaerobe, consistent with homologues of this enzyme complex not being exclusive to aerobes. Indeed, the overall effect of the denoiser was to reduce support for aerobic metabolism in LBCA, which was low in the reconciliaton-based input (0.015) but appeciable in the GLD-based reconstructions (0.435-0.498); all denoised reconstructions were confidently predicted to be anaerobes (P ¿ 0.99).

Among the families the denoiser removes are gene families that clash with the broader picture. This is clearest in the dense copy-number reconstructions, where several Heme/copper-type cytochrome/quinol oxidase subunits enter the input at probability 1.0 and are reduced to below 0.05. In the sparse reconciliation reconstruction, by contrast, the input asserts almost nothing to begin with: of the six families at *PP ≥* 0.5, the four strictly above 0.5 are all retained, and of the two sitting exactly at 0.50 one is raised (COG1530, ribonuclease G/E, to 1.00) while the other falls (COG1784, TctA family transporter, to 0.23). The two cases together show the denoiser weighing isolated genes against the genomic context: confidently called aerobic machinery is pushed down in a context dominated by anaerobic metabolism, while weakly supported families are pulled up but not far enough to be called present.

### The last archaeal common ancestor

We recently used a reconciliation approach that integrates improved models of gene evolution allowing for distinct DTL rates across taxa, to resolve the archaeal root and reconstruct the ancestral gene content of the Last Archaeal Common Ancestor (LACA) (Huang et al., 2026). This work, based on reconciliations using arCOGs, identified a narrow root region within or between the’Euryarchaeota’ and all other archaea and inferred a free-living, (hyper)thermophilic, methanogenic LACA. The arCOGs with a presence probability above 0.5 in LACA (Huang et al., 2026) correspond to 1,018 COG families (see methods) and were subjected to denoising. Specifically, the denoiser model mapped 1,321 gene families with presence probability above 0.5 to LACA which comprises 880 of those previously identified and 441 newly added families. Conversely, the denoiser led to the removal of 138 families.

Overall, the gene set is consistent with previous inferences of a methanogenic LACA (Huang et al., 2026) and includes pathways such as the Wood-Ljungdahl pathway and an associated methanogenesis module as well as archaeal informational processing machinery. Some of the added families such as for example the AV-type ATP synthase C and I, seem to complete pathways and enzyme complexes. Conversely, bacterial information processing and cell biological machinery as well as enzyme complexes remain nearly absent as evidenced by the lack of the bacterial F-type ATP synthase. This is reassuring because a potential concern about the denoiser approach might be the risk of simply filling in gaps based on average gene family frequencies across prokaryotes. However, both the denoised LBCA and LACA reconstructions reflect domain-specific gene repertoires, suggesting that the model has learned genome grammar rather than a single average genome.

### Conservative and liberal reconstructions converge

As discussed in the introduction, different ancestral reconstruction methods result either in conservative (incomplete, gappy) or liberal (overly-complete) reconstructions. The previous sections illustrate how the denoiser can improve recent reconciliation-derived reconstructions. Next, we investigated its performance on empirical reconstructions from the same datasets using a phylogenetic profile-based method, GLD, which appears to moderately over-estimate ancestral contents under some conditions.

Encouragingly, the denoiser appears to move opposite kinds of input toward each other, substantially reconciling differences in gene content and the resulting cell biological and metabolic interpretation. At the LBCA, the reconciliation and profile-based reconstructions disagree sharply: the reconciliation recovers only faint signal at the root (the sparse input above), while a profile-level gain-loss-duplication reconstruction (recount, a Count-class model with a tree-Brownian autocorrelated log-rate prior; Szöllősi and Williams, 2026; Csűrös, 2010) infers a dense ancestor of *∼*2,000 confidently present families. The denoiser greatly expands the reconciliation-based input, from the 493 families carrying any support to 1,519 called present at posterior *>* 0.5, adding 1,145 cooccurring members of confident modules, and substantially prunes the GLD-based input, judging *∼* 20% (*∼*394 families) of the confidently present input to be false, with a denoised reconstruction of 1,836 gene families. The denoised LBCAs share 1,430 families (Jaccard 0.75–0.76, where 1 is identical gene sets and 0 disjoint; Supplementary Material, Sec. S7). The remaining disagreements include a set of poorly-characterised families with accessory or defense functions (present in the GLD-based denoised LBCA) and some core metabolic genes that fill gaps in otherwise incomplete pathways in the reconciliation-based, denoised LBCA. The only substantive disagreement between the two denoised reconstructions relates to the completeness of the quinone-linked respiratory chain, although both reconstructions agree that LBCA was likely an anaerobe lacking a terminal oxidase, as expected.

The sparseness of reconciliation-based reconstructions results, in part, from a difficulty in distinguishing phylogenetic noise from biological processes such as HGT. It therefore seems likely that improving the fit of the reconciliation model should improve ancestral reconstruction and result in larger and more biologically realistic root gene content inferences. Indeed, these improvements are evident in the LACA analyses, which are drawn from a recent study that used a recently-developed tool, AleRax, that implements more flexible reconciliation models than have been available hitherto. By combining a root origination prior with clade-wise duplication, transfer and loss rates, the raw reconciliation-based reconstruction recovers 1,018 gene families at the root, and shows reasonable agreement with the GLD reconstructions (e.g. Jaccard = 0.59 for the min-4 condition). By contrast, failure to optimise the root origination prior by maximum likelihood greatly reduced the number of gene families confidently inferred to the root. After denoising, agreement among all analyses was substantially improved, with the biggest gains occurring where disagreement among inputs was greatest (Fig. 8).

**Figure 8:**
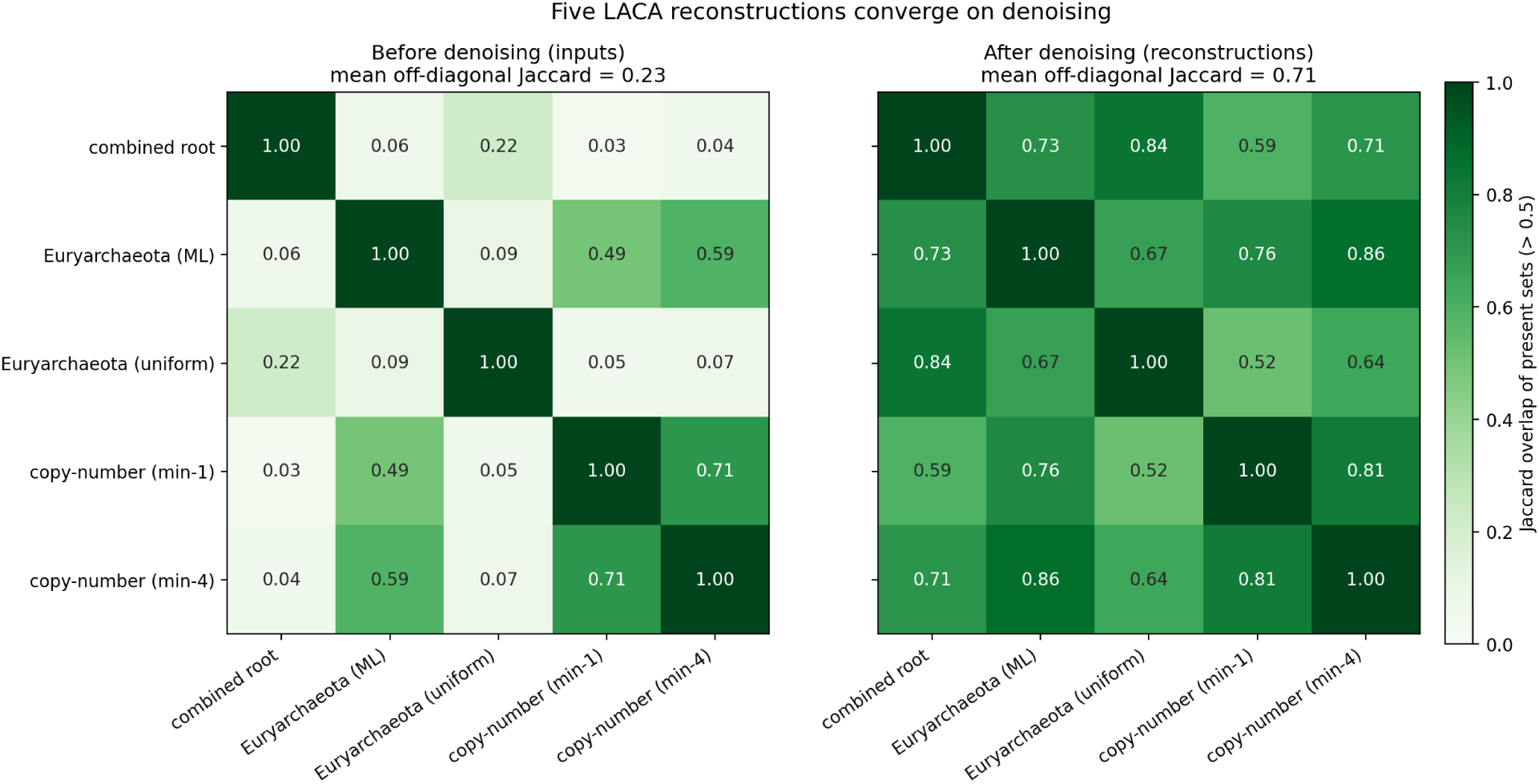
The five denoised LACA reconstructions converge. Pairwise Jaccard overlap of the present-gene sets (*>* 0.5) of the five reconstructions, each denoised identically from a different upstream archaeal input, on a shared colour scale. *Left*, the raw inputs (families the input calls present at posterior *>* 0.5): discordant, mean off-diagonal Jaccard 0.23, falling to 0.03 between the combined root and the dense copy-number reconstruction, and only 0.09 even for the two Euryarchaeota rootings, which differ only in how root gene origination is modelled (maximum-likelihood vs. uniform). *Right*, the denoised reconstructions: concordant, mean 0.71. The two uniform-origination roots (the combined root and Euryarchaeota uniform) are most similar to each other (0.84), and the focal maximum-likelihood root (Euryarchaeota ML) is most concordant with the copy-number min-4 ancestor (0.86). The two copy-number rows are dense gain-loss-duplication reconstructions thresholded at gene-occurrence 1 and 4. Generated by `scripts/plot_laca_jaccard_heatmap.py`.

## Discussion

Here, we have attempted to improve ancestral gene content estimation by making use of interactions among genes. To do so, we established a prior over the joint structure of gene content, learned from extant genomes and validated across the major bacterial and archaeal lineages. We used whole-phylum holdouts to assess model performance such that test and training data were separated by 1 or more billion years of independent evolution (barring HGT), to encourage the denoiser model to learn features of gene content that generalise across clades, rather than relying on rote memorisation of clade-specific patterns. Several observations suggest that this approach was successful. Denoisers trained in this way appear to be powerful for accurately reconstructing extant genomes from highly partial inputs under whole-phylum holdout, and greatly increase agreement between ancestral reconstructions inferred using methods that make different kinds of errors, filling in gaps in overly-conservative reconciliation-based reconstructions, and pruning back overly-generous profile-based inferences. The learned couplings predict physical protein interactions and gene essentiality, and the denoiser correctly identifies and removes what are likely to be false-positives in input reconstructions. Finally, our analyses also show that denoising genomes prior to gene content-based prediction of phenotypic traits (as in (Koldaeva et al., 2026)) can greatly improve performance, particuarly when the input genomes are partial.

A learned co-occurrence prior raises a long-standing concern: genes can appear coupled because the genomes that carry them share ancestry, so co-occurrence across genomes confounds vertical inheritance with functional dependence (Felsenstein, 1985). Phylogeny-aware methods address this directly, testing for correlated gain and loss on the species tree (Pagel, 1994; Barker and Pagel, 2005) or modelling the joint evolution of many genes along it (Liu et al., 2023); our model does not, treating genomes as independent samples. Three features of the validation bear on whether this matters. The interactome benchmark scores *J* against STRING’s experimental channel, a ground truth with no co-occurrence component, so the signal it recovers is not shared ancestry alone. The whole-phylum hold-out means recovery cannot rest on the clade structure of the training genomes. And at the archaeal node, the denoiser does not import bacterial machinery, which a model echoing the prevalence of bacterial genomes in training would. These show that *J* carries functional signal beyond ancestry; they do not make its raw couplings free of phylogenetic correlation, and a tree-aware fit (Liu et al., 2023) would be a natural way to separate the two.

Several recent models also learn from gene content across genomes, but occupy different niches. ProteomeLM (Malbranke et al., 2026) and genomic language models (Hwang et al., 2024) read protein relationships from proteome- or neighbourhood-scale context, keep them implicit in a neural network, and operate on present-day genomes; our couplings are explicit and signed, and the model is generative, used to reconstruct genomes rather than to score interactions. Large-scale phylogenetic profiling (Tremblay et al., 2021) scores gene pairs one at a time, and an interesting recent reconstruction method imputes each gene’s presence separately on the species tree (Mattick et al., 2026). The specific contribution of the approach described here is that one gene’s reconstruction is used to inform another, by fitting a joint gene content model once per species across the archaeal and bacterial tree, and then using it to denoise.

The biological reconstructions based on denoised gene contents are consistent with the emerging picture of the deepest nodes. The denoised LBCA is predicted to be a free-living, motile, diderm, biosynthetically autonomous bacterium with a chemiosmotic, quinone-linked but anaerobic metabolism. This is in line with the reconciliation-based rooting of the bacterial tree between Terrabacteria and Gracilicutes (Coleman et al., 2021), and the expected absence of a terminal heme-copper oxidase considering that LBCA predates the Great Oxidation Event by well over a billion years (LBCA 4.4–3.9 Ga, the GOE to *∼*2.4–2.3 Ga) (Davín et al., 2025; Lyons et al., 2014). The denoised LACA represents a free-living archaeaon that encoded an archaeal information machinery as well as a methanogenesis pathway in line with recent inferences (Huang et al., 2025). Unsurprisingly, terminal oxidases were acquired later, and independently, in many descendant lineages, most of them after the Great Oxidation Event (Williams et al., 2017; Davín et al., 2025).

The current method has some limitations The COG vocabulary misses lineage-specific and unannotated genes, so the true ancestral counts exceed the COG-mappable counts we report. The denoiser can correct false negatives that are inconsistent with learned co-occurrence, but it cannot recover a module for which the upstream reconstruction gave no evidence at all, nor can it reject a false positive that happens to be consistent with the learned gene content grammar. And while we trained denoiser models under a range of false positive and false negative rates, to investigate the effect of both contamination and phylogenetic noise on ancestral inferences, the extent to which these rates recapitulate the properties of real data remain unclear, particularly in deep time. Here, we investigated how denoising improves agreement between distinct reconstructions of LBCA and LACA, providing additional support for some key conclusions about the nature of these ancestors. As the denoiser appears to learn general, cross-domain features of gene content grammar, the approach may also be a promising one for addressing ongoing debates about the nature of their ancestor, the last universal common ancestor, as methods continue to be developed.

## Materials and Methods

### Genomes and Gene Family Vocabulary

We assembled a set of species-representative genomes from GTDB r220 (Parks et al., 2022), i.e. by using one genome per named prokaryotic species: 113,104 genomes, 107,235 bacterial and 5,869 archaeal, spanning more than 100 phyla. Their gene content was estimated on basis of COGs and encoded over *N* = 4,789 COG families (Galperin et al., 2021). Specifically, each genome was represented as a vector ***x*** *∈ {−*1, +1*}^N^* with one entry per COG family, +1 if that family is present in the genome one or more times and *−*1 if it is absent. All models were trained under a 10-way cross-validation, holding out whole phyla (validation fraction 0.2 by genome count), because a model trained on close relatives could recover gene content from phylogenetic inertia alone. We trained a’generalist model’ using the full set, including incomplete metagenome-assembled genomes, whose unassembled families appear as false negatives of the kind the ancestral problem presents. Furthermore, domain specialist models (bacterial for the LBCA, mixed for the LACA) were re-split within a domain and subsampled to *∼*15,000 training and *∼*3,000 validation genomes. For high-completeness analyses, we additionally required CheckM completeness *≥* 90% (bacteria) or *≥* 80% (archaea) and contamination *≤* 5%. The archaeal reconciliations used for denoising are based on arCOG gene family trees (Huang et al., 2025), with each arcog potentially corresponding to more than one COG family. Thus, we mapped each arCOG to its corresponding COG using a arCOG database mapping file and combined the presence probabilities of all arCOGs assigned to the same COG by noisy-OR, *P* = 1 *− _i_*(1 *− p_i_*), under which a COG is present if at least one of its arCOGs is. We applied this translation identically to every arCOG-native input, so that differences between those inputs reflect the upstream reconstructions rather than the translation.

### The Model and Mean-Field Denoising

We modelled the distribution over genomes as the pairwise (Ising) form (1), in which *h_i_* is the field on family *i* and *J_ij_*the coupling between families *i* and *j*, i.e. positive when two families tend to co-occur and negative when they exclude one another; ***J*** is symmetric with zero diagonal, so that no family couples to itself. Denoising is the problem of finding the high-probability genome consistent with a corrupted observation. We solved it by mean-field relaxation, in which each family responds to the average state of its partners rather than to their joint configuration: starting from the noisy input, we re-estimated each family as

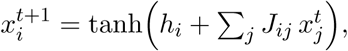

where *x*^t^*_i_* ɛ [-1,1] is the soft presence of family *i* after *t* updates, and one sweep applies this update to all *N* families. We trained end to end through a fixed number of sweeps *T*, each sweep corresponding to one layer of the network. We tried different numbers of layers *T ∈ {*8, 12, 16, 20*}* and report *T* = 20. Four terms are learned on top of (2): a reaction-field correction that removes the self-reinforcement bias of naive mean field (Supplementary Material, Sec. S1.3; Thouless et al., 1977); step-indexed gates setting, per family and sweep, how far to trust the corrupted observation against the couplings; a temperature, learned from the fraction of families present in the input, setting how decisively each sweep affects presence probabilities; and conditioning of those gates, through a small network over 419 functional modules, on how complete each module already is. We compared three variants at matched depth: couplings alone; couplings plus a learned third-order term on the families themselves, which captures dependencies among three or more families rather than just pairs; and that model plus 1,000 unobserved variables coupled to the families. We used the second. The unobserved variables improve recovery only marginally and at substantial cost – a mean gain of 0.005 MCC across the noise spectrum for a 23% increase in parameters (31.7M vs. 25.8M) – so the third-order term alone is the production architecture. It has *∼*29M parameters, *∼*23M of them couplings.

### Training

From each clean genome, we synthesised a noisy observation by deleting each present family with probability *f*_N_ and inserting absent families with probability *f*_P_ = 0.01. We drew *f*_N_ per genome from a curriculum that grows harder over training, ending with most of its mass in the deep-ancestral band *f*_N_ *∈* [0.75, 0.90], and generated four copies of each genome, three corrupted and one clean, on up to eight A100 GPUs. The objective combines a cross-entropy between the recovered and the clean genome with a term asking how well each family is predicted from the rest of the same clean genome, which fits fields and couplings directly to observed co-occurrence, independently of the corruption. From the depth-20 model we fine-tuned the domain specialists and a ladder of false-positive curricula: uniform per-family noise, then coherent grafted modules, and finally dense contamination by common families. Each genome contributed rescue copies, with present families deleted for the model to restore, and prune copies, with absent families added in proportion to their frequency across genomes for the model to remove, both scored against the unchanged clean genome. A final LACA fine-tune penalised the variance of the output across a genome’s corrupted copies, so that sparse and dense corruptions of one genome map to a single reconstruction.

### Benchmarks Against External Data

We benchmarked the couplings, averaged over the ten cross-validation models and average-product corrected, against the experimental channel of STRING v12.0 (Szklarczyk et al., 2023) over the 19 human bacterial pathogens of Humphreys et al. (2024), mapping each proteome to COGs with diamond. We estimated the essentiality of each *E. coli* family as the confidence with which the model restores that family when it alone is deleted from the genome, and compared this with the experimental data of Goodall et al. (2018). The phenotype tests used three gradient-boosted-tree predictors (oxygen use, cell envelope and optimal growth temperature; A. Koldaeva, unpublished) under the same phylum hold-outs, corrupting every family in the genome before denoising, so that the classifier could not read uncorrupted families outside its own feature set.

### Application to Ancestral Nodes

The reconciliation posteriors we denoised are the bacterial root of the time-calibrated reconciliation of Davín et al. (2025) for the LBCA, rooted following Coleman et al. (2021), and the Euryarchaeota-rooted archaeal reconciliation of Huang et al. (2025) for the LACA. We relaxed each through the ten-model ensemble of the appropriate specialist, entering each family at its reconciliation presence probability *p* as *x* = 2*p −* 1 and calling that family present when the ensemble posterior exceeded 0.5. The dense profile-level inputs come from recount, a gain-loss-duplication reconstruction with a tree-Brownian autocorrelated rate prior (Szöllősi and Williams, 2026; Csűrös, 2010), on the same data. Module coherence is the fraction of KEGG modules, among those containing at least four families, whose completeness falls between 25 and 74%, benchmarked against present-day genomes of similar gene-family count.

### Reproducibility

Code, figures and tables can be regenerated from committed data with a script that compares outputs against stored values; checkpoints and training data are on Zenodo.

## Acknowledgements

We thank Anzhelika Koldaeva for the oxygen use classifier and the phenotype-under-gene-loss data, and the OIST Saion HPC team. Computation used the OIST Saion GPU cluster. TAW, AS and GJSz were supported by a grant from the John Templeton Foundation (63451). The opinions expressed in this publication are those of the authors and do not necessarily reflect the views of the John Templeton Foundation.

## Data and code availability

Source code is at https://github.com/ssolo/gene-content-grammar. Trained models, training data, and the analysis pipeline are archived on Zenodo (concept DOI 10.5281/zenodo.20526264); the per-COG reconstructions of the LBCA and LACA, with input copy number, ensemble posterior, and cross-split standard deviation per family, are released alongside. Code and data are made available under a Creative Commons Attribution–NonCommercial 4.0 licence

