## Supplementary Material for "Improved ancestral genome reconstruction using a learned gene-content grammar"

[1]Department of Biological Physics, Eötvös Loránd University, Budapest, Hungary
[2]Model-based Evolutionary Genomics Unit, Okinawa Institute of Science and Technology, Okinawa, Japan
[3]School of Biological Sciences, University of Bristol, Bristol, United Kingdom

This supplement collects the detailed validation, training, and convergence analyses summarised in the main text. Section S1 gives the model architecture and training detail; Section S2 the whole-phylum recovery stratified by phylum; Section S3 the per-node model comparison and ground-truth recovery on extant genomes; Section S4 the realistic false-positive training and the contamination-rejection frontier; Section S5 the genome-mass normalization of the coupling; and Section S6 the external validation detail (interactome and gene essentiality). The convergence story follows: Section S7 the conservative-build versus generous-prune operation at the LBCA; Section S8 the five-input LACA convergence; and Section S9 the per-gene confidence and the shared cellular core. Throughout, the noise channels are a false-negative (deletion) rate $f_\mathrm{N}$ and a false-positive (spurious-gene) rate $f_\mathrm{P}$, and all internal validation uses the ten whole-phylum cross-validation folds described in the main text.

#### S1 Model architecture and training detail

##### S1.1 Ising distribution and mean-field relaxation

A genome is $\boldsymbol{x} \in \{-1, +1\}^N$ over $N = 4{,}789$ orthologous gene families (COGs), $x_i = +1$ for presence. Gene content is modelled as a pairwise Ising (Boltzmann) distribution

$$P(\boldsymbol{x}) \;\propto\; \exp\Big(\sum_i h_i\, x_i \;+\; \sum_{i<j} J_{ij}\, x_i x_j\Big), \tag{S1}$$

with learned fields $\boldsymbol{h} \in \mathbb{R}^N$ and a symmetric, hollow coupling matrix $\boldsymbol{J}$ ($J_{ii} = 0$). Intuitively, the field $h_i$ says how common family $i$ is on its own, and the coupling $J_{ij}$ says whether families $i$ and $j$ tend to occur together ($J_{ij} > 0$) or to exclude one another ($J_{ij} < 0$); the exponent is a score that is highest for the gene combinations the training genomes actually exhibit, so a genome that respects the learned co-occurrence structure is assigned high probability. Fitting a single joint distribution, rather than scoring gene pairs independently, lets $\boldsymbol{J}$ separate directly coupled families from those correlated only through shared partners, the advantage direct-coupling analysis exploits for residue contacts (Weigt et al., 2009); joint pairwise models of gene presence/absence have been used to predict functional and physical interaction (Croce et al., 2019; Fukunaga & Iwasaki, 2022), building on phylogenetic profiling (Pellegrini et al., 1999; Kensche et al., 2008). The maximum-probability configuration consistent with an observation is approached by the naive mean-field fixed-point iteration on the magnetisations,

$$x_i^{t+1} \;=\; \tanh\Big(h_i + \textstyle\sum_j J_{ij}\, x_j^t\Big), \qquad t = 0, \dots, T-1, \tag{S2}$$

initialised at the noisy observation $\boldsymbol{x}^0$. Each sweep re-estimates every gene from the current guess of its neighbours; iterating drives the configuration toward a self-consistent, high-probability genome, the denoised output. Reconstructing a corrupted binary pattern under a learned distribution is the task associative-memory and Boltzmann-machine models address (Hopfield, 1982; Ackley et al., 1985), here over an interpretable model fit to gene content. The number of sweeps $T$ is the *relaxation depth*: it sets how far the configuration may move from the observation before the prediction is read out, so small $T$ trusts the input and large $T$ lets the learned couplings propagate evidence further. $T$ is an iteration count, not a softmax temperature; the temperature is a separate learned quantity (Section S1.3).

#### S1.2 A ladder of three architectures

We validate three nested architectures, in increasing expressive power.

**(i) Plain pairwise Ising (NoHidden, no higher-order).** The bare model (S1)–(S2): visible genes coupled only through $\boldsymbol{J}$, with the gated, step-dependent update of Section S1.3 but no beyond-pairwise term.

**(ii) Ising with hidden units and a higher-order head.** Pairwise couplings cannot express that a module is present only when all its members are. We augment the visible state with a hidden vector $\boldsymbol{z} \in \mathbb{R}^H$, $H = 1{,}000$, coupled through learned maps $A \in \mathbb{R}^{N\times H}$ and $U \in \mathbb{R}^{H\times N}$,

$$\boldsymbol{x}^{t+1} = \tanh\big(\boldsymbol{h} + \boldsymbol{x}^t\,\widetilde{\boldsymbol{J}} + \boldsymbol{z}^t A^\top\big), \tag{S3}$$

$$\boldsymbol{z}^{t+1} = \phi\big(U\,\boldsymbol{x}^{t+1}\big), \tag{S4}$$

with $\widetilde{\boldsymbol{J}}$ the symmetrised visible coupling and $\phi$ a pointwise nonlinearity. Marginalising the hidden units induces effective couplings of arbitrary order among the genes, so (S3)–(S4) is a higher-order Ising denoiser.

**(iii) Ising with a higher-order head, no hidden units.** The same beyond-pairwise interactions can be carried by a learned third-order correction head attached directly to the visible state, with *no* explicit hidden vector. This matches (ii) at matched depth while being lighter, is represented Fig S1, and is our production architecture.

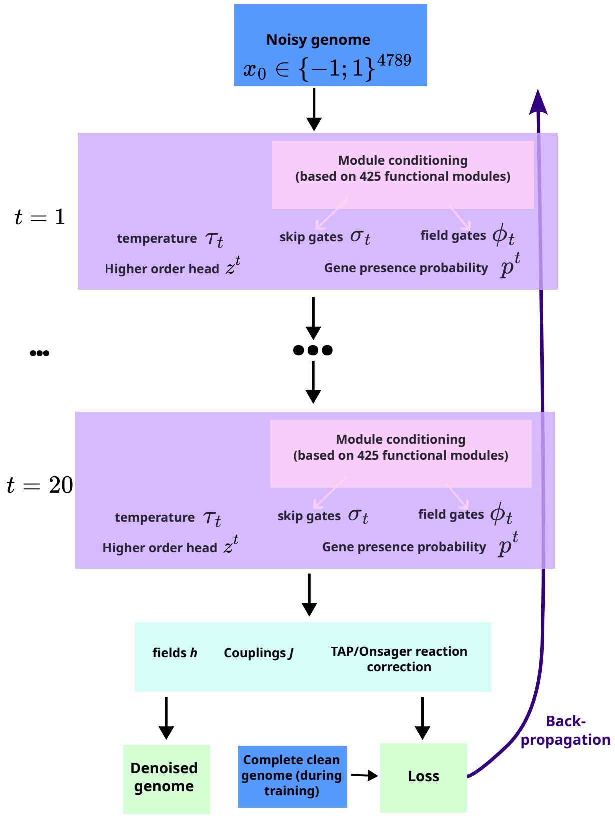


**Figure S1: Representation of the production version of the denoiser.** A corrupted genome is provided as input to the denoiser, and goes through a series of 20 steps. At each step, gates are applied to the coupling term and to the input, and an adaptive temperature is applied to the update. The output of the final step is a denoised genome, at inference time, or is used to compute a loss to optimize the parameters during training, through backpropagation.

**Nomenclature.** We use one consistent naming scheme throughout. **Pairwise** $T$ is architecture (i), the plain pairwise Ising at relaxation depth $T$; **HO** $T$ is architecture (iii), the production no-hidden higher-order model; **HO+hidden** $T$ is architecture (ii). The depth-20 HO model is the *generalist* from which all fine-tunes branch. Three domain fine-tunes of HO $T$=20 follow: **mix-FT** (mixed-domain rebalance – the best-validated generalist, used for the *archaeal* LACA), **bac-FT** (bacteria-only specialist, used for the *bacterial* LBCA), and **arc-FT** (archaea-only, a control). mix-FT and bac-FT are distinct models for different nodes. False-positive-tolerant variants (the high-FP line) carry an **-fp** suffix (**mix-FT-fp**, **bac-FT-fp**); these are the production models fine-tuned with the *realistic* coherent false-positive curriculum of Section S4, and are the FP-tolerant models used in every reconstruction below that runs under injected false positives.

#### S1.3 Gated update and adaptive temperature

A fixed mean-field map cannot denoise gently at low noise and aggressively at high noise simultaneously. We make the update gated and step-dependent. Writing the coupling term as $\boldsymbol{c}^t = \boldsymbol{x}^t\widetilde{\boldsymbol{J}}+\boldsymbol{z}^t A^\top$ (the $\boldsymbol{z}$ term absent in architecture (i)) and the observation as $\boldsymbol{x}^0$,

$$\boxed{\boldsymbol{p}^t = \boldsymbol{h} + \big(\mathbf{1} + \boldsymbol{\phi}_t\big) \odot \boldsymbol{c}^t + \boldsymbol{\sigma}_t \odot \boldsymbol{x}^0, \qquad \boldsymbol{x}^{t+1} = \tanh\big(\boldsymbol{p}^t/\tau_t\big),} \tag{S5}$$

with three learned, step-indexed gates: a *skip gate* $\boldsymbol{\sigma}_t = \alpha_t \mathbf{1} + \boldsymbol{w}_t \odot \boldsymbol{x}^0 + (\boldsymbol{x}^0 V_t) V_t^\top$ (rank $K_s$=32) that re-injects and trusts/distrusts the observation per gene and step; a *field gate* $\boldsymbol{\phi}_t$ (rank $K_f$=16) that rescales the coupling drive; and an *adaptive temperature* $\tau_t = \text{softplus}(a_t + b_t \rho)$ that depends on the fraction $\rho$ of genes present, annealing the relaxation with genome richness. Fields and gates are conditioned on a coarse summary of input module content through a small MLP over $M = 419$ functional modules, so the denoiser knows it is relaxing, e.g., a genome that already looks respiratory. The coupling term in (S5) is the analogue of a TAP/Onsager reaction correction to naive mean field (Thouless et al., 1977; Nguyen et al., 2017): the field gate $\boldsymbol{\phi}_t$ supplies the learned, step-indexed reaction term that the bare update of Eq. (S2) lacks. The gated higher-order model has $\sim$29M parameters at the production depth $T$=20 ($\sim$39M for the hidden-unit variant), of which $\boldsymbol{J}$ accounts for $\sim$23M.

#### S1.4 Training: objective, staged curriculum, and relaxation depth

**Noise model.** From each clean extant genome $\boldsymbol{x}^*$ a noisy observation $\boldsymbol{x}^0$ is synthesised by a false-negative channel (present→absent with per-gene probability $f_\text{N}$) and a false-positive channel (absent→present with probability $f_\text{P}$=0.01). During training $f_\text{N}$ is drawn per genome from a stage-dependent distribution of increasing difficulty (Table S1), from a gentle $0.9 \cdot \text{Beta}(1, 5)$ (mean $f_\text{N} \approx 0.15$) up to $0.9 \cdot \text{Beta}(5, 1)$, which concentrates mass in the deep-ancestral regime $f_\text{N} \in [0.75, 0.90]$, and a final uniform sweep over $[0, 0.9]$.

**Objective.** The models reported here are trained with an ELBO-style loss. (The code's default is a presence-weighted squared error, but every Ising and higher-order model we report uses the `elbo` loss below.) It sums a binary cross-entropy between the recovered presence probabilities $p_i = \frac{1}{2}(1+x_i^T)$ and the clean genome $y_i = \frac{1}{2}(1+x_i^*)$ with a Besag pseudolikelihood of the *clean* genome under the pairwise model,

$$\mathcal{L} = -\tfrac{1}{N} \textstyle\sum_i \big[y_i \log p_i + (1-y_i) \log(1-p_i)\big] \;+\; \alpha\, \tfrac{1}{N} \textstyle\sum_i \text{softplus}\big(-2\, x_i^* \,(h_i + \sum_j \widetilde{J}_{ij} x_j^*)\big), \qquad \text{(S6)}$$

with $\alpha$=0.3: the first term rewards accurate reconstruction, the second fits $\boldsymbol{h}, \widetilde{\boldsymbol{J}}$ directly to the pairwise statistics of real genomes, independent of the corruption. For the hidden-unit architecture (HO+hidden, Section S1.2) we add a *module-completeness auxiliary* at weight $\lambda$=0.3: a small head (linear–ReLU–linear–sigmoid) maps the final hidden state $\boldsymbol{z}^T$ to a predicted per-module present-fraction $\hat{\boldsymbol{f}} \in [0, 1]^M$, penalised by its mean-squared error against the true completeness of the *clean* genome, $f_m(\boldsymbol{x}) = |\mathcal{M}_m|^{-1} \sum_{i \in \mathcal{M}_m} \frac{1}{2}(x_i+1)$, over the same $M$=419 modules used for input conditioning (Section S1.3). This regulariser shapes $\boldsymbol{z}$ to encode module structure, and through it the coupling $\boldsymbol{J}$; the no-hidden architectures (i) and (iii) carry no hidden state for it to act on, so the production HO model and its fine-tunes are trained *without* it.

**A staged training curriculum.** The model is trained *de novo* (no pretrained checkpoint) by a curriculum that tempers two axes together — the difficulty of the corruption and the set of parameters left free to move (Table S1). Because the update is a deep, unrolled recurrence, we found this staging to make optimisation substantially more stable; the model is small ($\sim$ 29M parameters at $T$=20, $\sim$23M of them the coupling $\boldsymbol{J}$), so a flatter schedule may well converge too, but the numbers reported here are those of the curriculum below, which we did not re-run under a simpler schedule. The first three stages (**S1–S2b**) pre-train on a cheap surrogate, single-step *masked pseudolikelihood*: a fraction (15%) of genes is hidden and each is predicted from the rest through the model's one-step conditional $p(x_i \mid \boldsymbol{x}_{\setminus i})$, with no $T$-step relaxation. This warms up the couplings

| stage | objective | FN ($f_\mathrm{N}$/0.9) | lr (ep.) | newly trainable |
|---|---|---|---|---|
| S1 | masked PL | — | $10^{-3}$ (100) | $\boldsymbol{h}, \boldsymbol{J}$ (Ising core) |
| S2a | masked PL | — | $5\times10^{-4}$ (100) | $A, U, W$; conditioner+aux |
| S2b | masked PL | — | $3\times10^{-4}$ (200) | all parameters (joint) |
| S3a | denoising | Beta(1, 5) | $10^{-4}$ (200) | scalar skip gate |
| S3b | denoising | Beta(2, 3) | $8\times10^{-5}$ (150) | (as S3a) |
| S3c | denoising | Beta(2, 1) | $5\times10^{-5}$ (150) | diagonal gates $w, u, d$ |
| S3d | denoising | Beta(5, 1) | $3\times10^{-5}$ (150) | low-rank gates $V, P$ (full) |
| S3e | denoising | Uniform | $10^{-5}$ (100) | gates+conditioner; $\boldsymbol{J}$ frozen |

**Table S1: The eight-stage *de novo* training curriculum**, all run at $T$=8. Three masked-pseudolikelihood pre-training stages (S1–S2b) warm the parameters up in groups; five denoising stages (S3a–S3e) optimise Eq. (S6) while ramping the false-negative difficulty and switching on the gates. "newly unfrozen" lists the parameter groups added at that stage on top of those already trainable. These eight stages are the complete *de novo* curriculum; the relaxation depth is then extended $8 \to 20$, a third-order head is trained on top for architecture (iii), and three domain specialists are fine-tuned from the result — separate follow-on passes, not additional curriculum stages.

$\boldsymbol{h}, \boldsymbol{J}$ (the gates play no role in the one-step conditional and are first trained in S3) before the far more expensive denoising objective, and is where the parameters are released in groups: **S1** trains only the Ising core ($\boldsymbol{h}, \boldsymbol{J}$); **S2a** adds the hidden couplings $A, U, W$ and the module conditioner with its auxiliary head, holding ($\boldsymbol{h}, \boldsymbol{J}$) fixed (the adaptive-temperature parameters are trainable from S1 onward); **S2b** then trains all parameters jointly. The five denoising stages **S3a–S3e** optimise Eq. (S6) while ramping the false-negative difficulty from gentle (Beta(1, 5), mean $f_\mathrm{N} \approx 0.15$) to extreme (Beta(5, 1), mean $\approx 0.75$) and finally the full uniform spectrum, and while switching on the gates in order of expressiveness: the per-gene diagonal gates at S3c and the low-rank gates at S3d, reaching the complete model. The final stage **S3e** polishes the gates and conditioner at a low learning rate with $\boldsymbol{J}$ *frozen*, so the couplings settle under the milder regimes rather than being distorted by the hardest noise. Learning rate decreases monotonically across stages; each stage warms up over 5% of its steps and then cosine-anneals.

**Optimisation.** Every stage is optimised with AdamW (weight decay $10^{-4}$, applied to all groups including $\boldsymbol{J}$) and gradients clipped to global norm 1, in fp16 mixed precision (with TF32 enabled for the FP32 matrix-multiply paths such as the pseudolikelihood term). Per-GPU batches hold 8,000 genomes for the denoising stages and 2,048 for the masked-PL stages, and are scaled down with $T$ in the deepest passes to fit activation memory. Of the $K$=4 corrupted replicates drawn per genome one is left uncorrupted (clean fraction 0.125, floored to one replicate), so the model also sees clean-in/clean-out examples. Each stage runs a fixed epoch budget and hands its last-epoch weights to the next — there is no best-validation checkpoint selection or early stopping. During the depth-extension and higher-order passes the coupling $\boldsymbol{J}$ is optimised at 1/10 of the stage learning rate — and frozen outright in the bacterial and mixed fine-tunes — while the third-order attention head trains at 0.3×, so the pre-fit couplings are not distorted as new capacity is annealed in. Reported metrics are from single training runs (only the fixed validation-corruption sets are seeded).

**Relaxation depth $T$.** The depth $T$ — the number of mean-field sweeps — is not a gradient-learned scalar but is set during training by a short *depth curriculum*. The eight stages above are run at $T$=8; the trained model is then deepened in steps $8 \to 12 \to 16 \to 20$. At each extension the already-trained per-timestep parameters (the step-indexed gates $\boldsymbol{\sigma}_t, \boldsymbol{\phi}_t$ and temperature $\tau_t$) are kept for the first $T_{\text{old}}$ iterations, and the newly added iterations $t \in [T_{\text{old}}, T_{\text{new}})$ are initialised by interpolating from the last trained step toward *neutral*, identity-like values ($\alpha$=1, all gate weights 0, temperature $\tau$=1),

$$\text{param}_t = (1-\rho_t)\,\text{param}_{T_{\text{old}}-1} + \rho_t \cdot \text{neutral}, \qquad \rho_t = \frac{t - T_{\text{old}} + 1}{T_{\text{new}} - T_{\text{old}} + 1},$$

so a freshly deepened model reproduces its shallower predecessor and the extra sweeps begin as near-identity refinements; a brief fine-tune then adapts them. We take $T$=20 as the production depth: validation recovery improves with depth up to ∼16–20 and then plateaus. This progressive deepening is what the main text means by $T$ being determined during training — $T$ itself is a discrete hyperparameter, while the per-step dynamics that exploit the chosen depth are learned.

**Higher-order head.** For the production architecture (iii) the third-order attention head (Section S1.2) is attached on top of the depth-20 pairwise-plus-gates model and trained in a short additional pass. Its output projection is zero-initialised, so training begins from the *exact* pairwise model ($\Delta \equiv 0$) and the head is then annealed in (attention alone; then attention together with the gates and conditioner, with $\boldsymbol{J}$ frozen). It contributes a small, monotone recovery gain, largest on the noisiest genomes.

**Data, splits, and specialists.** Training uses the 113,104 species-representative genomes of GTDB r220 (Parks et al., 2022) (one genome per named prokaryotic species; 107,235 bacterial, 5,869 archaeal) in a 10-way split that holds out *whole phyla* (validation fraction 0.2 by genome count, so each model trains on ∼90,000), with $K$=4 independently corrupted replicates per genome, data-parallel on up to 8 A100 (80 GB) GPUs (eight for the *de novo* pretraining, four per higher-order or fine-tuning job). From the depth-20 no-hidden higher-order generalist we fine-tune three *domain specialists* under the same schedule: a mixed-domain rebalance (mix-FT), an archaea-only (arc-FT), and a bacteria-only (bac-FT) model, each re-split within its domain by whole phyla. The bacterial specialist subsamples to ∼15,000 training / ∼3,000 validation genomes; the archaeal specialist uses all ∼5,900 archaea and the mixed specialist ∼10,000 genomes balanced across domains. The denoising machinery is the same iterative relaxation that powers modern generative denoisers (Ho et al., 2020), here over an explicit, interpretable distribution over gene content.

### S2 Whole-phylum recovery stratified by phylum

To check that the headline recovery and calibration numbers are not carried by a handful of well-sampled lineages or by genome quality, we recompute both metrics within strata. For every held-out validation genome across the ten whole-phylum folds we corrupt the clean profile at false-negative rate $\text{fn} \in \{0.5, 0.75, 0.9\}$ (fixed $\text{fp} = 0.01$), denoise it through that fold's generalist HO-$T$=20 model, and pool the gene-level confusion counts and the 20-bin reliability histogram within each stratum before computing the Matthews correlation coefficient (MCC) and the expected calibration error (ECE) (∼ 245,000 genome evaluations in total). Whole-phylum hold-out means each phylum in Table S3 is scored by a model that never saw a single genome of it in training.

Recovery is robust to genome incompleteness (Table S2): even genomes below 80% CheckM completeness recover at MCC = 0.73 (fn = 0.5) to 0.59 (fn = 0.9), rising only modestly to 0.79/0.65 for near-complete genomes; the most-complete bin (99–100%) is marginally *worse* and slightly less calibrated (ECE = 0.068 at fn = 0.9), consistent with its larger accessory-gene load. Per phylum (Table S3) the spread is tight across all 105 phyla with ≥ 30 held-out genomes (88 bacterial, 17 archaeal): archaea – the LACA regime – recover as well as bacteria, while the single largest phylum, Pseudomonadota, sits at the low end (MCC = 0.60 at fn = 0.9, ECE up to 0.077), as expected for the clade with the most expansive accessory genome.

**Table S2:** Recovery (MCC) and calibration (ECE) stratified by CheckM genome completeness, pooled over held-out validation genomes at false-negative rates fn = 0.5/0.75/0.9 (fixed fp = 0.01), generalist HO-$T$=20. $n$ = genomes in bin (summed across CV splits).

| **Completeness** | $n$ | MCC | | | ECE | | |
|---|---|---|---|---|---|---|---|
| | | 0.5 | 0.75 | 0.9 | 0.5 | 0.75 | 0.9 |
| 0-80% | 51,181 | 0.731 | 0.654 | 0.587 | 0.032 | 0.040 | 0.040 |
| 80-90% | 44,171 | 0.773 | 0.698 | 0.634 | 0.026 | 0.033 | 0.041 |
| 90-95% | 38,596 | 0.788 | 0.714 | 0.651 | 0.025 | 0.031 | 0.041 |
| 95-99% | 53,022 | 0.789 | 0.714 | 0.651 | 0.028 | 0.036 | 0.049 |
| 99-100% | 57,673 | 0.776 | 0.696 | 0.632 | 0.039 | 0.052 | 0.068 |

**Table S3:** Recovery (MCC) and calibration (ECE) stratified by GTDB phylum, pooled over held-out validation genomes at fn = 0.5/0.75/0.9 (fixed fp = 0.01), generalist HO-$T$=20. Whole-phylum holdout: each phylum is scored by a model that never trained on it. size = mean present COGs per genome; Recall = TP/(TP+FN) and Prec. (precision) = TP/(TP+FP) of the recovered gene content at fn = 0.75. Phyla with $< 30$ genomes omitted (105 phyla). $n$ summed across CV splits.

| **Phylum** | $n$ | size | Recall | Prec. | MCC | | | ECE | | |
|---|---|---|---|---|---|---|---|---|---|---|
| | | | 0.75 | 0.75 | 0.5 | 0.75 | 0.9 | 0.5 | 0.75 | 0.9 |
| *Bacteria* | | | | | | | | | | |
| Pseudomonadota | 55,928 | 1,522 | 0.728 | 0.805 | 0.750 | 0.664 | 0.601 | 0.047 | 0.062 | 0.077 |
| Actinomycetota | 35,208 | 1,374 | 0.762 | 0.806 | 0.779 | 0.701 | 0.637 | 0.032 | 0.041 | 0.053 |
| Bacteroidota | 29,574 | 1,201 | 0.776 | 0.786 | 0.783 | 0.708 | 0.644 | 0.025 | 0.032 | 0.040 |
| Bacillota_A | 14,790 | 1,049 | 0.780 | 0.783 | 0.792 | 0.720 | 0.657 | 0.021 | 0.029 | 0.036 |
| Patescibacteria | 13,743 | 435 | 0.689 | 0.640 | 0.714 | 0.629 | 0.561 | 0.023 | 0.029 | 0.021 |
| Desulfobacterota | 11,140 | 1,352 | 0.784 | 0.821 | 0.794 | 0.727 | 0.667 | 0.025 | 0.029 | 0.039 |
| Verrucomicrobiota | 9,816 | 1,183 | 0.776 | 0.799 | 0.789 | 0.719 | 0.658 | 0.023 | 0.028 | 0.036 |
| Chloroflexota | 8,247 | 1,183 | 0.768 | 0.765 | 0.768 | 0.690 | 0.626 | 0.029 | 0.038 | 0.046 |
| Bacillota | 7,736 | 1,449 | 0.746 | 0.853 | 0.794 | 0.718 | 0.650 | 0.032 | 0.043 | 0.060 |
| Cyanobacteriota | 6,642 | 1,221 | 0.747 | 0.774 | 0.761 | 0.681 | 0.617 | 0.034 | 0.043 | 0.053 |
| Acidobacteriota | 5,673 | 1,433 | 0.789 | 0.819 | 0.791 | 0.723 | 0.662 | 0.027 | 0.031 | 0.042 |
| Spirochaetota | 5,392 | 1,142 | 0.753 | 0.778 | 0.776 | 0.694 | 0.625 | 0.025 | 0.034 | 0.045 |
| Bacillota_I | 2,903 | 660 | 0.757 | 0.725 | 0.771 | 0.698 | 0.640 | 0.018 | 0.025 | 0.024 |
| Planctomycetota | 2,339 | 1,294 | 0.784 | 0.788 | 0.778 | 0.708 | 0.650 | 0.030 | 0.035 | 0.042 |
| Bdellovibrionota | 2,176 | 1,172 | 0.766 | 0.787 | 0.783 | 0.705 | 0.640 | 0.023 | 0.030 | 0.039 |
| Gemmatimonadota | 2,140 | 1,321 | 0.788 | 0.812 | 0.793 | 0.725 | 0.665 | 0.025 | 0.029 | 0.038 |
| Elusimicrobiota | 1,504 | 897 | 0.756 | 0.784 | 0.791 | 0.718 | 0.653 | 0.014 | 0.018 | 0.023 |
| Bacillota_B | 1,440 | 1,289 | 0.803 | 0.835 | 0.816 | 0.754 | 0.693 | 0.020 | 0.023 | 0.033 |
| Bacillota_C | 1,180 | 1,223 | 0.763 | 0.809 | 0.792 | 0.715 | 0.646 | 0.025 | 0.033 | 0.046 |
| Omnitrophota | 1,170 | 871 | 0.772 | 0.787 | 0.797 | 0.731 | 0.667 | 0.013 | 0.015 | 0.018 |
| Myxococcota | 878 | 1,546 | 0.775 | 0.821 | 0.780 | 0.705 | 0.642 | 0.032 | 0.040 | 0.056 |

*continued on next page*

| **Phylum (cont.)** | $n$ | size | Recall | Prec. | MCC | | | ECE | | |
|---|---|---|---|---|---|---|---|---|---|---|
| | | | 0.75 | 0.75 | 0.5 | 0.75 | 0.9 | 0.5 | 0.75 | 0.9 |
| Aquificota | 792 | 1,053 | 0.776 | 0.813 | 0.802 | 0.738 | 0.674 | 0.018 | 0.021 | 0.028 |
| Armatimonadota | 618 | 1,264 | 0.774 | 0.801 | 0.788 | 0.713 | 0.648 | 0.024 | 0.029 | 0.040 |
| Zixibacteria | 532 | 1,159 | 0.795 | 0.834 | 0.816 | 0.757 | 0.701 | 0.013 | 0.013 | 0.018 |
| Eremiobacterota | 501 | 1,209 | 0.758 | 0.785 | 0.775 | 0.696 | 0.634 | 0.026 | 0.033 | 0.043 |
| Bacillota_G | 498 | 1,259 | 0.783 | 0.841 | 0.811 | 0.748 | 0.682 | 0.018 | 0.022 | 0.034 |
| Desulfobacterota_B | 453 | 1,441 | 0.796 | 0.827 | 0.799 | 0.732 | 0.674 | 0.024 | 0.026 | 0.036 |
| Cloacimonadota | 429 | 998 | 0.766 | 0.794 | 0.792 | 0.724 | 0.659 | 0.016 | 0.019 | 0.025 |
| Bipolaricaulota | 424 | 879 | 0.756 | 0.709 | 0.750 | 0.669 | 0.592 | 0.026 | 0.035 | 0.036 |
| WOR-3 | 408 | 973 | 0.782 | 0.807 | 0.806 | 0.743 | 0.678 | 0.011 | 0.012 | 0.017 |
| Fibrobacterota | 388 | 1,178 | 0.759 | 0.812 | 0.794 | 0.718 | 0.653 | 0.022 | 0.028 | 0.039 |
| Dependentiae | 369 | 448 | 0.663 | 0.705 | 0.735 | 0.652 | 0.583 | 0.012 | 0.018 | 0.021 |
| Chlamydiota | 311 | 789 | 0.675 | 0.750 | 0.753 | 0.658 | 0.583 | 0.018 | 0.028 | 0.039 |
| Myxococcota_A | 288 | 1,432 | 0.786 | 0.831 | 0.796 | 0.729 | 0.666 | 0.024 | 0.027 | 0.038 |
| Poribacteria | 282 | 1,323 | 0.790 | 0.803 | 0.787 | 0.720 | 0.655 | 0.026 | 0.030 | 0.039 |
| Marinisomatota | 275 | 942 | 0.779 | 0.760 | 0.782 | 0.712 | 0.652 | 0.020 | 0.025 | 0.026 |
| KSB1 | 248 | 1,467 | 0.793 | 0.842 | 0.804 | 0.740 | 0.683 | 0.023 | 0.026 | 0.038 |
| Bacillota_F | 240 | 1,346 | 0.785 | 0.841 | 0.810 | 0.743 | 0.683 | 0.021 | 0.026 | 0.037 |
| UBA10199 | 240 | 1,041 | 0.775 | 0.785 | 0.795 | 0.719 | 0.652 | 0.016 | 0.021 | 0.028 |
| Atribacterota | 220 | 1,008 | 0.774 | 0.759 | 0.774 | 0.703 | 0.637 | 0.023 | 0.026 | 0.031 |
| Deinococcota | 211 | 1,431 | 0.746 | 0.817 | 0.780 | 0.694 | 0.628 | 0.031 | 0.042 | 0.056 |
| Synergistota | 208 | 1,149 | 0.773 | 0.763 | 0.769 | 0.694 | 0.619 | 0.031 | 0.037 | 0.047 |
| Deferribacterota | 200 | 1,228 | 0.788 | 0.828 | 0.811 | 0.744 | 0.683 | 0.018 | 0.023 | 0.031 |
| Caldisericota | 196 | 835 | 0.746 | 0.741 | 0.767 | 0.689 | 0.613 | 0.020 | 0.026 | 0.030 |
| Desulfobacterota_G | 192 | 1,307 | 0.799 | 0.840 | 0.816 | 0.754 | 0.700 | 0.016 | 0.018 | 0.026 |
| SZUA-79 | 176 | 943 | 0.825 | 0.762 | 0.799 | 0.740 | 0.679 | 0.021 | 0.029 | 0.022 |
| Bacillota_E | 169 | 1,238 | 0.789 | 0.800 | 0.794 | 0.724 | 0.659 | 0.023 | 0.029 | 0.037 |
| Latescibacterota | 164 | 1,342 | 0.795 | 0.805 | 0.788 | 0.723 | 0.661 | 0.026 | 0.030 | 0.038 |
| Sumerlaeota | 160 | 1,271 | 0.784 | 0.814 | 0.795 | 0.728 | 0.666 | 0.021 | 0.025 | 0.034 |
| Desulfobacterota_E | 148 | 1,207 | 0.805 | 0.818 | 0.804 | 0.749 | 0.697 | 0.018 | 0.018 | 0.022 |
| Nitrospirota_A | 148 | 952 | 0.797 | 0.710 | 0.756 | 0.686 | 0.628 | 0.034 | 0.042 | 0.038 |
| Fusobacteriota | 138 | 1,123 | 0.757 | 0.784 | 0.787 | 0.702 | 0.632 | 0.024 | 0.036 | 0.046 |
| Electryoneota | 135 | 1,185 | 0.800 | 0.837 | 0.820 | 0.760 | 0.704 | 0.013 | 0.013 | 0.018 |
| SAR324 | 124 | 1,262 | 0.783 | 0.774 | 0.771 | 0.699 | 0.638 | 0.029 | 0.038 | 0.045 |
| Bacillota_D | 118 | 1,145 | 0.800 | 0.797 | 0.801 | 0.735 | 0.676 | 0.021 | 0.025 | 0.029 |
| Ratteibacteria | 117 | 859 | 0.795 | 0.768 | 0.796 | 0.733 | 0.674 | 0.014 | 0.019 | 0.016 |
| UBP6 | 112 | 986 | 0.761 | 0.794 | 0.795 | 0.721 | 0.648 | 0.013 | 0.018 | 0.027 |
| Bacillota_H | 108 | 1,171 | 0.809 | 0.821 | 0.818 | 0.755 | 0.702 | 0.017 | 0.020 | 0.024 |
| Thermotogota | 108 | 1,114 | 0.734 | 0.835 | 0.798 | 0.722 | 0.653 | 0.017 | 0.021 | 0.034 |
| Fermentibacterota | 100 | 1,078 | 0.771 | 0.814 | 0.800 | 0.734 | 0.670 | 0.015 | 0.017 | 0.022 |
| Krumholzibacteriota | 93 | 1,236 | 0.792 | 0.839 | 0.812 | 0.753 | 0.699 | 0.015 | 0.014 | 0.020 |
| Calditrichota | 92 | 1,417 | 0.801 | 0.845 | 0.811 | 0.751 | 0.696 | 0.021 | 0.022 | 0.032 |
| Methylomirabilota | 89 | 1,368 | 0.794 | 0.830 | 0.803 | 0.739 | 0.679 | 0.021 | 0.023 | 0.034 |
| Margulisbacteria | 87 | 848 | 0.753 | 0.759 | 0.780 | 0.704 | 0.639 | 0.017 | 0.023 | 0.027 |

*continued on next page*

| **Phylum (cont.)** | $n$ | size | Recall | Prec. | MCC | | | ECE | | |
|---|---|---|---|---|---|---|---|---|---|---|
| | | | 0.75 | 0.75 | 0.5 | 0.75 | 0.9 | 0.5 | 0.75 | 0.9 |
| Aerophobota | 75 | 867 | 0.780 | 0.733 | 0.772 | 0.701 | 0.630 | 0.022 | 0.026 | 0.026 |
| Riflebacteria | 70 | 1,241 | 0.742 | 0.771 | 0.758 | 0.673 | 0.610 | 0.032 | 0.041 | 0.051 |
| CSP1-3 | 68 | 1,215 | 0.745 | 0.821 | 0.791 | 0.713 | 0.647 | 0.020 | 0.025 | 0.036 |
| UBP14 | 68 | 941 | 0.804 | 0.823 | 0.822 | 0.769 | 0.711 | 0.008 | 0.009 | 0.007 |
| Delongbacteria | 63 | 1,113 | 0.784 | 0.812 | 0.798 | 0.739 | 0.677 | 0.019 | 0.019 | 0.024 |
| VGIX01 | 60 | 1,064 | 0.798 | 0.833 | 0.816 | 0.764 | 0.708 | 0.009 | 0.007 | 0.015 |
| Desulfobacterota_D | 55 | 990 | 0.767 | 0.795 | 0.798 | 0.725 | 0.666 | 0.016 | 0.021 | 0.027 |
| Campylobacterota_A | 52 | 1,066 | 0.788 | 0.824 | 0.816 | 0.752 | 0.687 | 0.014 | 0.016 | 0.024 |
| Eisenbacteria | 50 | 1,262 | 0.794 | 0.828 | 0.810 | 0.745 | 0.684 | 0.016 | 0.017 | 0.026 |
| BMS3Abin14 | 48 | 1,162 | 0.801 | 0.806 | 0.805 | 0.741 | 0.682 | 0.017 | 0.022 | 0.025 |
| Desulfobacterota_C | 48 | 1,470 | 0.794 | 0.862 | 0.815 | 0.755 | 0.699 | 0.021 | 0.024 | 0.034 |
| RBG-13-66-14 | 48 | 962 | 0.780 | 0.819 | 0.810 | 0.750 | 0.694 | 0.011 | 0.010 | 0.012 |
| UBA6262 | 45 | 944 | 0.783 | 0.850 | 0.828 | 0.773 | 0.713 | 0.007 | 0.005 | 0.013 |
| UBA9089 | 45 | 964 | 0.798 | 0.784 | 0.803 | 0.738 | 0.676 | 0.018 | 0.020 | 0.022 |
| Hinthialibacterota | 42 | 1,472 | 0.798 | 0.857 | 0.811 | 0.754 | 0.691 | 0.020 | 0.020 | 0.034 |
| JAAXVQ01 | 40 | 1,094 | 0.753 | 0.842 | 0.809 | 0.741 | 0.684 | 0.015 | 0.018 | 0.024 |
| QNDG01 | 39 | 1,307 | 0.798 | 0.847 | 0.819 | 0.758 | 0.703 | 0.014 | 0.015 | 0.025 |
| JAAXHH01 | 38 | 1,352 | 0.803 | 0.824 | 0.804 | 0.742 | 0.688 | 0.022 | 0.026 | 0.029 |
| UBA3054 | 36 | 925 | 0.779 | 0.750 | 0.776 | 0.706 | 0.642 | 0.021 | 0.028 | 0.030 |
| CG03 | 34 | 862 | 0.762 | 0.785 | 0.794 | 0.724 | 0.673 | 0.014 | 0.016 | 0.019 |
| JAGLYR01 | 32 | 1,130 | 0.785 | 0.838 | 0.811 | 0.756 | 0.706 | 0.013 | 0.012 | 0.015 |
| Tectomicrobia | 32 | 1,539 | 0.797 | 0.822 | 0.785 | 0.721 | 0.657 | 0.031 | 0.034 | 0.046 |
| Bdellovibrionota_G | 30 | 1,020 | 0.750 | 0.797 | 0.785 | 0.714 | 0.645 | 0.015 | 0.019 | 0.030 |
| SZUA-182 | 30 | 1,194 | 0.790 | 0.822 | 0.802 | 0.743 | 0.679 | 0.020 | 0.020 | 0.029 |
| *Archaea* | | | | | | | | | | |
| Thermoplasmatota | 3,192 | 758 | 0.722 | 0.717 | 0.745 | 0.666 | 0.601 | 0.019 | 0.030 | 0.033 |
| Thermoproteota | 2,528 | 786 | 0.761 | 0.731 | 0.762 | 0.695 | 0.635 | 0.022 | 0.029 | 0.028 |
| Nanoarchaeota | 1,694 | 447 | 0.631 | 0.738 | 0.731 | 0.653 | 0.581 | 0.006 | 0.010 | 0.015 |
| Halobacteriota | 1,223 | 1,109 | 0.692 | 0.836 | 0.768 | 0.698 | 0.640 | 0.034 | 0.044 | 0.058 |
| Micrarchaeota | 662 | 460 | 0.683 | 0.730 | 0.754 | 0.676 | 0.609 | 0.008 | 0.012 | 0.010 |
| Aenigmatarchaeota | 495 | 427 | 0.679 | 0.759 | 0.764 | 0.692 | 0.631 | 0.006 | 0.008 | 0.008 |
| Methanobacteriota | 488 | 1,017 | 0.768 | 0.799 | 0.783 | 0.726 | 0.681 | 0.026 | 0.034 | 0.036 |
| Methanobacteriota_B | 464 | 1,000 | 0.703 | 0.875 | 0.804 | 0.736 | 0.674 | 0.027 | 0.033 | 0.050 |
| Asgardarchaeota | 296 | 972 | 0.698 | 0.800 | 0.747 | 0.688 | 0.631 | 0.028 | 0.029 | 0.042 |
| Iainarchaeota | 202 | 486 | 0.665 | 0.788 | 0.772 | 0.696 | 0.623 | 0.003 | 0.004 | 0.011 |
| Altiarchaeota | 165 | 715 | 0.764 | 0.756 | 0.779 | 0.718 | 0.655 | 0.009 | 0.018 | 0.018 |
| Nanohaloarchaeota | 145 | 431 | 0.675 | 0.775 | 0.777 | 0.698 | 0.638 | 0.004 | 0.007 | 0.007 |
| Methanobacteriota_A | 105 | 995 | 0.775 | 0.841 | 0.817 | 0.760 | 0.703 | 0.018 | 0.024 | 0.031 |
| Hydrothermarchaeota | 92 | 915 | 0.779 | 0.790 | 0.791 | 0.734 | 0.679 | 0.016 | 0.021 | 0.026 |
| EX4484-52 | 68 | 503 | 0.704 | 0.750 | 0.767 | 0.695 | 0.620 | 0.004 | 0.010 | 0.013 |
| Hadarchaeota | 54 | 665 | 0.769 | 0.727 | 0.773 | 0.706 | 0.636 | 0.014 | 0.022 | 0.019 |
| SpSt-1190 | 50 | 493 | 0.687 | 0.812 | 0.783 | 0.721 | 0.665 | 0.003 | 0.003 | 0.008 |

### S3 Per-node model comparison and extant ground-truth recovery

#### S3.1 Per-node reconstruction models: specialist vs generalist

The architecture sweep summarised in the main text identifies the production models per node – mix-FT for the LACA, bac-FT for the LBCA – each scored on its own domain's whole-phylum hold-out against the generalist it was fine-tuned from (Table S4). Scored on the subset held out of *both* the root and the fine-tune training sets, the bacterial specialist bac-FT is level with the generalist on recovery (0.643 vs. 0.637 MCC at $f_{\mathrm{N}}$=0.9); its defensible advantage is robustness to common-gene (marginal-type) contamination, not higher recall.

**Table S4: Per-node reconstruction models: domain specialist vs generalist.** Recovery (MCC) and calibration (ECE) at fn = 0.5/0.75/0.9 (fixed fp = 0.01), each model scored on the whole-phylum hold-out for its domain. **bac-FT** is scored on genomes held out of *both* the root and the fine-tune training sets, where the bacterial specialist is level with the both-domain generalist HO-$T$=20 on recovery (0.643 vs. 0.637 MCC at $f_{\mathrm{N}}$=0.9); its defensible advantage is robustness to common-gene contamination (Section S4, Fig. S11), not higher recall. The bacterial generalist row pools the bacterial phyla of the both-domain hold-out (Table S3). *The most fine-tuned model, the marginal-FP HQ variant (Section S4), is shown for reference: against bac-FT it is at or slightly above the specialist on clean recovery (0.789/0.727/0.665 vs. 0.780/0.705/0.643) while adding the contamination resistance the specialists lack.

| **Model** | **Hold-out** | MCC | | | ECE | | |
|---|---|---|---|---|---|---|---|
| | | 0.5 | 0.75 | 0.9 | 0.5 | 0.75 | 0.9 |
| *Mixed-domain (LACA)* | | | | | | | |
| generalist HO-$T$=20 | mixed | 0.778 | 0.702 | 0.640 | 0.029 | 0.037 | 0.047 |
| **mix-FT** (LACA) | mixed | 0.786 | 0.715 | 0.657 | 0.023 | 0.028 | 0.036 |
| mix-FT-fp-marginal* | mixed | 0.772 | 0.714 | 0.662 | 0.023 | 0.026 | 0.032 |
| *Bacterial (LBCA)* | | | | | | | |
| generalist HO-$T$=20 | bacteria | 0.776 | 0.699 | 0.637 | 0.030 | 0.039 | 0.049 |
| **bac-FT** (LBCA) | bacteria | 0.780 | 0.705 | 0.643 | 0.021 | 0.027 | 0.035 |
| bac-FT-fp-marginal* | bacteria | 0.789 | 0.727 | 0.665 | 0.026 | 0.031 | 0.039 |

#### S3.2 Ground-truth recovery on extant genomes

Before turning to ancestors – where there is no ground truth – we test the models directly on *known* genomes, one at a time, under the noise an ancestral input actually carries. A deep reconstruction does not fail by sprinkling in random rare genes; it fails by over-calling the genes that are *common* across genomes – the housekeeping families that look unremarkable anywhere. We therefore corrupt each held-out genome with *marginal-type* contamination: false negatives at $f_{\mathrm{N}} \in \{0.5, 0.75, 0.9\}$ (deletion), plus injected false positives at $f_{\mathrm{P}} = 0.01$ drawn in proportion to each family's cross-genome frequency (Section S4, Fig. S9) – the same kind of noise a reconciliation or a copy-number (Count/GLD) reconstruction produces at a deep node. We denoise the corrupted profile through the cross-validation split whose held-out *phylum* contains the organism. For a fine-tuned specialist a strictly never-seen test requires the phylum to be held out of *both* the root and the fine-tune pools (we use such a both-stage clean split for *E. coli* below – split 5 – not the harness's auto-selected split). We compare the un-fine-tuned generalist (HO-$T$=20) against the fully fine-tuned, contamination-trained *marginal* model (bac-FT-fp-marginal for bacteria, mix-FT-fp-marginal for archaea). Figures S2–S8 break the outcome down per functional category against the truth.

**Model organisms.** For *Escherichia coli* (a 2,178-COG genome), scored on split 5 (held out of both the root and fine-tune training sets), the generalist and the marginal model recover a near-identical fraction of the true genome under common-gene contamination: on clean recall the specialist is on par with the generalist (e.g. the generalist recovers 80/73/69% at $f_\mathrm{N} = 0.4/0.6/0.8$, the marginal model 79/71/66%; 74/67/57% at $f_\mathrm{N} = 0.5/0.75/0.9$; both arms are a single shared corruption draw, and over 500 draws the marginal model averages $77 \pm 1/70 \pm 1/62 \pm 2\%$ at $f_\mathrm{N} = 0.4/0.6/0.8$). The marginal model's real advantage is not higher recall but robustness to common-gene contamination: the marginal-HQ model holds recall ($57 \to 58\%$ from $f_\mathrm{P} = 0.01$ to $f_\mathrm{P} = 0.05$ at $f_\mathrm{N} = 0.9$), while the hard-FN specialist bac-FT under uniform contamination collapses ($59 \to 39\%$); and it strips far more of the injected common-gene junk. For *Methanocaldococcus jannaschii* (a 1,100-COG archaeon) the two are closer (Fig. S5): the generalist's bacteria-trained prior already treats much archaeal-absent content as suspect, so it removes a comparable share of contamination; the marginal model matches it (recall 83/76/73%, removed 73/73/73%) without the generalist's recall sag at moderate noise.

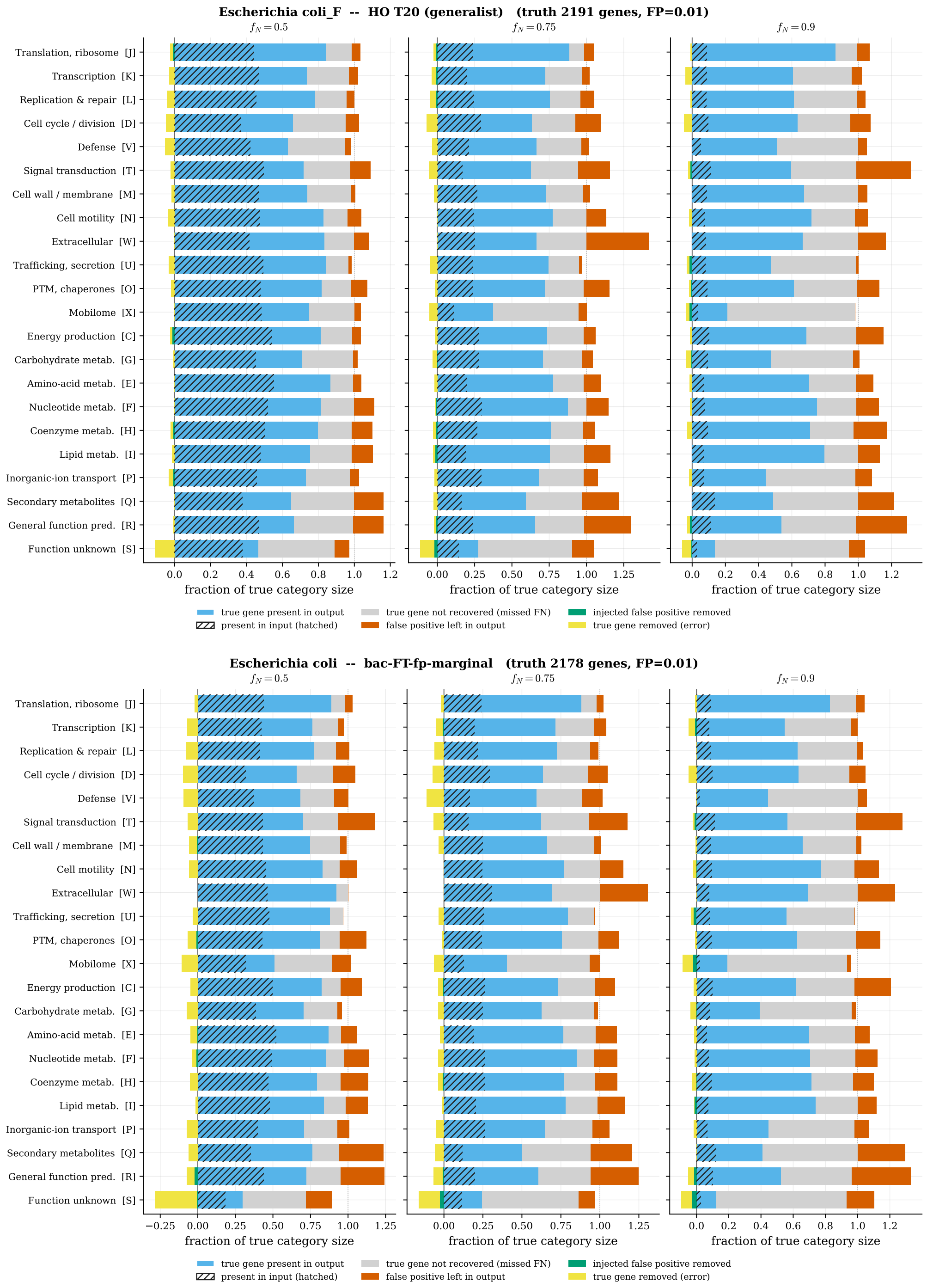


**Figure S2: Ground-truth recovery of *E. coli* under marginal-type contamination (split 5, held out of both the root and fine-tune training sets).** Per functional category, what the denoiser does to the corrupted genome at $f_{\rm N} = 0.5/0.75/0.9$ (columns; $f_{\rm P} = 0.01$ injected as common-gene *marginal-type* false positives, Section S4, Fig. S9) against the known truth, for the un-fine-tuned generalist HO-$T$=20 (top) and the fully fine-tuned marginal model bac-FT-fp-marginal (bottom). Bars are normalised to each category's true gene count (1.0 = the full true category); the hatched part was retained in the noisy input, the solid extension is what the denoiser *recovered*, a red segment beyond it is false positive left in the output, and bars left of zero are *removed* from the input (recall = recovered fraction, higher better). On this split the two models recover a near-identical fraction of the true genome (marginal-model recall 74/67/57% at $f_{\rm N} = 0.5/0.75/0.9$); the marginal model's advantage is contamination rejection – it strips far more of the spurious common genes and holds recall as $f_{\rm P}$ rises ($57 \rightarrow 58\%$ from $f_{\rm P} = 0.01$ to 0.05 at $f_{\rm N} = 0.9$) where the hard-FN specialist bac-FT under uniform contamination collapses ($59 \rightarrow 39\%$).

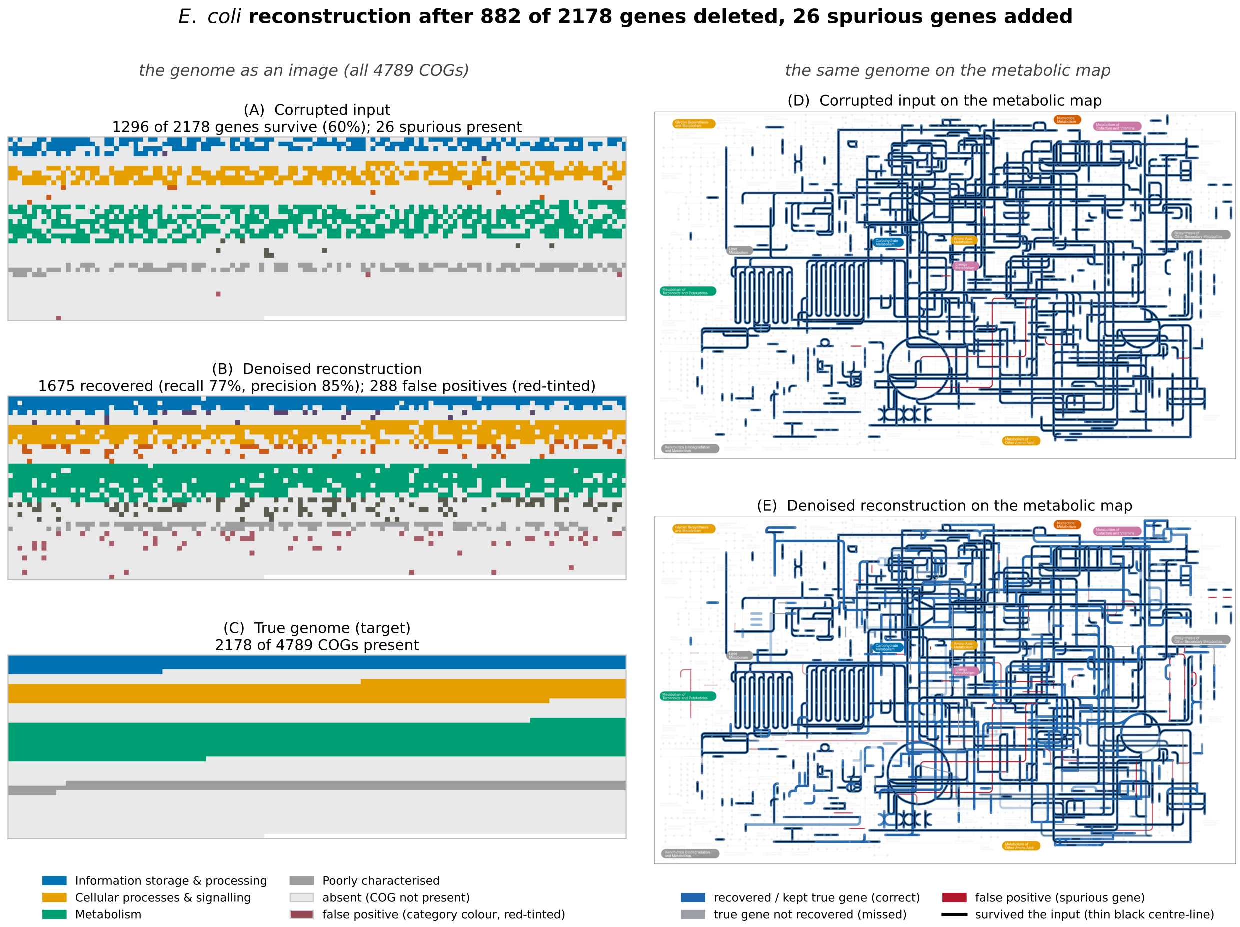


**Figure S3: Ground-truth recovery of *E. coli* at a moderate false-negative rate ($f_{\mathrm{N}} = 0.4$: 882 of 2,178 genes deleted, 26 common-gene false positives added; split 5).** The genome-as-image mosaic (left; true genes lead each functional-category group, the denoiser's false positives appended at the category's end in its own colour tinted red) beside the before/after iPath metabolic map (right; blue = correct gene, red = false positive). The production contamination-trained model recovers 77% of the genome at 85% precision. This is the milder-noise companion to the deep-ancestral $f_{\mathrm{N}} = 0.8$ figure in the main text.

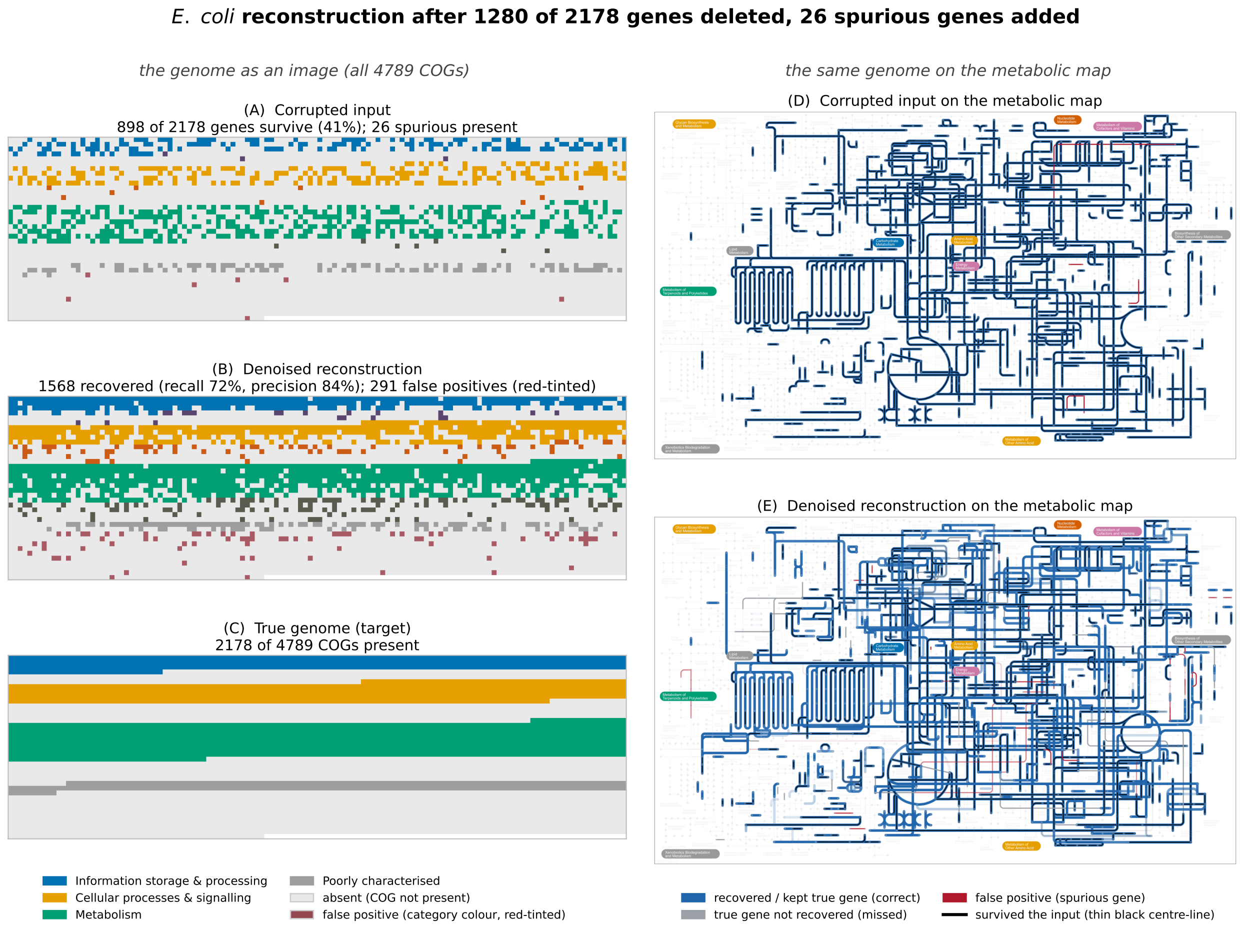


**Figure S4: Ground-truth recovery of *E. coli* at** $f_{\mathrm{N}} = 0.6$ (1,280 of 2,178 genes deleted, 26 added; split 5; layout and colours as Fig. S3). The denoiser rebuilds the genome to 72% recall at 84% precision.

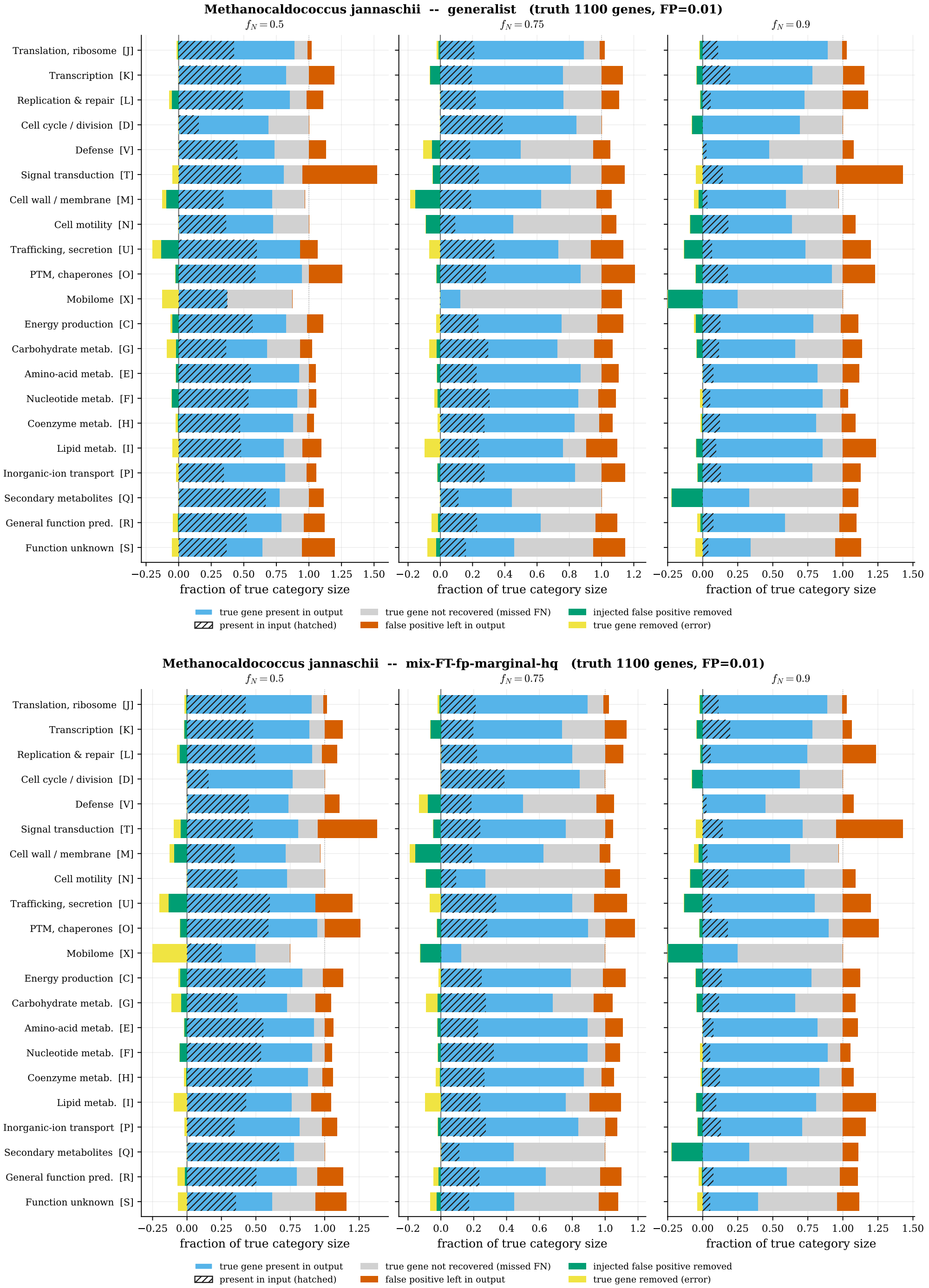


**Figure S5: Ground-truth recovery of *M. jannaschii* under marginal-type contamination (phylum held out at fine-tune).** Generalist HO-$T$=20 (top) and the marginal model mix-FT-fp-marginal (bottom), at $f_{\mathrm{N}} = 0.5/0.75/0.9$, $f_{\mathrm{P}} = 0.01$; bars and colours as in Fig. S2. On this archaeon the two are close on recall (marginal 83/76/73% vs generalist 82/75/73%) and on contamination removal (marginal 73/73/73% vs generalist 59/65/78%) – the generalist's bacteria-trained prior already treats much archaeal-absent content as suspect, so it removes a comparable share; the marginal model matches it without the recall sag at moderate noise.

**Median-size representatives.** We repeat the test on the *median-gene-count* held-out genome of each domain, one figure per organism comparing the generalist with the marginal model under the same marginal-type noise: *Kryptonium thompsonii* (a 1,263-COG bacterium of the deep-branching Ignavibacteria; Fig. S6) for bacteria. On the median bacterium recall is matched (marginal 84/78/82% at $f_{\mathrm{N}} = 0.5/0.75/0.9$ vs generalist 83/78/82%) but the marginal model removes far more of the contamination (46/43/40% vs 14/34/34%). The median-gene-count held-out archaeon, the uncultured `SM1-50 sp002506745` (754 COGs, 62 $\rightarrow$ 67% recall at $f_{\mathrm{N}} = 0.9$), is a 75.7%-complete medium-quality MAG of the ultra-divergent DHVEG-1 lineage, so its recall is a *lower bound* – which is why we re-select a high-completeness median archaeon. Re-selecting among *high-completeness* genomes ($\geq$ 90% CheckM, $\leq$ 5% contamination) gives the methanogen *Methanocorpusculum sp017387505* (90.5% complete, 930 COGs; Fig. S7, 77 $\rightarrow$ 78% recall) and the bacterium *UBA7675 sp002483085* (98.6% complete, 1,433 COGs; Fig. S8, recall matched at $\sim$76%, contamination removed lifted 29 $\rightarrow$ 35%). Across all four complete-to-near-complete medians the marginal model recovers as much of the true genome as the generalist and, on bacteria, removes two to three times as much contamination at moderate loss ($f_{\mathrm{N}} = 0.5$: 3.2× and 2.5×) and about a fifth more at higher loss – the behaviour an ancestral input demands.

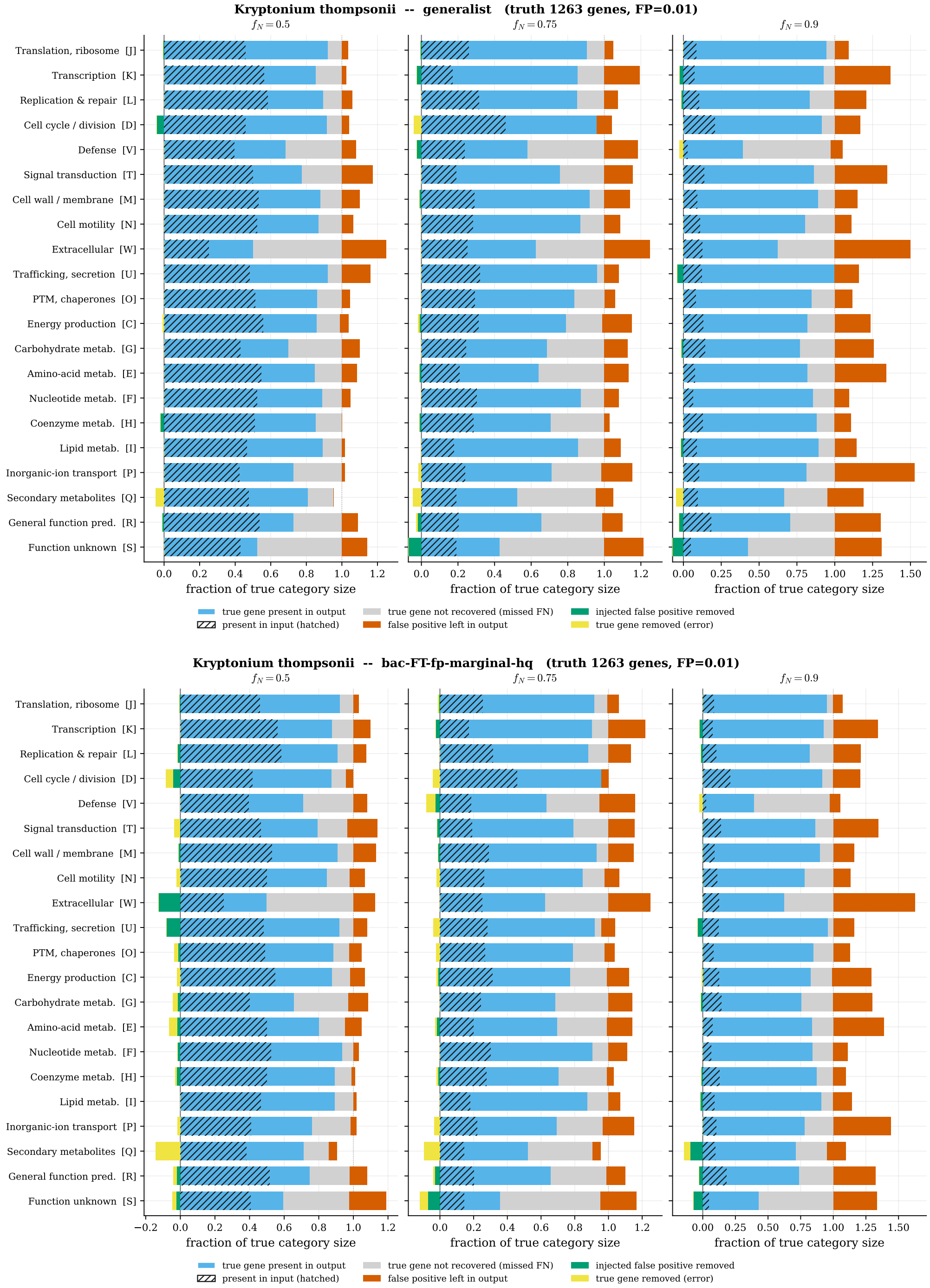


**Figure S6: Ground-truth recovery of *Kryptonium thompsonii* (median bacterium, 1,263 COGs; held out by phylum) under marginal-type contamination.** Generalist HO-$T$=20 (top) and the marginal model bac-FT-fp-marginal (bottom) at $f_{\mathrm{N}} = 0.5/0.75/0.9$, $f_{\mathrm{P}} = 0.01$; bars and colours as in Fig. S2. Recall is matched (marginal 84/78/82% vs generalist 83/78/82%) but the marginal model removes far more contamination (46/43/40% vs 14/34/34%) – the gain is in rejecting the spurious common genes, not in recall, on this already-well-recovered lineage.

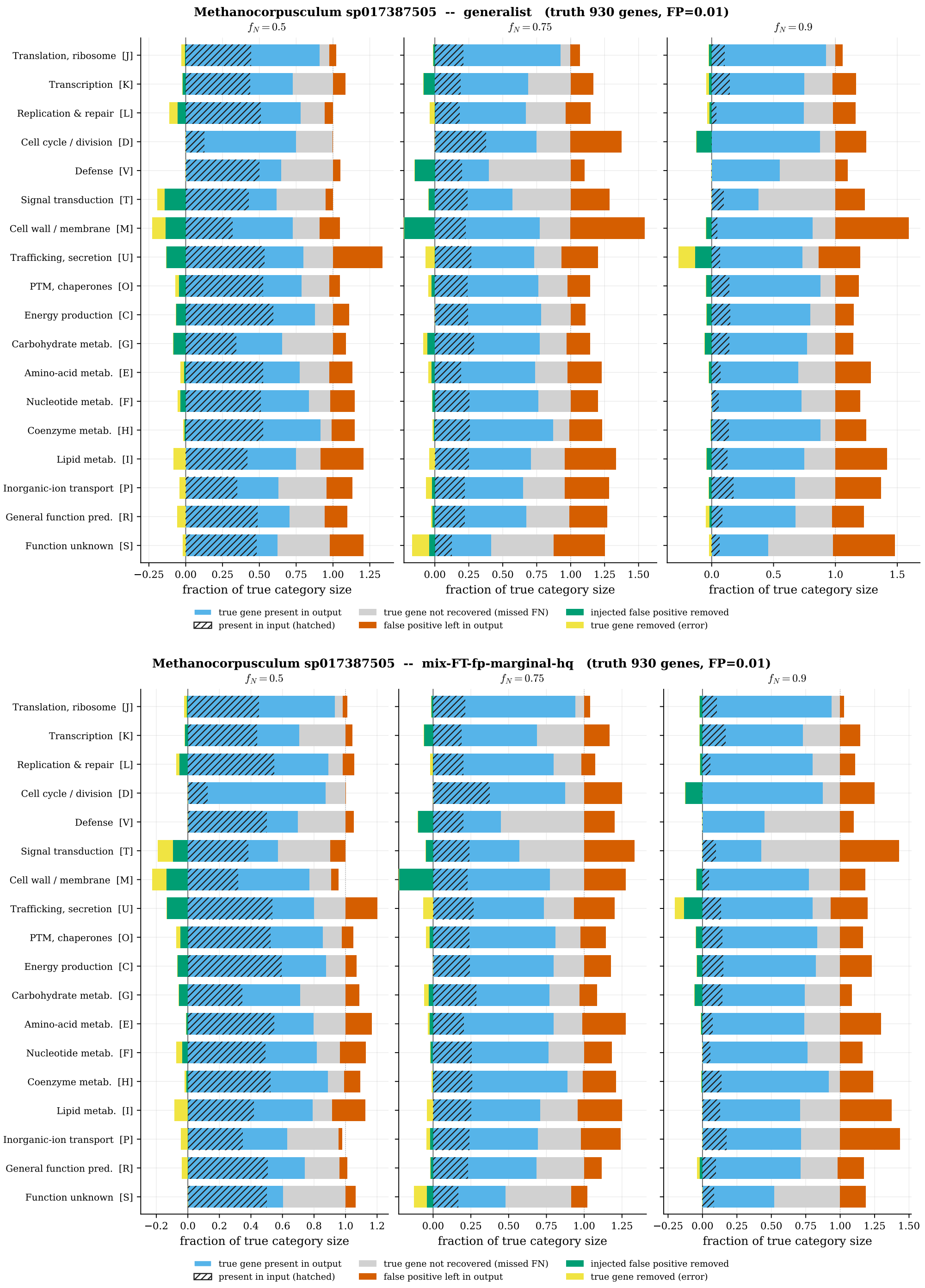


**Figure S7: Ground-truth recovery of a *high-completeness* median archaeon, *Methanocorpusculum sp017387505*** (90.5% CheckM completeness, 930 COGs, a Methanomicrobiales methanogen; held out by phylum; the high-quality counterpart to the lower-bound median archaeon SM1-50) under marginal-type contamination. Generalist HO-$T$=20 (top) and the marginal model mix-FT-fp-marginal (bottom); bars and colours as in Fig. S2. The marginal model recovers slightly more (80/78/78% vs 79/75/77%); here the bacteria-trained generalist is already aggressive on this archaeon and removes a few points more contamination (67/74/69% vs the marginal's 62/64/62%) – the one place the marginal model trades a little removal for recall.

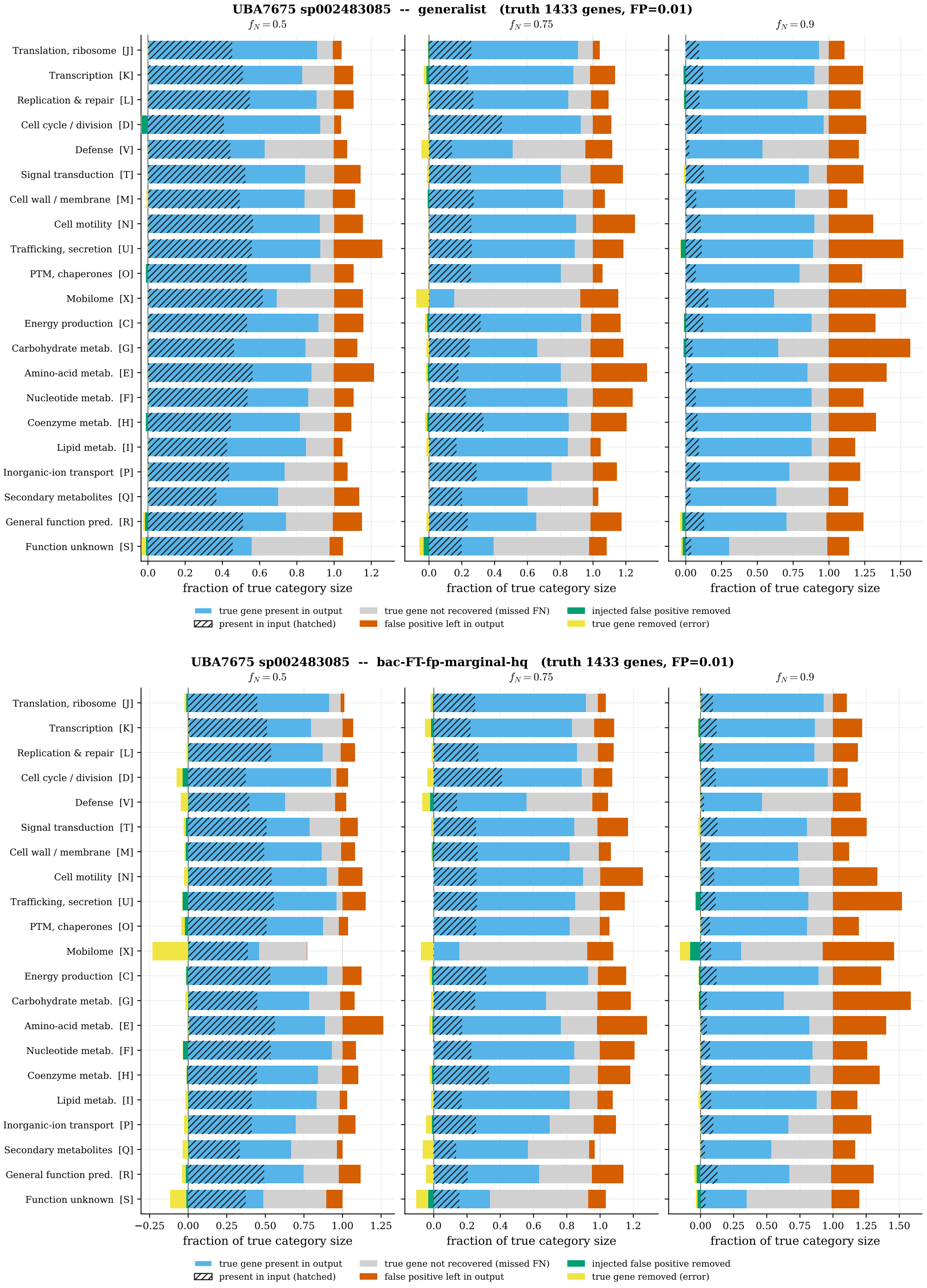


**Figure S8: Ground-truth recovery of a *high-completeness* median bacterium, *UBA7675 sp002483085*** (98.6% CheckM completeness, 1,433 COGs, a Bacteroidota/Kapaibacteriia genome; held out by phylum; the high-quality counterpart to Kryptonium, Fig. S6) under marginal-type contamination. Generalist HO-$T$=20 (top) and the marginal model bac-FT-fp-marginal (bottom); bars and colours as in Fig. S2. Recall is matched within a point (81/76/76% vs the generalist's 82/78/78%) while the marginal model again removes much more contamination (44/32/35% vs 18/26/29%) – the consistent bacterial pattern: the generalist leaves the spurious common genes in, the marginal model strips it.

**What the recovery test shows.** On genomes whose phylum was held out of training, the denoiser recovers 75–84% of the true gene content at $f_{\mathrm{N}} = 0.5$ and 57–82% at $f_{\mathrm{N}} = 0.9$, and removes 14–78% of injected false positives, concentrated in the function-unknown, mobilome and secondary-metabolite categories rather than in core machinery. The generalist-versus-marginal contrast is one of *robustness*, not recall: on clean recovery the two are on par, but the marginal model holds recall as the false-positive rate rises where the generalist collapses, and on bacteria it strips far more of the injected junk.

### S4 Realistic false-positive training and contamination rejection

#### S4.1 The marginal-FP model: deciding dense contamination

The FP-tolerant models are tuned against *sparse* coherent contamination – a few grafted foreign modules (the realistic curriculum of Section S4.2). They cannot adjudicate a *dense* input: a reconstruction that is already near-complete, whose extra breadth may be either real accessory content or aggregated noise. To give the denoiser that decision we fine-tune the HQ FP-tolerant models one step further with a *marginal-FP* curriculum. Alongside the clean-target and high-FN rescue copies, each genome gets prune copies whose absent COGs are re-injected as dense false positives sampled by their per-COG cross-genome marginal frequency, against the same clean target. The model thus reads input coherence and chooses, per genome, whether to trim or to fill.

Concretely, a prune copy is made with probability 0.5: it applies a low false-negative rate ($\leq 0.15$) and then injects $n_{\mathrm{add}} = \mathrm{frac} \cdot n_{\mathrm{true}}$ false positives ($\mathrm{frac} \leq 0.6$) drawn *without replacement* from the absent COGs by $p_c =$ the per-COG cross-genome marginal frequency. The clean genome is the unchanged target, so the model is trained to delete exactly this dense over-reconstruction. Because $p_c$ is heavy-tailed – a small near-universal core over a long rare tail (Fig. S9A) – the injection is dominated by *common* genes (mean injected $p_c = 0.53$ vs. 0.26 for a uniform draw; Fig. S9B): a broad but module-incoherent contamination that mimics an over-rich GLD ancestor, rather than the random rare junk of uniform-FP or the whole grafted modules of coherent-FP.

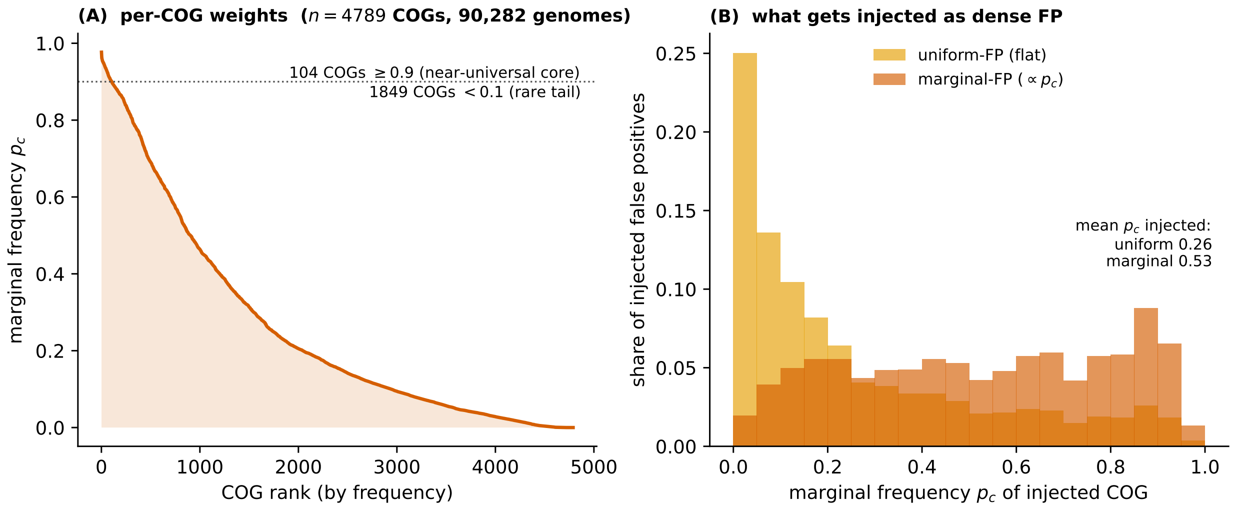


**Figure S9: The marginal-frequency dense-FP noise model.** (A) The per-COG cross-genome marginal frequency $p_c$ (fraction of the 90,482 training genomes carrying COG $c$), sorted: a near-universal core (114 COGs with $p_c \geq 0.9$) over a long rare tail (1,851 with $p_c < 0.1$). (B) The frequency composition of the injected false positives. Uniform-FP draws every absent COG equally, so it injects mostly rare genes (mean $p_c = 0.26$); the marginal-FP curriculum draws $\propto p_c$, concentrating the injection on common genes (mean $p_c = 0.53$) – a dense, plausible-looking but module-incoherent over-reconstruction the model is trained to trim back to the clean target.

On every benchmark the FP-tolerant models pass, the marginal-FP model matches them. On *random* per-COG FP it tracks bac-FT-fp throughout (Table S5, within 0.04 MCC at every rate). On the *coherent*-FP rejection frontier it sits with the FP-tolerant curriculum (Fig. S11). Under a 5% injection (∼215 spurious genes at the LBCA, ∼200 at the LACA) it resists inflation as well as bac-FT-fp (Table S7: LBCA +61, LACA +63, against default +175/+77). This is why the marginal model is the one we reconstruct the ancestors with: on the sparse reconciliation inputs the ancestral nodes actually present, it behaves as a rescuer, not a trimmer, yet it carries the contamination resistance the others lack, and its trimming engages only when an input is itself near-complete.

#### S4.2 False-positive-tolerant models and the coherent frontier

The reconstructions and the models' training assume a false-positive rate $f_P = 0.01$. The actual reconciliation inputs carry an apparent $f_P \approx 0.02$–$0.035$ – two to three times that, estimated by inverting the extant-recovery expansion curves. $f_P$ is the fraction of *truly absent* families wrongly scored present; a typical genome has ∼1,200 of its $N$=4,789 families present, so ∼3,600 are absent, and $f_P$ injects $f_P \times 3{,}600$ spurious genes – 36, 90, 180, 360 and 720 false positives at $f_P = 0.01$, 0.025, 0.05, 0.1 and 0.2.

We fine-tuned the two production models with a *realistic* false-positive curriculum, giving **mix-FT-fp** and **bac-FT-fp**. On top of a per-COG false-positive rate (rising to $f_P = 0.1$, Beta(1, 5)-weighted toward low rates) each training copy may also receive a block of *coherent* contamination – a whole injected absent module, or up to three modules grafted from a randomly chosen *other* training genome. This is the failure mode real reconciliation produces – lateral transfer or paralogue mis-assignment maps a neighbouring organism's module signature in as a block – which independent per-COG flips never present. The curriculum ladder is thus default → uniform-FP → realistic (coherent) FP → marginal (dense) FP.

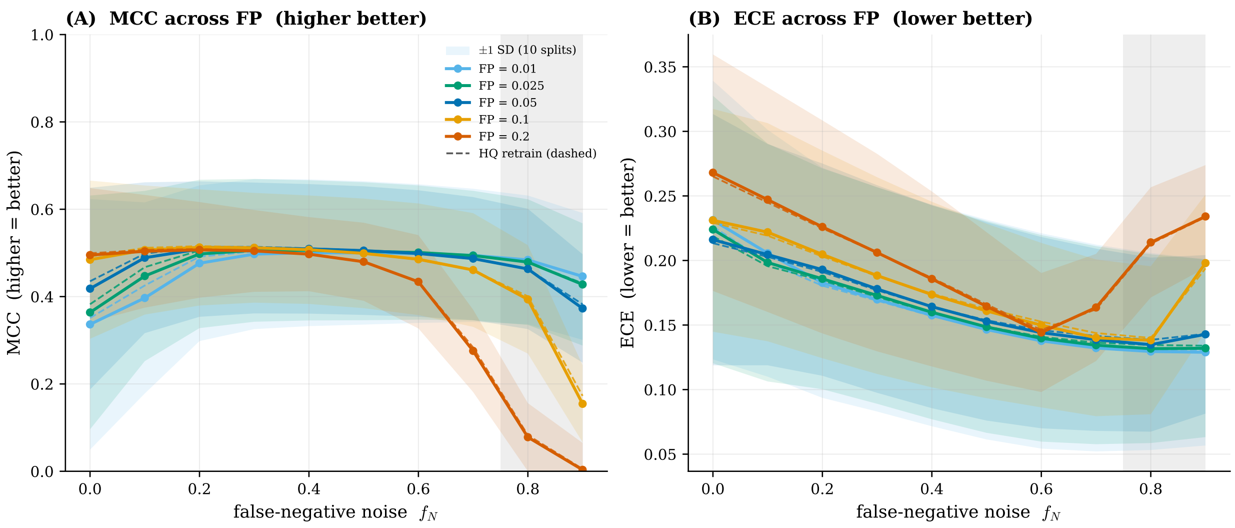


**Figure S10: The realistic FP-tolerant model is nearly flat across the realistic false-positive range – robust where the uniform-FP model was merely high.** Recovery (A, MCC, higher better) and calibration (B, ECE, lower better) of the realistic mix-FT-fp vs false-negative noise $f_{\mathrm{N}}$, one curve per injected false-positive rate $f_{\mathrm{P}}$; dashed curves are its high-completeness (HQ) retrain, which tracks within $\sim 0.02$ MCC. The vertical grey band marks the deep-ancestral regime; the coloured bands around each curve are $\pm 1$ standard deviation across the 10 phylum-holdout splits. The absent-by-default prior that lets this model reject *coherent* contamination (Fig. S11) comes at no cost in recovery – MCC peaks near 0.50, matching the parent mix-FT's 0.49 on the same reduction – and makes it almost *independent* of $f_{\mathrm{P}}$ through $f_{\mathrm{P}} \approx 0.05$: the $f_{\mathrm{P}} = 0.01/0.025/0.05$ curves overlap. It collapses only at $f_{\mathrm{P}} \geq 0.1$ in the deep-ancestral corner.

The realistic FP-tolerant model buys $f_{\mathrm{P}}$-robustness at no cost in recovery. At the deep-ancestral operating point ($f_{\mathrm{N}} = 0.9$) its MCC is 0.446 at $f_{\mathrm{P}} = 0.01$, matching mix-FT's 0.450 scored the same way, so the absent-by-default prior that suppresses *coherent* false positives costs it no recall. (The MCCs in this paragraph and in Fig. S10 are pooled over the annealing trajectory rather than read at the final step, so they are not comparable with the converged values quoted in the main text or in Table S5.) It is also remarkably flat in $f_{\mathrm{P}}$: at $f_{\mathrm{N}} = 0.9$ MCC holds at 0.428 ($f_{\mathrm{P}} = 0.025$) and 0.373 ($f_{\mathrm{P}} = 0.05$), and at the shallower noise of well-sampled lineages it is essentially $f_{\mathrm{P}}$-independent ($f_{\mathrm{N}} = 0.5$: MCC 0.500/0.504/0.505 at $f_{\mathrm{P}} = 0.01/0.025/0.05$). It collapses only at $f_{\mathrm{P}} \geq 0.1$ in the deep corner. Direct reconstruction confirms the spectra: injecting $f_{\mathrm{P}} = 0.05$ ($\sim$215 spurious genes at LBCA, $\sim$200 at LACA) grows the bac-FT-fp LBCA reconstruction only from 1,474 to 1,534 present families, and the mix-FT-fp LACA reconstruction from 1,014 to 1,083.

| $f_{\mathrm{P}}$ (random) | MCC (↑) | ECE (↓) |
|---|---|---|
| 0.01 | 0.674 | 0.035 |
| 0.025 | 0.637 | 0.046 |
| 0.05 | 0.555 | 0.075 |
| 0.1 | 0.264 | 0.173 |
| 0.2 | 0.106 | 0.196 |

**Table S5: Default specialist on random false positives** ($f_{\mathrm{N}} = 0.9$, bacteria, mean over ten phylum-holdout splits). Recovery (MCC) and calibration (ECE) as the random per-COG FP rate rises. The contamination-trained models are judged on their *target* noise – coherent grafted modules (Table S6, Fig. S11) – not on random flips, which every model strips.

| bacterial model (coherent FP) | recall (%) | FP removed (%) |
|---|---|---|
| default (bac-FT) | 68 | 8 |
| realistic-FP | 69 | 12 |
| marginal-FP | 70 | 13 |
| marginal + consistency | 70 | **17** |

**Table S6: Coherent contamination rejection, bacterial specialist** ($f_N = 0.9$, bacteria-only validation, at the standard call post > 0.5). Fraction of grafted foreign-module false positives removed. At matched recall (∼ 70%) the contamination curriculum doubles rejection (8 → 17%), most for the consistency fine-tune. The comparison is at *matched recall* across models, so the rejection ranking is robust to per-genome recall-level differences; the absolute recall column is on the auto-selected bac split. Full threshold sweep: Fig. S11.

Where it matters most – the ancestral present-count (Table S7) – the default model lets a 5% injection (∼ 215 spurious genes at LBCA, ∼ 200 at LACA) inflate LBCA by +175 families (1,508 → 1,683), whereas the FP-tolerant model limits it to +60 (1,474 → 1,534); LACA inflates +77 vs. +69. The FP-tolerant fine-tune is thus targeted insurance against contamination above the realistic rate – no measurable cost through $f_P = 0.05$, materially better false-positive rejection beyond it.

| | default | | FP-tolerant | | marginal-FP | |
|---|---|---|---|---|---|---|
| node | $f_P$=0 | $f_P$=0.05 | $f_P$=0 | $f_P$=0.05 | $f_P$=0 | $f_P$=0.05 |
| LBCA | 1,508 | 1,683 | 1,474 | 1,534 | 1,519 | 1,580 |
| LACA | 1,039 | 1,116 | 1,014 | 1,083 | 1,018 | 1,081 |

**Table S7: Ancestral present-counts, FP-tolerant and marginal-FP vs. default.** Families called present (posterior > 0.5) at the ancestral nodes, with no injection ($f_P$=0) and with 5% injected false positives (∼ 215 spurious genes at LBCA, ∼ 200 at LACA). The default model inflates the count under injection (+175 at LBCA); both the FP-tolerant (+60) and the marginal-FP HQ model (+61) resist it. On the clean reconciliation input the marginal-FP HQ model matches the realistic-FP HQ retrain to within one family (1,519 vs. 1,520 at LBCA; 1,018 at the combined-root LACA); its distinctive trimming only engages on *dense* contaminated inputs.

**The payoff: rejecting coherent contamination.** Random per-COG FP is the easy regime – a lone absent gene rarely fits a genome's couplings, so every model silences most of it. The false positives that actually survive reconciliation are functionally *coherent*: whole modules, or a foreign organism's module signature grafted in as a block. We test exactly this: delete present genes at the deep-ancestral rate, graft up to three modules from a random *other* held-out genome, denoise, and sweep the decision threshold, pooling recall against coherent-FP-removed over all ten splits. Coherent contamination is genuinely *hard* – a grafted module looks like a real one, so even the best model removes only a minority – but the realistic curriculum clearly *expands the precision/recall frontier* (Fig. S11): at matched recall it rejects ∼40% more coherent FP than the default (bacterial, $f_N = 0.9$: 11.8% vs. 8.2% removed at ∼67% recall; 23.6% vs. 16.1% at ∼48% recall), while the *uniform*-per-COG-FP curriculum barely moves it (9.0%). Restricting the realistic-FP fine-tune to high-completeness training genomes (≥90% CheckM) gives a small Pareto improvement rather than a trade-off: these HQ variants slightly *dominate* the standard realistic models on the coherent-contamination test (at $f_N$=0.9, $\tau$=0.5 they remove 12.0% at 69.1% recall vs. 11.8% at 67.2% for bacteria) and track them across the per-COG FP spectra.

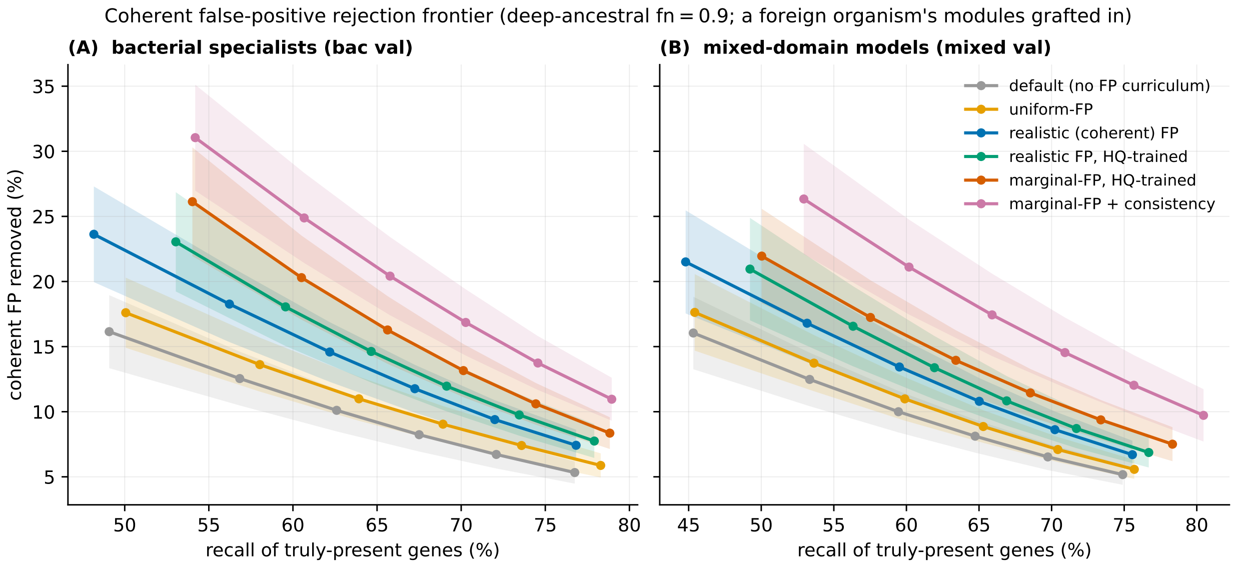


**Figure S11: The FP curricula expand the coherent-contamination rejection frontier, in ladder order.** Recall of truly-present genes vs. fraction of *coherent* false positives removed – a foreign organism's modules grafted into each held-out genome (up to three; deep-ancestral $f_{\mathrm{N}} = 0.9$) – swept over the decision threshold and pooled over the ten phylum-holdout splits (band: $\pm 1$ cross-split SD). (A) bacterial specialists on bacteria-only validation; (B) mixed-domain models on mixed validation. Each curriculum lifts the frontier in turn: the uniform-per-COG fine-tune (amber) barely improves on the default (grey); the realistic (coherent) curriculum (blue) and its high-completeness retrain (green) sit markedly higher, rejecting much more coherent contamination at matched recall; the **marginal-FP HQ** model (vermillion) is higher still; and the **marginal-FP + consistency** fine-tune (pink) sits highest of all in both domains. The ordering default < uniform < realistic/HQ < marginal < marginal+consistency mirrors the curriculum ladder. Coherent FP is hard (absolute removal stays well below the 55–80% of *random* FP), which is precisely why these are the false positives that survive reconciliation.

### S5 Genome-mass normalization of the coupling

Every reconstruction so far drives each gene's update with a coupling field $\sum_j \tilde{J}_{ij}\, s_j$: the pull of the company a gene keeps. That pull is not scale-free. It is a sum over the genes a genome carries, so its size grows with how full the genome is – in fluctuation, as $\sqrt{k}$ for a genome of $k$ genes. A gene-rich genome therefore feels a systematically stronger coupling drive than a gene-poor one, and the denoiser leans hardest on context exactly where the genome is already crowded. This is a nuisance for ancestral reconstruction, where the inputs are unusually gene-rich.

A second family of models removes the size bias with a single change: it rescales the coupling field by $\sqrt{\bar{k}/k}$, where $\bar{k} \approx 1{,}250$ is the mean gene count across genomes, holding the drive at the mean-genome scale. The intent is that a gene's reconstruction should depend on *which* company it keeps, not on *how much* of it the genome happens to contain. The field $h_i$, which sets each gene's base rate, is left untouched, so per-gene biases are unchanged; only the genome-to-gene pull is renormalized. Because the field is now *down*-weighted on full genomes, the prediction is concrete and falsifiable: on the gene-rich ancestral roots the normalization should *trim* the reconstruction, never inflate it, and the trim should be largest where the input is densest.

That is exactly what happens, and only just (Table S8). Matching ensembles split-for-split so the sole difference is the normalization, the reconstructed gene count moves by at most 32 genes, between 1.5% and 2.9% of the reconstruction, on every input. The direction follows the prediction

**Table S8: The genome-mass-normalized models reproduce every ancestral reconstruction to within** 3%. Matched split-for-split ensembles; "asserted" = input probability > 0.5, "present" = posterior > 0.5. *built* = genes called present that the input did not assert; *pruned* = asserted genes left absent. The normalization lifts only Euryroot, whose asserted count sits below the mean genome as the $\sqrt{\bar{k}/k}$ rescaling predicts, and trims the three gene-rich roots; the merged root is trimmed as well despite being sparser still.

| | | present | | | built / pruned | |
|---|---|---|---|---|---|---|
| input | asserted | cons | norm | Δ | cons | norm |
| LACA Euryroot | 1018 | 1345 | 1365 | +20 | 445/118 | 459/112 |
| LACA merged | 64 | 1019 | 990 | −29 | 956/1 | 927/1 |
| LACA GLD-min4 | 1422 | 1433 | 1406 | −27 | 180/169 | 182/198 |
| LACA GLD-min1 | 2017 | 1704 | 1672 | −32 | 113/426 | 118/463 |
| LBCA | 1408 | 1132 | 1113 | −19 | 58/334 | 60/355 |

on the gene-rich inputs: the three roots that assert more genes than the mean genome (LACA GLD-min1 2,017, GLD-min4 1,422, LBCA 1,408, against $\bar{k} \approx$ 1,250) are trimmed by 19–32 genes, while Euryroot (1,018 asserted) grows (+20), where $\sqrt{\bar{k}/k} > 1$ raises rather than lowers the drive. The merged root is the exception: it asserts only 64 families yet is trimmed as well (−29), its reconstruction being built almost entirely by the denoiser rather than carried over from the input. The input-to-output agreement (Jaccard) is unchanged to the third decimal, and the build-vs-prune balance shifts by only a handful of genes.

The build/prune columns also rank the inputs by the kind of error the denoiser judges them to carry. An input the model *prunes* heavily is one it reads as *over*-called (false-positive-prone); one it *builds* heavily it reads as *under*-called (false-negative-prone). The ordering is clean and unchanged by the normalization: the dense reconciliation roots LBCA (24% of asserted genes pruned) and LACA GLD-min1 (21%) are the most over-called; the soft combined and Euryroot inputs are the most under-called. So the five inputs span a false-positive-to-false-negative axis from the dense, over-called roots to the sparse, under-called ones, and the normalized models place them in the same order. Functionally the two model families are near-indistinguishable: the genes whose call flips between them are few (at most 56 pruned and 29 built on any input) and fall in the generic metabolic and poorly-characterized categories, not in any of the curated systems that define the reconstructions. The decisive archaeal signature of the LACA and the bacterial character of the LBCA are identical under both. Renormalizing the genome-to-gene pull thus does what it was meant to – it removes a genome-size bias and trims the densest inputs by the predicted handful of genes – while leaving the biological content of every ancestral reconstruction intact.

### S6 External validation detail

#### S6.1 Predicting protein interactions

The internal validation above measures recovery and calibration against held-out genomes corrupted by a known noise model. We now test, against an independent ground truth, whether the learned couplings $J_{ij}$ between gene families correspond to physical interactions between their protein products. If the model has captured biology rather than marginal gene frequencies, strongly-coupled families should interact more often than chance. This also sets our model beside a recent line of proteome-scale interactome predictors – the proteome language model ProteomeLM (Malbranke et al., 2026) and the deep-learning pathogen screen of Humphreys et al. (2024), which combines direct coupling analysis (DCA), RoseTTAFold2-Lite and AlphaFold. Our $J$ is the gene-content analogue of the coevolution those methods exploit: DCA reads residue–residue coevolution within paired alignments; $J$ reads gene–gene co-occurrence across genomes.

**Experimentally-validated interactions only.** We benchmark $J$ as those studies do – recovering protein–protein interactions (PPIs) annotated in STRING (Szklarczyk et al., 2023) – with one essential safeguard. STRING's `combined_score` aggregates evidence channels, one of which, *cooccurrence* (phylogenetic profiling), is exactly the signal $J$ encodes; *textmining* is likewise indirect. Scoring a co-occurrence predictor against them would be circular. We therefore take as ground truth STRING's *experimental* channel alone – interactions supported by physical and biochemical assays. (As a yardstick: STRING's own cooccurrence channel predicts the experimental channel at an area under the ROC curve (AUROC; 0.5 = chance, 1 = perfect) of 0.64.)

**Setup.** We took the 19 human bacterial pathogens of the Humphreys et al. (2024) screen (STRING v12.0; a WHO-priority/ESKAPE panel spanning Proteobacteria, Firmicutes and Actinobacteria). DIAMOND assigned each protein to a COG family against the NCBI COG-2020 reference (Galperin et al., 2021) (82–90% of every proteome mapped), giving it an index into $J$. Each of the 75.9 million intra-species protein pairs was scored by the coupling $J_{ij}$ of its two families and labelled positive if its STRING experimental score reached $\geq 700$ (high confidence) or $\geq 900$ (highest). We report AUROC, area under the precision–recall curve (AUPRC) and precision among the top-$N$ pairs. The coupling is the ten-split ensemble mean $J$ (the plain-pairwise and production higher-order matrices agree to three decimals, so the choice is immaterial), average-product-corrected (APC) as in coevolutionary contact prediction. Unlike the recovery and calibration benchmarks, which hold out whole phyla, this test uses the full ensemble coupling, so each species' own lineage is present in training: it asks what the couplings encode, not how the model generalises to unseen taxa, the same posture as sequence-based interactome predictors, including ProteomeLM, which score their full model.

**The couplings carry real interaction signal.** Across all 19 pathogens the coupling recovers experimentally-validated interactions well above chance (Fig. S12): pooled AUROC 0.62 at experimental $\geq 700$, rising to 0.65 for the highest-confidence interactions ($\geq 900$); per species, mean AUROC 0.63 ($\geq 700$, range 0.61–0.66) and 0.68 ($\geq 900$, reaching 0.89 for the best-resolved species), with pooled AUPRC 0.015 – 13.5× the random baseline ($1.1 \times 10^{-3}$). This matches STRING's purpose-built cooccurrence channel almost exactly (0.62 vs 0.64): a single matrix trained only to denoise gene content reproduces the phylogenetic-profiling PPI signal STRING computes by a dedicated pipeline.

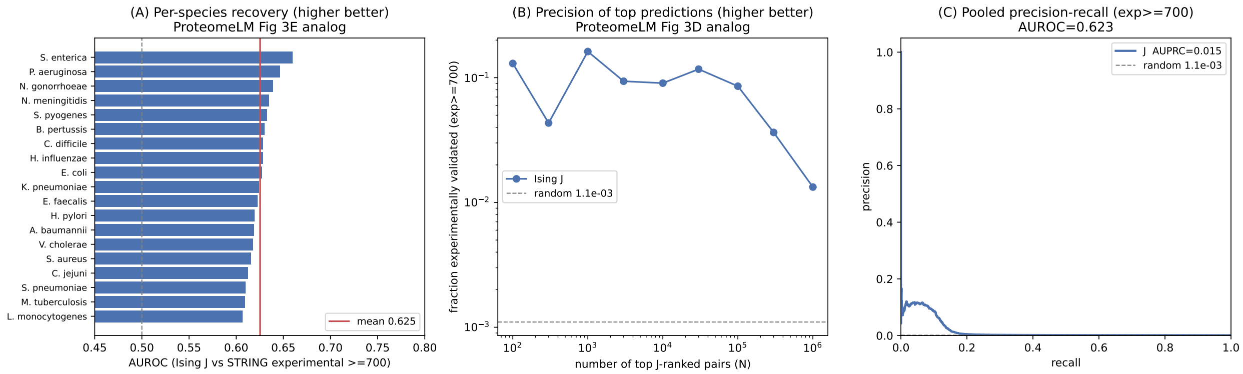


**Figure S12: The Ising coupling $J$ predicts experimentally-validated bacterial protein interactions.** Ground truth is STRING v12.0's *experimental* channel only (not `combined_score`, which embeds the cooccurrence signal $J$ is built from). (A) Per-species AUROC across the 19 human bacterial pathogens at experimental $\geq 700$ (higher is better, 0.5 = chance). (B) Fraction of the top-$N$ $J$-ranked pairs that are experimentally validated, vs $N$. (C) Pooled precision–recall curve and AUPRC, with the random baseline. $J$ is unsupervised, uses gene presence/absence only, and has seen no interaction data.

**Quantitative comparison with ProteomeLM.** The fair comparison places $J$ among unsupervised coevolution baselines, not the supervised state of the art (Table S9). On these 19 pathogens ProteomeLM's interactome screen reaches per-species AUC 0.87–0.92 against STRING high-confidence interactions, and on the human interactome it reaches 0.83 on the same physical-interaction target where direct-coupling analysis (DCA) reaches 0.73 (Humphreys et al., 2024; Malbranke et al., 2026). The most like-for-like point is ProteomeLM's *unsupervised* read-out, its raw attention coefficients; on its own benchmark these discriminate co-expressed and same-complex pairs at AUROC 0.79–0.95 but direct physical contacts only at 0.75–0.87, and with the protein sequence removed (an orthologous-group-identity encoding) the attention falls to 0.70 on *E. coli* direct interactions, so it is a sequence-aided functional-co-occurrence read-out of the kind $J$ encodes. Our coupling reaches AUROC 0.62 on the experimental channel, below all of these, as it must be: $J$ uses no protein sequence, only presence/absence; it predicts at family rather than protein resolution; and it is scored against the harder experimental-only target. Against STRING's combined-score target (the easier, partly-circular comparison for a co-occurrence method) the same $J$ rises to AUROC 0.69 ($\geq 700$) and 0.74 ($\geq 900$), within the range of the DCA coevolution baseline. The point is the lower bound: a single 4,789 $\times$ 4,789 coupling matrix, trained only to denoise gene content, carries interaction signal comparable to DCA and to STRING's co-occurrence channel (itself a phylogenetic-profiling pipeline), evidence that the couplings driving the ancestral reconstructions are biologically real.

**Table S9: Interactome recovery on the 19 human bacterial pathogens, in context.** AUROC against STRING v12.0. ProteomeLM and DCA numbers are as reported by Malbranke et al. (2026) and Humphreys et al. (2024), scored against STRING-derived physical interactions (ProteomeLM's interactome screen uses high-confidence STRING pairs; DCA is on the human interactome). ProteomeLM's unsupervised attention is shown on its strongest channel, co-expression; on direct physical contacts it scores lower (see text). Our $J$ is unsupervised and sequence-free; against the experimental channel (non-circular) it matches STRING's co-occurrence channel, and against the combined-score target (partly circular for a co-occurrence method) it matches the DCA coevolution baseline.

| method | input | supervision | STRING target | AU[illegible] |
|---|---|---|---|---|
| ProteomeLM screen (Malbranke et al., 2026) | proteome LM (sequence) | supervised | physical | 0.87[illegible] |
| ProteomeLM attention (Malbranke et al., 2026) | proteome LM (sequence) | unsupervised | coexpression | up to 0.9[illegible] |
| DCA (Humphreys et al., 2024) | residue coevolution | unsupervised | physical (human) | 0[illegible] |
| STRING co-occurrence channel | gene co-occurrence | – | experimental | 0[illegible] |
| **Ising** $J$ (this work) | gene presence/absence | unsupervised | experimental | 0[illegible] |
| **Ising** $J$ (this work) | gene presence/absence | unsupervised | combined | 0.69[illegible] |

**Robustness to training-set composition.** The AUROC reflects the learned couplings, not the lineages present in training. Holding each pathogen's whole GTDB phylum out of training and re-scoring against the held-out coupling leaves the experimental-channel AUROC essentially unchanged (mean change +0.003 at experimental $\geq 700$, relative to keeping the phylum in training). The comparison is defined for 17 of the 19 species; the two Campylobacterota (*H. pylori*, *C. jejuni*) have their phylum in training in all ten splits, so no held-out coupling exists for them. For *E. coli* a nested leave-clade-out series, holding out its phylum, then class, order, family, and the species itself, leaves the AUROC at 0.63 throughout, with the coupling matrices near-identical along the series (pairwise correlation above 0.999, since removing one clade barely shifts a co-occurrence statistic estimated over the whole sample). The interaction signal is thus a global property of the co-occurrence structure rather than memorised clade composition.

**Dynamic and higher-order couplings.** We also test whether the bare $J_{ij}$ is the best interaction read-out the model affords, scoring each COG pair by the denoiser's knockout *response* $R_{ij}(T)$, the drop in gene $j$'s denoised belief when gene $i$ is deleted from the species' own genome after $T$ relaxation sweeps, which folds in indirect ($i \to k \to j$) paths and the gates (Table S10). Across all 19 pathogens (plain pairwise model, response at its shallow optimum) the response improves on the static coupling for only the four best-resolved gammaproteobacteria (*E. coli* +0.031, *P. aeruginosa* +0.014, *S. enterica* +0.012, *K. pneumoniae* +0.011 in experimental AUROC) and is worse for the other 15, by 0.02 on average; the static $J$ is the better general read-out (mean experimental AUROC 0.627 versus 0.600 at $T$=2). The depth profile is consistent: $T$=1 under-propagates and by $T \gtrsim 5$ the genome relaxes back to its attractor and the single-gene signal washes out. Against STRING's combined target the static $J$ scores higher still (0.73 versus 0.61), so the response never beats it there. The higher-order head is no better: on the contamination-trained models the knockout response falls well below the static $J$ (experimental AUROC $\sim$0.53 versus 0.63), and reading the attention maps directly is *anti*-predictive (AUROC 0.42–0.47). Unlike ProteomeLM, where attention *is* the interaction read-out, in our architecture the pairwise interaction structure lives in the explicit coupling $J$, while the attention head carries higher-order denoising corrections orthogonal to physical interaction.

**Table S10: Static vs dynamic vs higher-order couplings.** Mean AUROC across the 19 pathogens (COG-pair level) against the experimental (non-circular) and combined STRING targets, for the static coupling $J$ and the knockout response $R(T{=}2)$ on the plain pairwise model; the higher-order row is the response on the contamination-trained models (*E. coli*). Bold marks the best read-out per target: the static $J$ wins both. ProteomeLM and DCA (combined target) are shown for reference.

| predictor | experimental AUROC | combined AUROC |
|---|---|---|
| static $J$ | **0.627** | **0.725** |
| dynamic $R(T{=}2)$, pairwise | 0.600 | 0.611 |
| dynamic $R(T{=}2)$, higher-order | 0.53 | 0.57 |
| HO attention map (ProteomeLM-style) | 0.45 | 0.45 |
| DCA (Humphreys et al., 2024) (ref.) | – | 0.73 |
| ProteomeLM (Malbranke et al., 2026) (ref.) | – | 0.87–0.92 |

**A supervised head reaches ProteomeLM's range, on a degree bias.** ProteomeLM combines its attention coefficients in a supervised classifier; the analogue here is a classifier over the explicit coupling $J$ together with the model's per-gene denoising confidence $c_i = 1/(1 - m_i^2)$, with the pair feature $c_i c_j$ ($m$ is the one-step reconstruction of the genome). On *E. coli* $c_i c_j$ predicts the experimental channel at AUROC 0.78 by itself, and a gradient-boosted classifier over $[J, c_i c_j]$ reaches 0.84, sequence-free and within range of ProteomeLM's supervised, sequence-based head (0.87–0.92). Across all 19 pathogens, scored in-distribution, the boosted head reaches mean AUROC 0.80 (range 0.75–0.84), $c_i c_j$ alone 0.74, and the explicit $J$ 0.62, so the supervised lift over $J$ is the degree feature on every species, not *E. coli* alone. The number is misleading. The feature $c_i c_j$ is rank one: it scores a pair by how strongly the model is committed to each of its two genes, so it can report only that each gene is interaction-prone, never that the pair interacts, and it predicts because interaction networks are dominated by a few high-degree genes. It is a per-gene degree bias, the documented failure mode of supervised interaction predictors (Bernett et al., 2024), not a coupling; the explicit $J$, which is pair-specific, scores 0.63. The head's apparent strength also inflates with how much of the target's lineage it has seen: across the leave-clade-out series around *E. coli* the boosted score rises from 0.827 (whole phylum held out) to 0.843 (only the species held out), $c_i c_j$ from 0.774 to 0.799, and a linear head's weight on $c_i c_j$ from 0.09 to 0.27, with the weight on $J$ unchanged. The same node bias and in-distribution sharpening operate in any supervised attention read-out, ProteomeLM's included, whose interactome is scored with the target lineages in training and has not been tested under lineage hold-out. The whole-phylum hold-out is what separates a transferable coupling from a degree bias that inflates in-distribution, and it leaves $J$ at 0.63 throughout.

#### S6.2 Gene essentiality

ProteomeLM also predicts gene *essentiality*, and the denoiser affords a direct unsupervised analogue. A gene whose presence the rest of the genome strongly re-implies should be functionally central, so we measure each gene's *leave-one-out reconstructability* – delete it from the *E. coli* genome, denoise, and read the model's posterior that it is present – and benchmark against the experimental essentiality calls of Goodall et al. (2018) (358 essential vs 3,793 non-essential genes; 2,026 map to a COG present in *E. coli*, 16% essential).

The reconstructability predicts essentiality above chance, and beyond mere conservation: AUROC 0.64 for the production higher-order model, with area under the precision-recall curve (AUPRC, whose baseline equals the positive-class prevalence) 0.22 against a prevalence of 0.16.

This is not a conservation artefact. The model's single-site field $h_i$ (its learned base rate) is *anti*-predictive of *E. coli* essentiality (AUROC 0.38), because the field is pan-prokaryotic whereas essentiality is species-specific, and the signal survives residualising on $h$ (AUROC 0.62). The context-dependent reconstructability, not raw gene frequency, carries the signal. As with interactions, this unsupervised gene-content signal is real but modest, well below the supervised, sequence-based ProteomeLM, whose essentiality classifier reaches AUC ∼0.93 across taxa and 0.95 on *E. coli* (Malbranke et al., 2026). The contrast with the interactome is informative: there the higher-order head added nothing, whereas here it helps, because essentiality reflects a gene's embedding in the whole functional network, the beyond-pairwise structure that head captures.

**A supervised read-out from the relaxation trajectory.** The posterior above uses only the endpoint of the relaxation; its whole course carries more, namely how fast and how strongly the rest of the genome re-implies a deleted gene at each iteration. We fit a gradient-boosted classifier to each gene's reconstruction $x_i^1, \ldots, x_i^T$ under its own deletion, together with the field $h_i$ and the gene's coupling degree ($\sum_j |J_{ij}|$, $\sum_j J_{ij}^2$), benchmarked against Goodall et al. (2018) by five-fold cross-validation over genes (no gene in both train and test). It reaches AUROC 0.80, against 0.64–0.74 for the endpoint posterior alone and still using no sequence, approaching the supervised, sequence-based ProteomeLM-Ess (0.93 across taxa, 0.95 on *E. coli*); the remaining gap is attributable to sequence rather than to the read-out. Across the leave-clade-out series around *E. coli* the trajectory classifier is flat (0.79–0.81 at every rung from whole-phylum to species hold-out), whereas the endpoint posterior varies non-monotonically (0.70–0.74, peaking at the intermediate rung and returning to its whole-phylum value once only the species is held out): the trajectory captures the transferable essentiality structure, while the endpoint is the noisier read-out.

### S7 Conservative and generous reconstructions at the LBCA

Two established ways of reconstructing an ancestral gene content disagree sharply at the bacterial root. Probabilistic gene-tree/species-tree *reconciliation* is cautious: at the last bacterial common ancestor (LBCA) it confidently places almost nothing – only 6 families reach a posterior of 0.5, and only ∼ 493 carry any appreciable signal at all (the *sparse* input). A *profile-level* gain-loss-duplication reconstruction (GLD – the Count family of methods, here recount), which models gene gains, losses, and duplications from presence/absence *profiles* alone with no gene-tree information, is generous: it returns a *dense* ancestor of ∼ 2,000 confidently present families. The catch is that a profile fit, taken family by family, never checks whether those families form a coherent organism. We answer the organism-level question by feeding each reconstruction to the denoiser and seeing what it keeps.

**Build or prune.** Handed the two inputs, the same model moves in opposite directions (Fig. S13, Table S11). Fed the *dense* GLD ancestor, it acts as a **pruner**: it judges ∼ 20% of the confidently-present input to be false and switches it off (394 families at the min1 cut, 391 at min4), adding only a couple of hundred. Fed the *sparse* reconciliation, it acts as a **builder**: it keeps 374 of the 493 families the reconciliation gave any support, and adds ∼ 1,150 more that the reconciliation had left at zero – assembling a genome where there was barely one. It does decline 119 of the 493 supported families, but those are the weakly-supported ones: their median reconciliation posterior is just 0.01, only 6 of them reached even 0.1 and just 1 reached 0.2 (Fig. S13B). The model keeps essentially everything the reconciliation actually stood behind, declines only what it barely suggested, and supplies the rest itself.

| reconstruction | input present | output | added | removed | net |
|---|---|---|---|---|---|
| sparse reconciliation ($x = p$) | 493[†] | 1,519 | 1,145 | 119[‡] | +1,026 (build) |
| GLD min1 | 2,014 | 1,836 | 216 | 394 (20%) | −178 (prune) |
| GLD min4 | 1,981 | 1,776 | 186 | 391 (20%) | −205 (prune) |

**Table S11: Genes added and removed relative to each reconstruction's own input.** "added" = present in the denoised output but not the input (rescued losses); "removed" = present in the input but switched off (rejected gains). [†]Families with any reconciliation support; only 6 reach posterior 0.5 (four strictly above). [‡]The 119 declined families were weakly supported to begin with – median input posterior 0.01, only 6 above 0.1 and 1 above 0.2 (Fig. S13B) – so the model keeps what the reconciliation backed and declines what it barely raised. The GLD inputs binarise at input probability $> 0.5$.

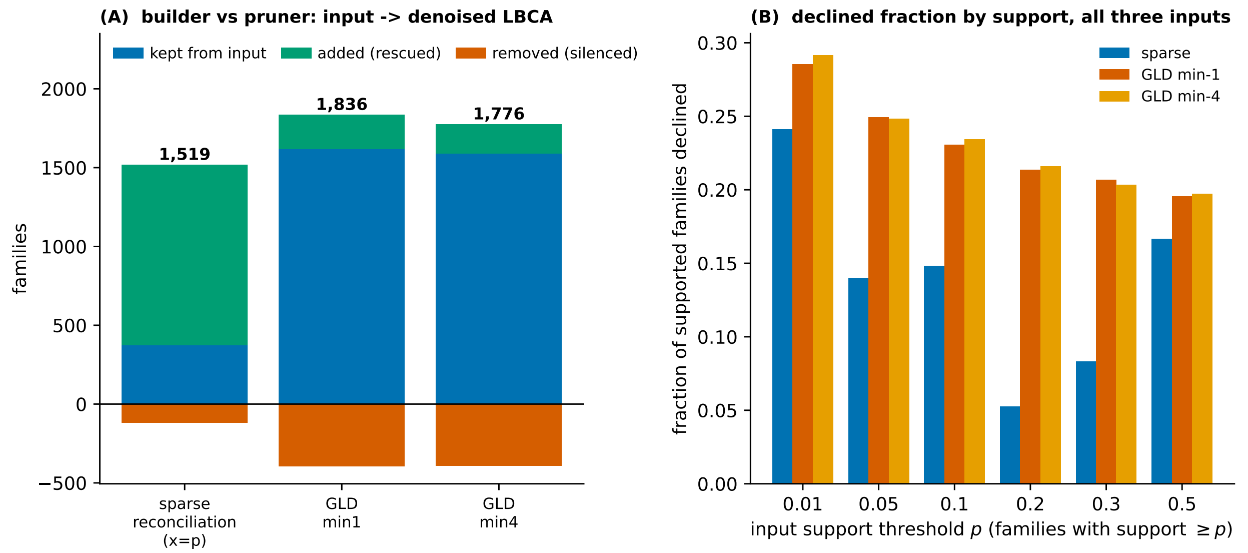


**Figure S13: Builder and pruner.** (A) Each denoised LBCA split into genes kept from its input plus genes added (above the axis) and genes removed (below); the final gene count sits above each bar. The sparse reconciliation is built up, the dense GLDs trimmed down. (B) the *fraction* of each input's supported families (support $\geq p$) that the model declines, grouped over all three LBCA inputs: the sparse reconciliation declines only its weak tail (steeply, 24% at any support to 5% by $\geq 0.2$), while the dense GLD min-1/min-4 prune ∼20% of families *uniformly* across all support.

**The two ancestors meet in the middle.** Opposite operations on opposite inputs land in nearly the same place (Table S12). The dense and sparse LBCAs share ∼1,430 families; measured as a Jaccard index that is 0.75–0.76, i.e. the two reconstructions overlap in about three-quarters of their combined gene set. Each keeps a small remainder the other lacks: ∼400 families unique to the (larger) GLD ancestor, ∼90 unique to the sparse build.

| comparison | shared families | Jaccard | GLD-only | sparse-only |
|---|---|---|---|---|
| min1 vs. sparse | 1,433 | 0.75 | 403 | 86 |
| min4 vs. sparse | 1,426 | 0.76 | 350 | 93 |

**Table S12: The built and pruned LBCAs largely agree.** Overlap of the denoised gene sets (present at posterior $> 0.5$). The two GLD cuts – min1 keeps families seen in $\geq 1$ genome, min4 in $\geq 4$ – give the same answer, so the picture does not hinge on that choice.

**Where they disagree.** The two leftover sets have opposite character. The families **unique to the GLD** ancestor are an accessory/regulatory shell: poorly-characterised "general function" and "unknown" families dominate ($\sim 130$ of them), followed by defence, signalling, and post-translational systems. These are peripheral genes that the GLD reconstruction, prone to inflating root content, over-calls, and they complete few pathways. The families **unique to the sparse build** are instead metabolic core (carbohydrate, amino-acid, energy, lipid, and coenzyme genes) and they finish KEGG metabolic modules the GLD had left fragmentary. Although the sparse build is $\sim 320$ genes *smaller*, it completes 77 full KEGG modules to the dense min1's 81 (77 at min4), nearly as metabolically complete despite the size gap. The substantive disagreement is in the respiratory chain and in how elaborate a chemiosmotic, quinone-linked chain the LBCA carried; neither reconstruction retains a terminal oxidase, as befits an ancestor $\sim 1.5$ Gyr older than the Great Oxidation Event.

#### S8 The five-input LACA convergence

Five reconstructions of the LACA are denoised identically – raw $x = p$ through the cross-input-consistent marginal (mix-FT-fp-marginal-cons, 10-split ensemble), present at posterior mean $> 0.5$. They differ in how the root was inferred and how its gene content is expressed:

- **COG-based uniform-origination**: an older archaeal-root reconciliation – two rootings combined under a uniform gene-origination prior – already expressed in COG space and read directly as per-COG root presence.
- **Euryroot arCOG-aggregate ML-origination**: the updated Euryarchaeota-rooted reconciliation, a maximum-likelihood gene-event reconstruction in native arCOG families.
- **Euryroot arCOG-aggregate uniform-origination**: the *same* Euryarchaeota-rooted arCOG reconciliation, but run *without* origination inference. A deliberately sparse reconciliation (93 families above 0.5) that isolates the effect of the origination model from the tree.
- **arCOG-aggregate recount min-1** and **min-4**: the Count/GLD (Brownian) root reconstruction of the same archaeal data, at gene-observation thresholds $\Omega_{\text{min}}$=1 and 4.

The four arCOG-native inputs are translated to the denoiser's COG vocabulary the *same* way: each arCOG sub-family is mapped to its COG via the arCOG database, and the posterior presences of all arCOGs landing on one COG are combined by **noisy-OR** ($P(\text{COG}) = 1 - \prod_i (1 - p_i)$); the COG-based input is already COG-native and needs no translation.

| LACA input | input $> 0$ | $\geq 0.01$ | $> 0.1$ | $> 0.5$ | denoised |
|---|---|---|---|---|---|
| COG-based uniform-orig. | 832 | 575 | 254 | 64 | 1,023 |
| Euryroot arCOG-agg. ML-orig. | 2,112 | 1,968 | 1,539 | 1,018 | 1,321 |
| Euryroot arCOG-agg. uniform-orig. | 805 | 615 | 311 | 93 | 909 |
| recount min-1 (arCOG-agg., dense) | 2,466 | 2,299 | 2,181 | 2,017 | 1,708 |
| recount min-4 (arCOG-agg., dense) | 2,240 | 1,789 | 1,573 | 1,422 | 1,405 |

**Table S13: The five LACA inputs and their denoised reconstructions** (raw $x = p$, 10-split consistency ensemble, present $> 0.5$). "Input present" is threshold-dependent: the diffuse COG-based posterior collapses from 832 ($p > 0$) to 64 ($p > 0.5$), while the dense Count inputs and the sharp Euryroot core hold up.

| Pearson $r$, *inputs* | COG uorig | Eury ML | Eury unif | min1 | min4 |
|---|---|---|---|---|---|
| COG-based uniform-orig. | 1 | 0.42 | 0.57 | 0.26 | 0.34 |
| Euryroot ML-orig. | | 1 | 0.54 | 0.71 | 0.77 |
| Euryroot uniform-orig. | | | 1 | 0.30 | 0.39 |
| recount min-1 | | | | 1 | 0.81 |
| recount min-4 | | | | | 1 |

**Table S14: Correlation of the five *inputs*, before denoising.** Pearson correlation of the per-COG input posteriors – a *threshold-free* measure, appropriate because the diffuse uniform-origination inputs leave almost nothing above 0.5. On the graded signal the inputs are far more alike than a hard cut implies: the two Euryroot variants – same tree and arCOGs, only the origination model differs – correlate at 0.54, and the two uniform-origination builds at 0.57. Mean pairwise $r = 0.51$.

| Pearson $r$, *reconstructions* | COG uorig | Eury ML | Eury unif | min1 | min4 |
|---|---|---|---|---|---|
| COG-based uniform-orig. | 1 | 0.92 | **0.97** | 0.84 | 0.92 |
| Euryroot ML-orig. | | 1 | 0.89 | 0.92 | **0.97** |
| Euryroot uniform-orig. | | | 1 | 0.79 | 0.88 |
| recount min-1 | | | | 1 | 0.95 |
| recount min-4 | | | | | 1 |

**Table S15: Correlation of the denoised reconstructions** (same Pearson measure). Denoising lifts the mean pairwise correlation to 0.90, from 0.51 across the inputs. The two uniform-origination builds become tightest (**0.97**), as does the ML Euryroot with min-4 (**0.97**).

| Jaccard, *reconstructions* (> 0.5) | COG uorig | Eury ML | Eury unif | min1 | min4 |
|---|---|---|---|---|---|
| COG-based uniform-orig. | 1 | 0.73 | **0.84** | 0.59 | 0.71 |
| Euryroot ML-orig. | | 1 | 0.67 | 0.76 | **0.86** |
| Euryroot uniform-orig. | | | 1 | 0.52 | 0.64 |
| recount min-1 | | | | 1 | 0.81 |
| recount min-4 | | | | | 1 |

**Table S16: Hard overlap of the denoised present sets** (Jaccard, present > 0.5). Mean pairwise 0.71. The two uniform-origination sparse builds converge tightest to *each other* (**0.84**); the ML-origination Euryroot is most dense-concordant (tightest with min-4, **0.86**).

**Convergence.** Denoising pulls the five disparate inputs into close agreement. Measured threshold-free, the mean pairwise Pearson correlation rises from 0.51 across the raw inputs (Table S14) to 0.90 across the reconstructions (Table S15); on the hard present/absent overlap the reconstructions reach Jaccard 0.71 (Table S16). The pull is largest exactly where the inputs disagree most – the COG-based input correlates with recount min-1 at only 0.26 as an input yet 0.84 after denoising (Jaccard 0.59); the ML Euryroot rises 0.71→0.92 against min-1 and 0.77→0.97 against min-4. The new uniform-origination Euryroot is the sharpest test: it correlates only 0.54 with its own ML-origination twin as an input – the origination model alone rescales the root that much – yet the two converge to 0.89, and it lands closest of all to the *other* uniform-origination build, the COG-based one (correlation 0.97, Jaccard 0.84), so the origination prior, not the reconciliation pipeline, is what the denoiser is reconciling. All five reconstructions are decisively archaeal – the 20S proteasome, Cdc6, archaeal primase and SAM synthetase, the V/A-type ATPase, $F_{420}$, and the Wood–Ljungdahl / methanogenesis modules are present in every one (Fig. S14) – and they share an 868-family five-way core.

**Where they still differ: metabolic breadth, not phenotype.** The convergence is not complete, and the residual is interpretable. Take the *conservative* dense reconstruction, recount min-4 – the tightest match to Euryroot (Jaccard 0.86). It still calls ∼ 140 families that Euryroot does not, but these are *not* a second phenotype: only ∼19% are general-function/unknown and all five reconstructions share the diagnostic archaeal systems. Instead ∼ 57% carry a metabolic annotation – energy (C), amino-acid (E), carbohydrate (G), and lipid (I) – so the dense root paints a metabolically *broader* ancestor and the sparse builds a leaner core of the same archaeal cell. The more permissive min-1 cut widens the *same* gap to ∼ 400 families (again about half metabolic), so the over-call is the dense GLD reconstruction's characteristic root-inflation – it grows with the gene-occurrence threshold but stays metabolic in character at either cut, not a disagreement about what the archaeal ancestor was.

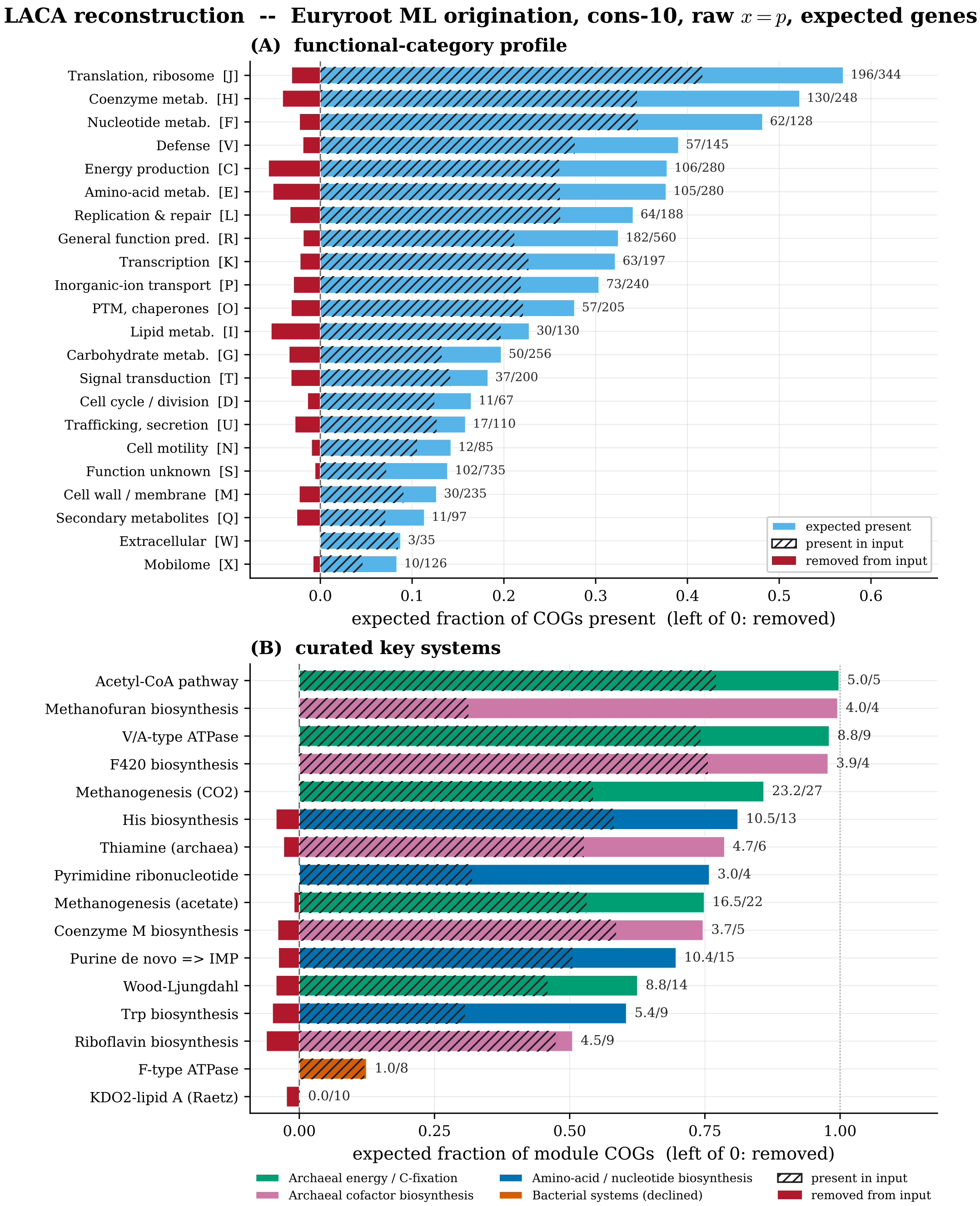


**Figure S14: The focal LACA reconstruction** (updated Euryarchaeota-rooted sparse; arCOG → COG noisy-OR, cons-10 ensemble, raw $x = p$). Bars show the *expected number of genes*: per category/module the posterior probabilities are summed rather than thresholded, so the output is $\mathrm{E}[\#\mathrm{present}] = \sum \mathrm{posterior}$, the hatched input is $\sum \min(\mathrm{input}, \mathrm{output})$ per COG, and removed (left of 0) is $\sum \max(0, \mathrm{input} - \mathrm{output})$. (A) per-COG functional-category profile (solid = expected present at output, hatched = expected present in input, grey = expected removed); (B) curated key systems. The Euryroot input carries $\sim$1,032 expected genes and the denoiser lifts it to $\sim$1,322 (+290 net). Decisively archaeal; the dense GLD and COG-based reconstructions share the same archaeal core.

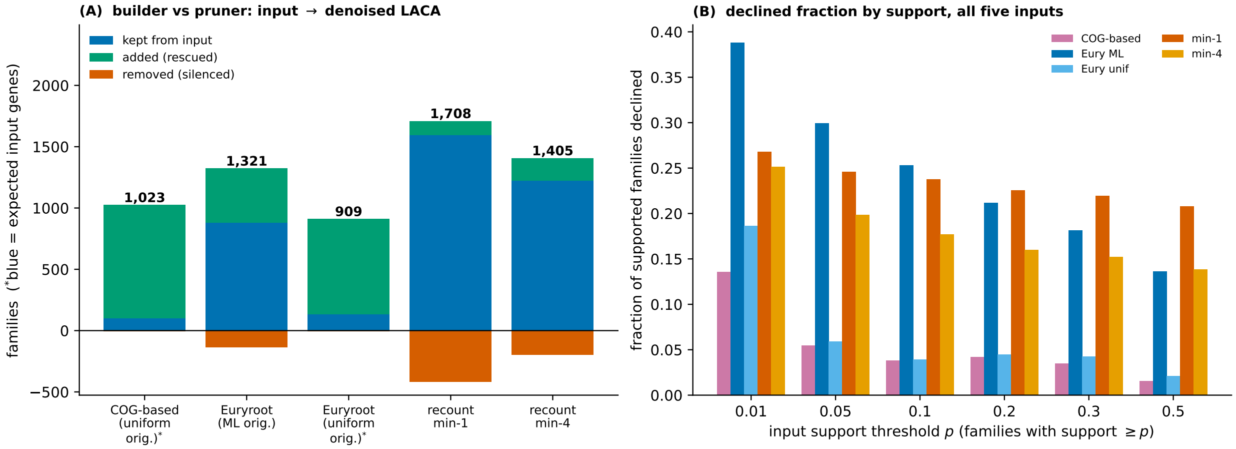


**Figure S15: Builder and pruner (LACA).** (A) each denoised LACA split into kept + added (rescued, above) and removed (silenced, below). For the two diffuse uniform-origination roots (asterisked) the kept (blue) base is the *expected* input contribution – $\sum$(input posterior) over the present COGs – because their $> 0.5$ count badly understates a graded input; the sharp ML and dense roots stay binary ($> 0.5$). Both uniform-origination roots are near-pure builders (COG-based 64→1,023, Euryroot 93→909); recount min-1 is pruned, min-4 about flat. (B) the *fraction* of each input's supported families (support $\geq p$) that the model declines, grouped over all five inputs: the builders decline little, the dense recount roots prune ∼20% across all support, and the ML Euryroot declines its broad weak tail most steeply.

### S9 Per-gene confidence and the shared cellular core

Every COG carries a denoised posterior in $[0, 1]$, read as the mean over ten cross-validation split models and called present above 0.5. Confidence is therefore two-dimensional: the distance of $p$ from the 0.5 boundary, and the cross-split standard deviation of $p$ across the ten models. A family is solid only when both hold: a presence probability far from 0.5 *and* low spread.

**Each count is a confident core plus a soft shell.** By the reconstructed COG presence probability (Table S17), the headline present-counts overstate what is certain. Of the 1,836 families present in the dense LBCA (reconstructed from the GLD/Count min-1 root), 1,177 clear $p > 0.9$ – a confident core – while ∼265 (14%) sit in the borderline band 0.4–0.6, effectively undecided; the sparse build is similar (820 of 1,519). The LACA has the same shape (1,203 of its 1,708 dense, 906 of its 1,321 Euryroot-sparse families confident).

| | reconstruction | present (> 0.5) | confident core ($p > 0.9$) | borderline (0.4–0.6) |
|---|---|---|---|---|
| LBCA | GLD min1 (dense) | 1,836 | 1,177 | 265 (14%) |
| | sparse reconciliation | 1,519 | 820 | 263 (17%) |
| LACA | COG-based uniform-orig. | 1,023 | 636 | 149 (15%) |
| | Euryroot ML-orig. | 1,321 | 906 | 176 (13%) |
| | Euryroot uniform-orig. | 909 | 549 | 157 (17%) |
| | recount min-1 (dense) | 1,708 | 1,203 | 219 (13%) |
| | recount min-4 (dense) | 1,405 | 997 | 161 (11%) |

**Table S17: Each reconstruction is a confident core plus a soft shell** (full ten-split ensemble, all five LACA reconstructions). "confident core" = mean posterior $p > 0.9$; "borderline" = $0.4 < p < 0.6$, essentially undecided calls. The two uniform-origination sparse builds carry the largest borderline shells (15–17%); the dense recount roots the smallest.

**The dense reconstructions are also the least split-stable.** The two axes agree on where the doubt lies. The dense GLD calls carry by far the largest cross-split spread (mean sd 0.084 at the LBCA, with 482 present families exceeding sd 0.15 – their presence flips with the held-out subset), while the sparse builds are the most reproducible (LBCA sparse: 150 such families; the most conservative LACA builds steadier still, mean sd 0.04–0.05 – Table S18). Individual split models put the dense LBCA anywhere from 1,585 to 1,908. The dense over-call is weak on both axes, which is why the ensemble mean rather than any one split model is reported.

| reconstruction | present | mean SD | flip (SD > 0.15) | graded input |
|---|---|---|---|---|
| *LBCA* – bacterial marginal-HQ ensemble | | | | |
| GLD min-1 (dense) | 1,836 | 0.084 | 26.3% | 11% |
| GLD min-4 | 1,776 | 0.084 | 26.3% | 11% |
| sparse reconciliation | 1,519 | 0.062 | 9.8% | 10% |
| *LACA* – cross-input-consistent ensemble | | | | |
| recount min-1 (dense) | 1,708 | 0.071 | 22.1% | 12% |
| recount min-4 (dense) | 1,405 | 0.054 | 14.9% | 19% |
| Euryroot ML-orig. (focal) | 1,321 | 0.055 | 14.1% | 75% |
| Euryroot uniform-orig. | 909 | 0.047 | 6.1% | 54% |
| COG-based uniform-orig. | 1,023 | 0.041 | 3.5% | 43% |

**Table S18: Every reconstruction quantified and ranked by absolute noise.** Mean cross-split SD of the ensemble posterior and the fraction of present calls whose presence flips with the held-out split (SD > 0.15), over each ancestor's ten phylum-holdout split models, with the share of present calls sitting on a graded (non-binary, 0.02–0.98) input bit. Rows are ordered noisiest to cleanest within each ancestor. Absolute SD is comparable *within* an ancestor (one shared ensemble); the LBCA marginal-HQ model and the LACA cross-input-consistent fine-tune of it are different ensembles, so cross-ancestor magnitudes are only indicative. **Split-noise tracks how hard a reconstruction builds**: within each ancestor the dense over-builder is noisiest and the most conservative build cleanest – at the LBCA the GLD roots flip a quarter of their calls (26%) against the sparse reconciliation's 9.8%; at the LACA the dense recount min-1 flips 22% against the under-building COG-based and uniform-origination roots' 3.5–6%.

**The shared core is rock-solid; the differences are not.** The confidence split (the confident core, $p > 0.9$) separates what every reconstruction agrees on from the families one method calls and another does not. At the **LBCA**, intersecting the dense GLD min-1 reconstruction with the sparse reconciliation leaves 1,433 shared families, 779 of them confident in *both* – the part of the LBCA we are sure about whatever the input. The disagreement is one-sided: of the 403 families the dense GLD adds over the sparse build, only 56 (14%) are confident, so the accessory shell is a tentative *over-call*.

At the **LACA** the same split distinguishes a *good* sparse reconstruction from an *under-building* one (Table S19). The ML-origination Euryroot and the conservative dense recount min-4 are the tightest pair (confident-core Jaccard 0.81): min-4 calls only 142 families Euryroot does not, and just 17 (12%) are confident – the same tentative over-reach as the LBCA. The *uniform*-origination Euryroot is the opposite case: it supports only 909 families, and of the 503 that min-4 adds over it, 156 (31%) are confident – against this no-origination input the dense root is not over-calling but confidently recovering genes the reconciliation cannot build. All five LACA reconstructions share a 506-family confident core.

| Jaccard, *confident cores* ($p > 0.9$) | COG uorig | Eury ML | Eury unif | min1 | min4 |
|---|---|---|---|---|---|
| COG-based uniform-orig. | 1 | 0.64 | 0.78 | 0.52 | 0.62 |
| Euryroot ML-orig. | | 1 | 0.58 | 0.70 | **0.81** |
| Euryroot uniform-orig. | | | 1 | 0.45 | 0.54 |
| recount min-1 | | | | 1 | 0.79 |
| recount min-4 | | | | | 1 |

**Table S19: Overlap of the five confident cores** (families called at $p > 0.9$). The high-confidence calls overlap somewhat *less* than the full present sets (Table S16) – each reconstruction is confident about a partly distinct set – but the ML Euryroot and recount min-4 stay tightest (**0.81**). All five share a 506-family confident core.

**The shared cellular core.** Intersecting the LBCA and LACA reconstructions isolates a ∼197-family core present in both domains, dominated by the information-processing machinery shared by all cellular life – ribosomal proteins and assembly factors, aminoacyl-tRNA synthetases, the RNA-polymerase core, translation factors, and replication/repair enzymes – together with conserved central-metabolic and cofactor functions. This core is recovered identically across architectures and ensembles. The domain-specific remainders are equally interpretable: the LACA-only set is enriched for archaeal informational and coenzyme machinery, the LBCA-only set for the flagellum, peptidoglycan biosynthesis, two-component signalling, and fatty-acid synthesis.

**The phenotype read inherits the confidence structure.** The respiratory-chain disagreement sits almost entirely in the LBCA's soft shell. The shared chemiosmotic backbone – F-type ATP synthase, glycolysis, the TCA cycle – is in the double-confident core; the *elaboration* that distinguishes the reconstructions (the dense ancestor's added Complex I and menaquinone completeness and its $bc_1$ fragments; the sparse build's *bd*-type genes) is borderline and split-dependent. The defensible claim is that the LBCA had a complete chemiosmotic core with high confidence, while *which* respiratory elaboration it carried is genuinely uncertain at the gene level – a hypothesis about the uncertain shell, not a firm difference.

### S10 Phenotype prediction under gene loss

Main-text Fig. 5 asks whether the denoiser is *useful* as a preprocessing front end for downstream trait classifiers on degraded genomes. This section gives the protocol, the metrics, and the per-phenotype numbers.

#### S10.1 Three phenotypes, one protocol

We test three trait predictors of A. Koldaeva (gradient-boosted trees over a COG presence/absence vocabulary), spanning two binary traits and one continuous: **aerobicity** (aerobe vs. anaerobe), **cell envelope** (monoderm vs. diderm), and **optimal growth temperature** (OGT, continuous °C). Each is evaluated under the same ten whole-phylum holdout splits used for the denoiser, so no test phylum is seen in the classifier's training set. Denoising uses the full ten-model ensemble, as elsewhere in the paper, so a test genome's phylum is seen by nine of the ten denoiser checkpoints; the comparison therefore isolates the classifier's generalisation, not the denoiser's.

For each split and each held-out genome we corrupt the *entire* 4,789-COG presence vector at a false-negative (gene-loss) rate $f_N$ and a false-positive rate $r_{FP}$, then read the classifier on either (i) the raw corrupted vector or (ii) its Marginal-HQ-fp denoised reconstruction ($x$=$p$, present at posterior mean $> 0.5$), slicing back the classifier's feature columns only after denoising all 4,789 genes. Because the whole vector – out-of-window context included – is corrupted before denoising, every curve converges to the class prior at $f_N$=1 (all genes deleted): the control that certifies the classifier reads no uncorrupted out-of-window context. We sweep $f_N$ on a 0.05 grid (21 values) and $r_{FP} \in \{0, 0.01, 0.05\}$, with **ten independent noise realizations per cell** pooled before scoring; bands are ±1 s.d. across the ten splits.

#### S10.2 Metrics, including a calibration error in degrees

Recovery is Matthews correlation (MCC) for the two binary traits and RMSE (°C) for OGT. Calibration for the binary traits is the 10-bin expected calibration error (ECE).

For OGT we report a *calibration error in degrees*, the regression analogue of ECE. The OGT predictor is a mixture-of-experts: a meso/thermo gate (split at 45°C) with probabilities $p_{\mathrm{lo}}, p_{\mathrm{hi}}$, per-regime mean regressors $\mu_{\mathrm{lo}}, \mu_{\mathrm{hi}}$, and per-regime $q_{10}/q_{90}$ quantile regressors. Its predictive variance follows the law of total variance,

$$\sigma^2 \;=\; p_{\mathrm{lo}}\, s_{\mathrm{lo}}^2 \;+\; p_{\mathrm{hi}}\, s_{\mathrm{hi}}^2 \;+\; p_{\mathrm{lo}}\, p_{\mathrm{hi}}\, (\mu_{\mathrm{lo}} - \mu_{\mathrm{hi}})^2, \qquad s_\bullet = \frac{q_{90}^\bullet - q_{10}^\bullet}{2 \times 1.28}, \tag{S7}$$

where the first two terms are the within-expert spread (from the 80% quantile interval, $z_{0.9}$=1.28) and the last is the gating (meso/thermo) ambiguity. The stated s.d. is $\sqrt{\sigma^2}$. We bin the test genomes into ten equal-count bins by stated s.d. and, in each bin, compare the stated s.d. to the *realized* RMS error, taking the sample-weighted mean $|\,\text{stated} - \text{actual}\,|$. Zero means the stated uncertainty tracks the realized error magnitude; a large value means systematic over- or under-confidence. Panel (f) of main-text Fig. 5 caps this at 6°C: it stays in the 1.5–3.5°C range across the informative regime and only diverges at the $f_N \to 1$ prior limit, where the model is asked to describe a genome that no longer exists. (The saved OGT robust models shipped only three of the ten splits, so all ten were refit inline from `ogt_cog.csv` with the identical noise-augmented recipe – validated by recovering the canonical 4,789-COG set at Jaccard 0.99; see the reproduction notes.)

##### S10.3 Result: rescue, and the OGT crossover

At $r_{FP}$=0.01 (Table S20), denoising leaves clean genomes essentially untouched and sharply rescues the binary traits under loss: aerobicity MCC 0.35 → 0.80 and cell-envelope MCC 0.49 → 0.92 at $f_N$=0.9, where the raw input has collapsed. OGT behaves differently: denoising a clean genome *costs* accuracy (RMSE 3.4 → 6.8°C, the price of hallucinating a few genes onto an already-complete input), the two arms cross near $f_N \approx 0.8$, and only under heavy loss does denoising win (RMSE 11.7 → 8.7°C at $f_N$=0.9). Its calibration error improves under heavy loss (4.0 → 3.1°C at $f_N$=0.9), but is slightly worse across the mid-range ($f_N \approx 0.2$–0.65, by up to 0.7°C). The asymmetry is expected: aerobicity and cell envelope hinge on the presence of a handful of marker systems the denoiser restores, whereas OGT reads a diffuse genome-wide signal that a near-complete input already carries, so denoising helps only once enough of that signal has been destroyed.

| metric ($r_{FP}$=0.01, noisy→denoised) | $f_N$=0 | 0.75 | 0.9 | 0.95 |
|---|---|---|---|---|
| Aerobicity MCC ↑ | 0.85 → 0.84 | 0.64 → 0.85 | 0.35 → 0.80 | 0.20 → 0.57 |
| Mono/diderm MCC ↑ | 0.94 → 0.95 | 0.74 → 0.96 | 0.49 → 0.92 | 0.28 → 0.72 |
| OGT RMSE (°C) ↓ | 3.4 → 6.8 | 7.5 → 8.0 | 11.7 → 8.7 | 14.6 → 10.0 |
| OGT calib. err. (°C) ↓ | 1.7 → 1.5 | 2.9 → 2.5 | 4.0 → 3.1 | 3.4 → 3.4 |

**Table S20: Denoising rescue by phenotype**, noise-robust predictor, $r_{FP}$=0.01, mean over the ten whole-phylum splits (main-text Fig. 5). Binary traits are rescued sharply under loss with no clean-genome cost; OGT trades clean accuracy for robustness, the arms crossing near $f_N \approx 0.8$.

#### S11 Data and code availability

The source code for the denoiser, together with the reconstruction command-line interface and the scripts that regenerate the tables and figures reported here, is at `https://github.com/ssolo/gene-content-grammar`.

The trained model checkpoints, the training and validation data, and the analysis pipeline are archived on Zenodo under the concept DOI `10.5281/zenodo.20526264`, which resolves to the latest version. That record holds the ten cross-validation checkpoints for each production model – the generalist denoiser, the bacterial specialist used for the LBCA, and the cross-input-consistency fine-tune used for the LACA – together with the whole-phylum training and validation splits and the COG vocabulary.

The per-COG reconstruction tables for the LBCA and the LACA are released with the code. Each records, for every one of the 4,789 COG families, the input value the model was given, the ensemble posterior averaged over the ten cross-validation models, and the standard deviation across those models, under each of the reconciliation- and profile-based input encodings discussed above. The per-COG *E. coli* recovery table underlying the extant-genome test and the reconstruction-agreement tables are released alongside them.

The gain-loss-duplication reconstructions used as profile-level input were produced with `recount` (Szöllősi & Williams, 2026), which is archived separately.

Code and data are made available under a Creative Commons Attribution–NonCommercial 4.0 licence. Data derived from the GTDB retain that database's own terms, and no KEGG-derived content is redistributed.
